# Genome-wide association study of diabetic kidney disease highlights biology involved in renal basement membrane collagen

**DOI:** 10.1101/499616

**Authors:** Rany M. Salem, Jennifer N. Todd, Niina Sandholm, Joanne B. Cole, Wei-Min Chen, Darrel Andrews, Marcus G. Pezzolesi, Paul M. McKeigue, Linda T. Hiraki, Chengxiang Qiu, Viji Nair, Chen Di Liao, Jing Jing Cao, Erkka Valo, Suna Onengut-Gumuscu, Adam M. Smiles, Stuart J. McGurnaghan, Jani K. Haukka, Valma Harjutsalo, Eoin P. Brennan, Natalie van Zuydam, Emma Ahlqvist, Ross Doyle, Tarunveer S. Ahluwalia, Maria Lajer, Maria F. Hughes, Jihwan Park, Jan Skupien, Athina Spiliopoulou, Andrew Liu, Rajasree Menon, Carine M. Boustany-Kari, Hyun M. Kang, Robert G. Nelson, Ronald Klein, Barbara E. Klein, Kristine E. Lee, Xiaoyu Gao, Michael Mauer, Silvia Maeastroni, Maria Luiza Caramori, Ian H. de Boer, Rachel G. Miller, Jingchuan Guo, Andrew P. Boright, David Tregouet, Beata Gyorgy, Janet K. Snell-Bergeon, David M. Maahs, Shelley B. Bull, Angelo J. Canty, Colin N.A. Palmer, Lars Stechemesser, Bernhard Paulweber, Raimund Weitgasser, Jelizaveta Sokolovska, Vita Rovīte, Valdis Pīrāgs, Edita Prakapiene, Lina Radzeviciene, Rasa Verkauskiene, Nicolae Mircea Panduru, Leif C. Groop, Mark I. McCarthy, Harvest F. Gu, Anna Möllsten, Henrik Falhammar, Kerstin Brismar, GENIE Consortium, DCCT/EDIC Research Group, SUMMIT Consortium, Finian Martin, Peter Rossing, Tina Costacou, Gianpaolo Zerbini, Michel Marre, Samy Hadjadj, Amy J. McKnight, Carol Forsblom, Gareth McKay, Catherine Godson, A. Peter Maxwell, Matthias Kretzler, Katalin Susztak, Helen M. Colhoun, Andrzej Krolewski, Andrew D. Paterson, Per-Henrik Groop, Stephen S. Rich, Joel N. Hirschhorn, Jose C. Florez

## Abstract

Diabetic kidney disease (DKD) is a heritable but poorly understood complication of diabetes. To identify genetic variants predisposing to DKD, we performed genome-wide association analyses in 19,406 individuals with type 1 diabetes (T1D) using a spectrum of DKD definitions basedon albuminuria and renal function. We identified 16 genome-wide significant loci. The variant with the strongest association (rs55703767) is a common missense mutation in the collagen type IV alpha 3 chain *(COL4A3)* gene, which encodes a major structural component of the glomerular basement membrane (GBM) implicated in heritable nephropathies. The rs55703767 minor allele (Asp326Tyr) is protective against several definitions of DKD, including albuminuria and end-stage renal disease. Three other loci are in or near genes with known or suggestive involvement in DKD *(BMP7)* or renal biology (*COLEC11* and *DDR1*). The 16 DKD-associated loci provide novel insights into the pathogenesis of DKD, identifying potential biological targets for prevention and treatment.

---

The devastating diabetic complication of DKD is the major cause of end-stage renal disease (ESRD) worldwide^1,2^. Current treatment strategies at best slow the progression of DKD, and do not halt or reverse the disease. Although improved glycemic control influences the rate of diabetic complications, a large portion of the variation in DKD susceptibility remains unexplained: one third of people with T1D develop DKD despite adequate glycemic control, while others maintain normal renal function despite long-term severe chronic hyperglycemia^3^.

Though DKD demonstrates both familial clustering^4–6^ and single nucleotide polymorphism (SNP) heritability^7^, the specific genetic factors influencing DKD risk remain largely unknown. Recent genome-wide association studies (GWAS) have only identified a handful of loci for DKD, albuminuria, or estimated glomerular filtration rate (eGFR) in individuals with diabetes^7–13^. Potential reasons for the limited success include small sample sizes, modest genetic effects, and lack of consistency of phenotype definitions and statistical analyses across studies.

Through collaboration within the JDRF Diabetes Nephropathy Collaborative Research Initiative (DNCRI), we adopted three approaches to improve our ability to find new genetic risk factors for DKD: 1) assembling a large collection of T1D cohorts with harmonized DKD phenotypes, 2) creating a comprehensive set of detailed DKD definitions, and 3) augmenting genotype data with low frequency and exome array variants.

## RESULTS

### Phenotypic comparisons

We investigated a broad spectrum of DKD definitions based on albuminuria and renal function criteria, defining a total of 10 different case-control comparisons to cover the different aspects of disease progression (**Figure 1**). Seven comparisons were based on albuminuria and/or ESRD (including diabetic nephropathy [DN], defined as either macroalbuminuria or ESRD); two were defined based on eGFR (used to classify severity of chronic kidney disease [CKD]); and one combined both albuminuria and eGFR data (“CKD-DN”). Each phenotypic definition was analyzed separately in GWAS; to account for the 10 definitions each analyzed under two covariate adjustment models, we estimated^14^ the total effectively independent tests as 7.4, allowing us to compute a more conservative study-wide significance threshold (*P*<6.76×10^-9^), based on genome-wide significance (*P*<5×10^-8^) and Bonferroni correction for 7.4 effective tests.

**Figure 1.**
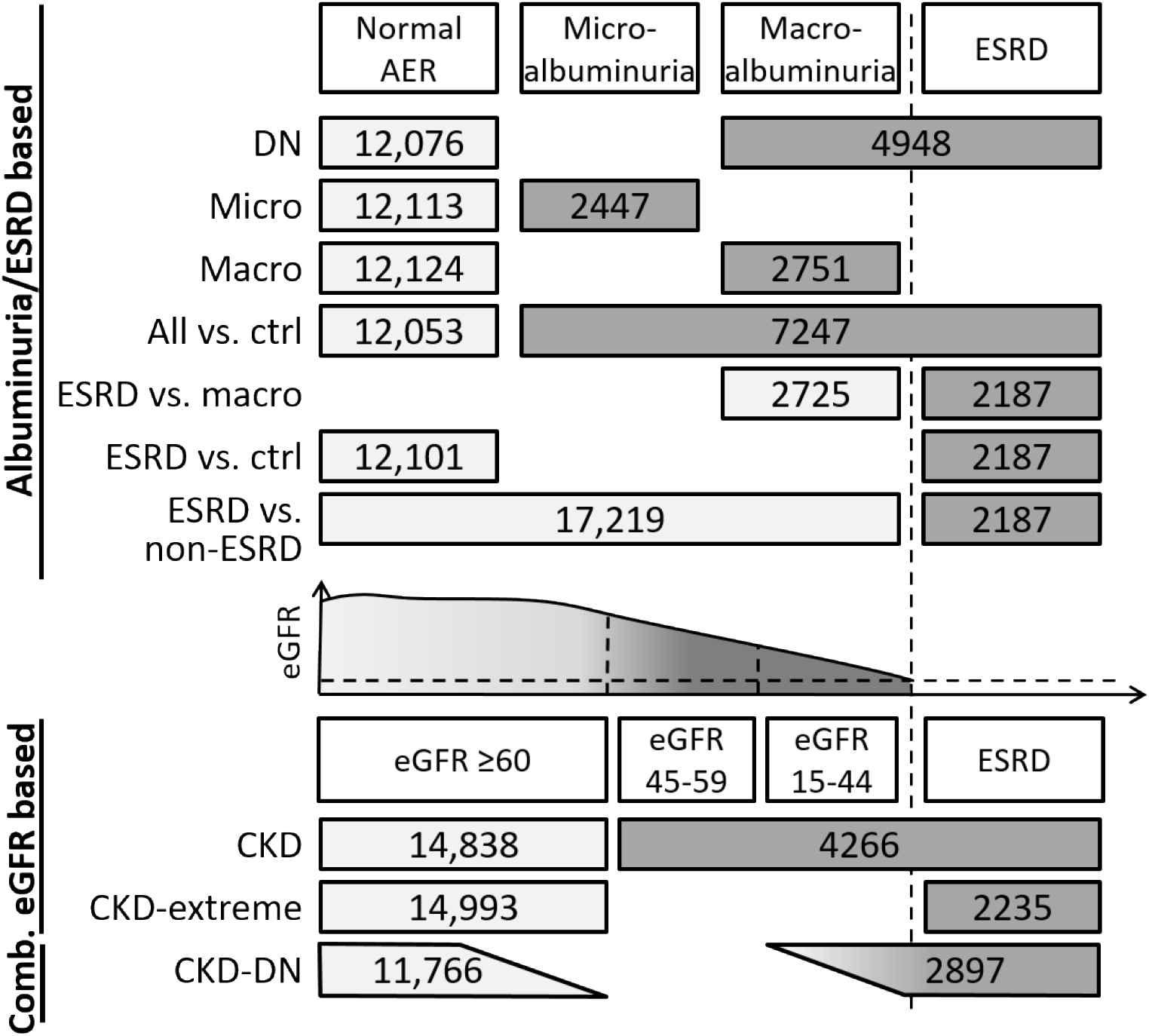
Phenotypic analysis of DKD. Schematic diagram of outcomes analyzed in this study. Numbers indicate the total number of cases (darker gray) and controls (lighter gray) included in the meta-analyses for each phenotype. Microalb.: microalbuminuria; macroalb.: macroalbuminuria; eGFR: estimated glomerular filtration rate; ESRD: End-stage renal disease, defined as eGFR <15 mL/min/1.73m^2^ or undergoing dialysis or having renal transplant; CKD: chronic kidney disease.

### Top genome-wide association results highlight *COL4A3*

GWAS meta-analysis included association results for up to 19,406 individuals with T1D of European descent from 17 cohorts for the 10 case-control definitions (**Table S1**). We identified 16 novel independent loci that achieved genome-wide significance (*P*<5×10^-8^), in which four lead SNPs also surpassed our more conservative study-wide significance threshold (**Table 1; Figure 2**, Manhattan plot; **Figures S1a-p**, regional association plots). The strongest signal was rs55703767 (minor allele frequency [MAF]=0.21), a common missense variant (G>T; Asp326Tyr) in exon 17 of *COL4A3.* This SNP was associated with protection from DN (odds ratio [OR]=0.79, *P*=5.34×10^-12^), any albuminuria (OR=0.84, *P*=3.88×10^-10^), the combined CKD-DN phenotype (OR =0.77, *P*=5.30×10^-9^), and macroalbuminuria (OR=0.79, *P*=9.28×10^-9^). Interestingly, we found that rs55703767 in *COL4A3* was more strongly associated in men (OR=0.73, *P*=1.29×10^-11^) than in women (OR=0.85, *P*=1.39×10^-3^; Phet=1.58×10^-2^). *COL4A3* encodes the alpha 3 chain of collagen type IV, a major structural component of the GBM^15^.

**Figure 2.**
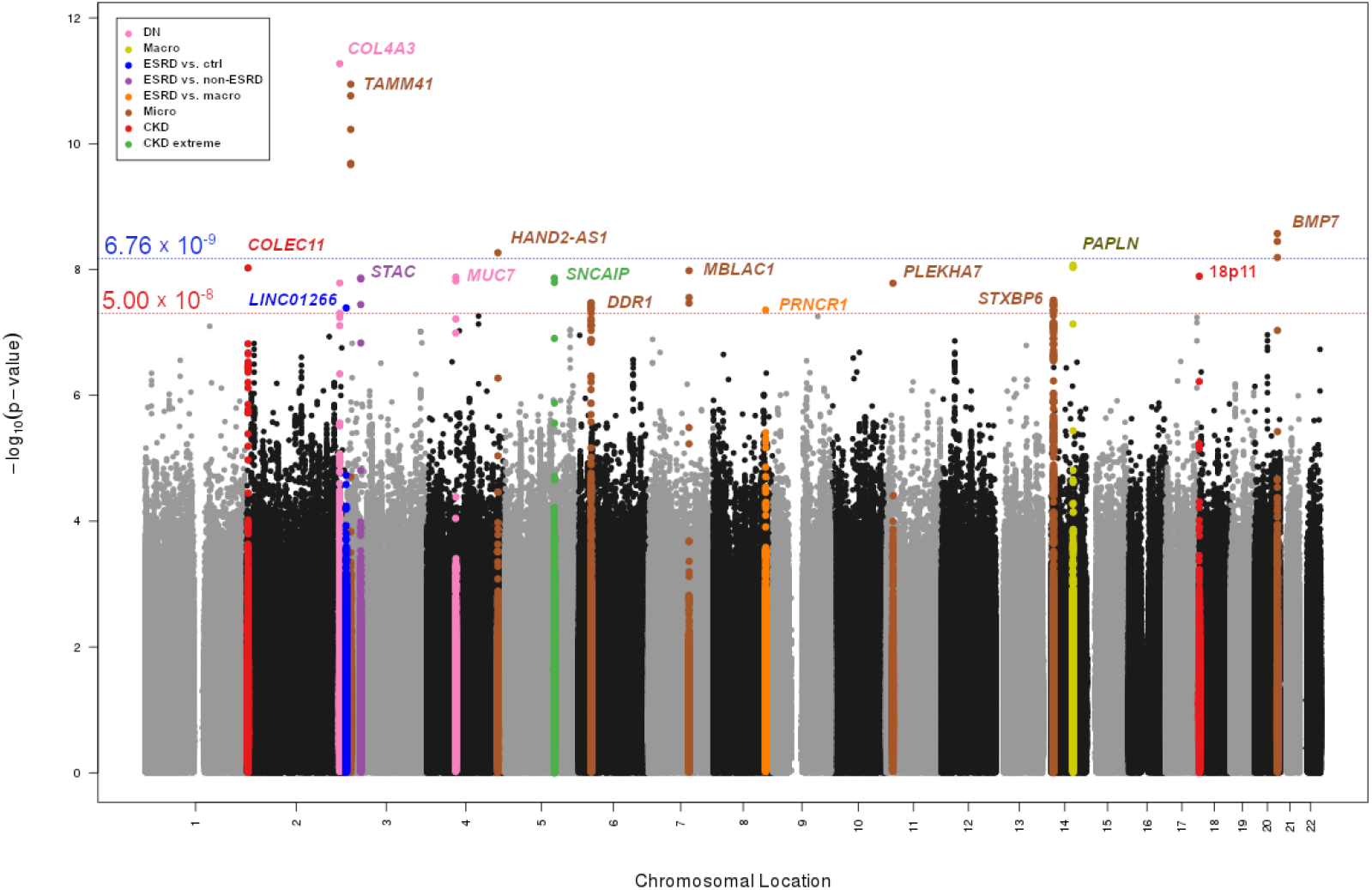
Genome-wide association testing of all 10 phenotypic comparisons. Multiphenotype Manhattan plot shows lowest P-value at each marker for each of the 10 phenotypic comparisons, under the standard and fully-adjusted model. Significance of SNPs (-log_10_[P-value], y axis) is plotted against genomic location (x axis). Loci surpassing genome-wide significance (red line) and/or study-wide significance (blue line) are colored by phenotype.

**Table 1.**
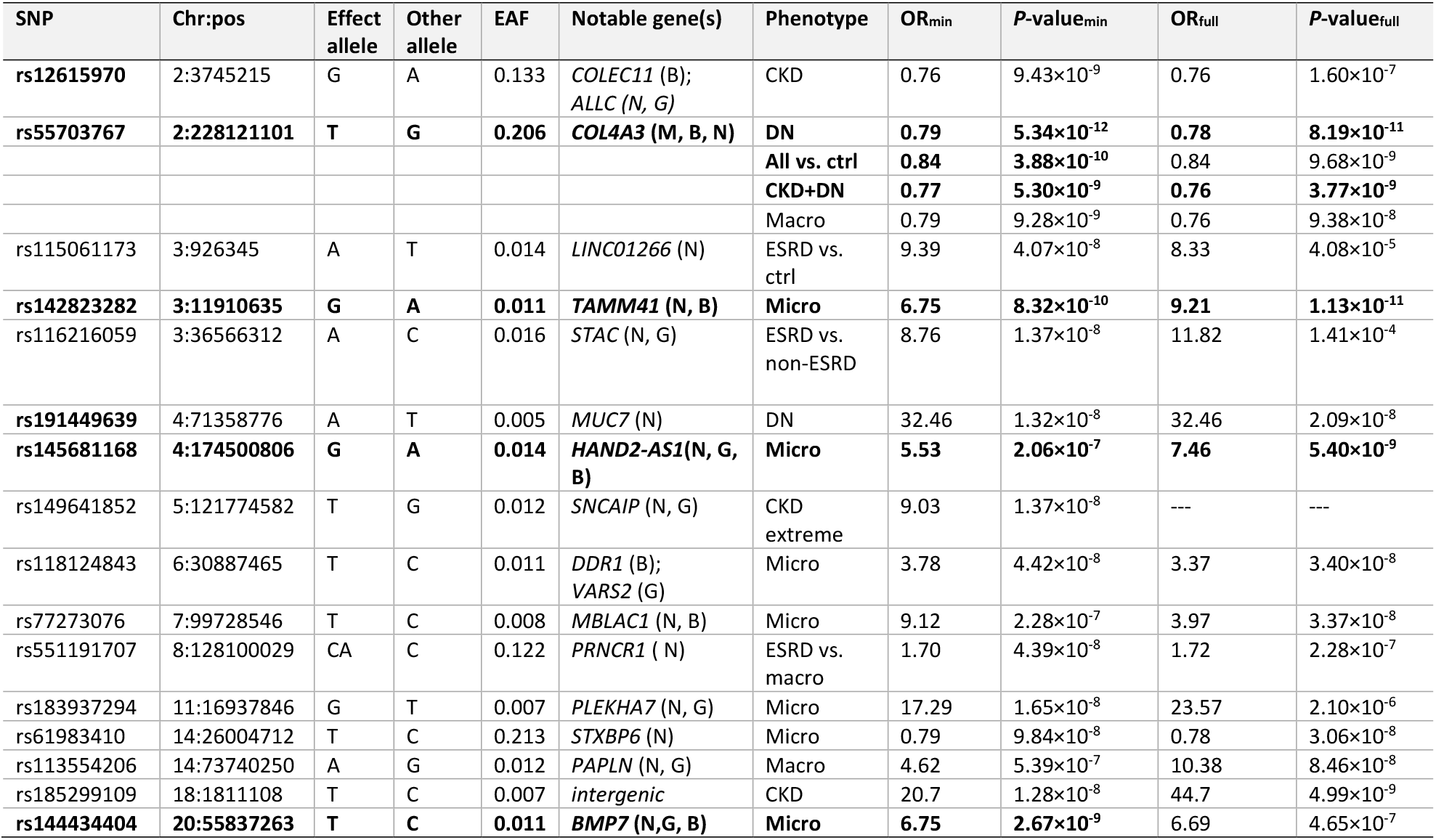
Loci associated with DKD at study-wide (*P*<6.76×10^-9^, bold) and genome-wide (*P*<5×10^-8^) significance. Notable genes from missense variant in the indicated gene (M); intronic, synonymous, or noncoding transcript variant within gene (G); gene nearest to index SNP (N); biological relevance to kidney biology (B). Chr, chromosome; pos, position; EAF, effect allele frequency; OR, odds ratio; min, minimally adjusted covariate model; full, fully adjusted covariate model.

### *COL4A3* variation and kidney phenotypes

In persons with T1D and normoalbuminuria, GBM width predicts progression to proteinuria and ESRD independently of glycated hemoglobin (HbA1c)^16^. We examined the influence of the *COL4A3* variant on GBM width in 253 Renin-Angiotensin System Study (RASS)^17^ participants with T1D who had biopsy and genetic data (**Table S2**). The DKD-protective minor T allele was associated with 19.7 nm lower GBM width (standard error (SE) 8.2 nm, P=0.02), with the lowest mean GBM width among TT homozygotes (**Figure 3; Table S3**), after adjusting for age, sex, and diabetes duration, and without detectable interactions with T1D duration or mean HbA1c. Furthermore, in a Pima Indian cohort of 97 subjects with DKD with morphometric and expression data from renal biopsies, *COL4A3* expression was negatively correlated with the GBM surface density (filtration surface density) (β=-0.27, P=0.02), which is associated with eGFR in DKD in both T1D and type 2 diabetes (T2D)^18,19^.

**Figure 3.**
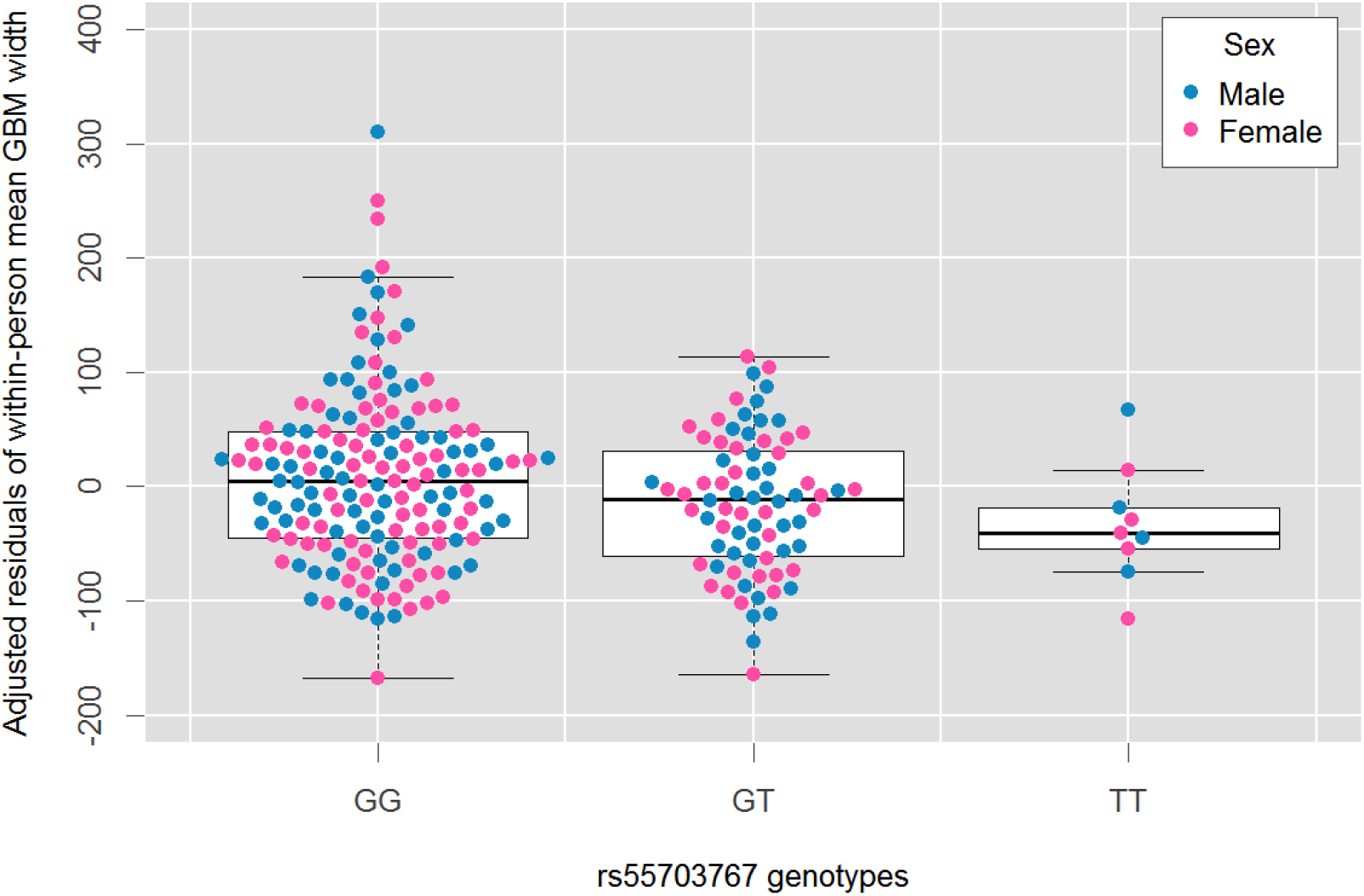
Adjusted residuals of GBM width by rs55703767 genotype and sex. Box and whisker plot of residuals of mean GBM width after adjusting for age, sex, and diabetes duration, stratified by GG, GT, or TT genotype at rs55703767, with overlay of individual data points for both females (pink) and males (blue).

To establish whether expression of *COL4A3* was further correlated with DKD phenotypes, we examined micro-dissected human kidney samples from 455 subjects (433 tubule samples and 335 glomerulus samples) with pathological and RNAseq data. Expression of *COL4A3* was positively correlated with the degree of fibrosis in tubules (corr=0.289, *P*=3.2×10^-9^) but negatively correlated with glomerulosclerosis (corr=0.16, *P*=4.8×10^-3^; **Figure S2**). *COL4A3* expression in glomeruli, but not in tubules, was also nominally correlated with eGFR (corr 0.108, *P*=0.047; **Figure S2**).

### Evidence for hyperglycemia specificity

Hyperglylcemia promotes the development of diabetic complications. If a genetic variant exerts a stronger effect in the setting of hyperglycemia, 1) it might not be detected in general CKD, 2) it may be detected whether hyperglycemia is conferred by T1D or T2D, 3) its effect may be stronger at higher glycemic strata, and 4) interventions that reduce glycemia may attenuate the association signal. *COL4A3* rs55703767 was not associated with eGFR in a general population sample of 110,517 mainly non-diabetic participants of European ancestry^20^ (**Table S4**). However, in a smaller cohort of 5,190 participants with T2D and DKD phenotypes in the SUrrogate markers for Micro- and Macro-vascular hard endpoints for Innovative diabetes Tools (SUMMIT) consortium, we detected a directionally consistent suggestive association of *COL4A3* rs55703767 with DN (2-tailed *P*=0.08; **Table S5**).

We further stratified the association analyses by HbA1c in the Finnish Diabetic Nephropathy (FinnDiane) Study, a cohort study with extensive longitudinal phenotypic data^21^. Based on the time-weighted mean of all available HbA1c measurements for each individual, 1,344 individuals had mean HbA1c <7.5% (58 mmol/mol), and 2,977 with mean HbA1c ≥7.5%. *COL4A3* rs55703767 was nominally significant (*P*<0.05) only in individuals with HbA1c ≥7.5% (**Figure 4, Table S6, Figure S3**). However, in a similar study setting of individuals with T2D from the GoDARTS study (N=3226)^13,22^, no difference was observed for *COL4A3* rs55703767 between HbA1c strata (**Figure S4**).

**Figure 4.**
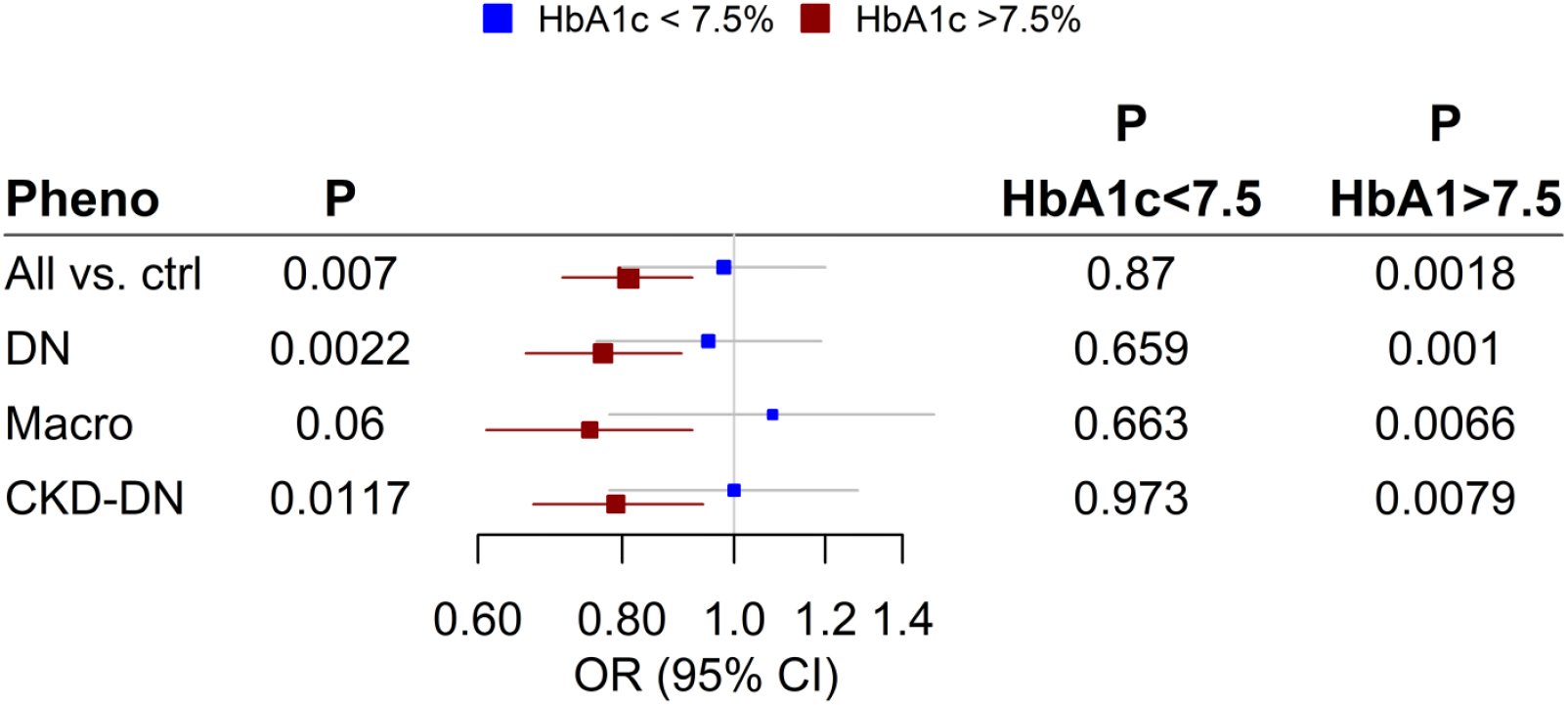
Association at rs55703767 *(COL4A3)* stratified by HbA1c below or above 7.5%, for the phenotypes reaching genome-wide significance in the combined meta-analysis. Analysis included 1344 individuals with time-weighted mean HbA1c <7.5% (58 mmol/mol), and 2977 with mean HbA1c ≥7.5% from the FinnDiane study; the individuals had median 19 HbA1c measurements (range 1 – 129).

We performed a similar HbA1c stratified analysis in the Diabetes Control and Complications Trial (DCCT), whose participants continue to be followed in the Epidemiology of Diabetes Interventions and Complications (EDIC)^23,24^. In DCCT-EDIC the effect of *COL4A3* rs55703767 was stronger among those recruited in the secondary cohort (mild retinopathy and longer diabetes duration at baseline) who were originally randomized to conventional treatment and therefore had higher HbA1c than the intensive treatment group (**Table S7**). Taken together, these independent lines of evidence strongly suggest that the *COL4A3* variant effect on DKD risk is amplified by poor glycemic control.

### Other association signals

Two other genome-wide significant signals were near genes encoding proteins related to collagen. Variant rs12615970 (MAF=0.13), located 53 kb downstream of *COLEC11*, was associated with CKD (OR=1.31, *P*=9.43×10^-9^), and rs116772905 (MAF=0.011) in exon 14 of *DDR1* was associated with microalbuminuria (OR=3.78, *P*=4.40×10^-8^). rs116772905 is in perfect linkage disequilibrium with rs118124843, the lead association with microalbuminuria for this locus under the full adjustment model (taking into account both BMI and HbA1c), located 29 kb downstream of *DDR1* (OR=3.97, *P*=3.37×10^-8^). *COLEC11* encodes a collectin protein containing both a collagen-like domain and a carbohydrate recognition domain for binding sugars, and *DDR1* encodes the discoidin domain-containing receptor 1, which binds collagens including type IV collagen.

In addition to *COL4A3* rs55703767, three other low-frequency variants associated with microalbuminuria achieved study-wide significance: rs142823282 (MAF=0.017), 22 kb upstream of *TAMM41* encoding a mitochondrial translocator assembly and maintenance protein^25,26^ (OR=6.75, *P*=1.13×10^-11^), rs144434404 (MAF=0.011), in intron 1 of *BMP7* encoding the bone morphogenetic protein 7 previously implicated in DKD^27^ (OR=6.75, *P*=2.67×10^-9^), and rs145681168 (MAF=0.014), in intron 3 of two transcripts of *HAND2* antisense RNA 1 *(HAND2-AS1;* OR=5.53, *P*=5.40×10^-9^) and 50 kb upstream of *HAND2*, encoding a heart and neural crest derivatives transcription factor.

Two additional common variants achieved genome-wide significance: rs551191707 (MAF=0.122) in *PRNCR1* associated with ESRD when compared with macroalbuminuria (OR=1.70, *P*=4.39×10^-8^) and rs61983410 (MAF=0.213) in an intergenic region on chromosome 14 associated with microalbuminuria (full model OR=0.78, *P*=3.06×10^-8^). The remaining eight variants associated with features of DKD had lower allele frequencies (four with 0.01≤MAF≤0.05 and four with MAF<0.01) and did not achieve study-wide significance.

As we had done for *COL4A3* rs55703767, we tested whether the associations of the 15 other variants were amplified by hyperglycemia. None of the 15 variants were significantly associated with eGFR in the general population (**Table S4**). In the smaller SUMMIT T2D cohort^13^ we were able to interrogate seven loci with comparable trait definitions. The odds ratios were directionally consistent in six of them (binomial sign test: *P*=0.0625, **Table S5**). In FinnDiane seven of the remaining 15 loci were observed with sufficient frequency (minor allele counts >10) to allow subgroup analysis. Two additional SNPs (rs149641852 in *SNCAIP* and rs12615970 near *COLEC11)* were nominally significant (*P*<0.05) only in individuals with HbA1c ≥7.5% (**Table S6, Figure S3**).

### Variants previously associated with DKD

We investigated the effect of variants previously associated at genome-wide significance with renal complications in individuals with diabetes^8–13,28^. Across the ten sub-phenotypes in our meta-analysis, we found evidence of association for seven of nine examined loci (*P*<0.05, Figure S5): We replicated two loci that were previously discovered without overlapping individuals with the current study: *SCAF8/CNKSR3* rs12523822, originally associated with DKD (*P*=6.8×10^-4^ for “All vs ctrl”)^8^; and *UMOD* rs77924615, originally associated with eGFR in both individuals with and without diabetes (*P*=5.2×10^-4^ for “CKD”)^20^. Associations at the *AFF3, RGMA-MCTP2*, and *ERBB4* loci, identified in the GEnetics of Nephropathy—an International Effort (GENIE) consortium^12^, comprised of a subset of studies included in this current effort, remained associated with DKD, though the associations were attenuated in this larger dataset *(RGMA-MCTP2* rs12437854 *P*=2.97×10^-5^; *AFF3* rs7583877 *P*=5.97×10^-4^; *ERBB4* rs7588550 *P*=3.53×10^-5^; **Figure S6**). Associations were also observed at the *CDCA7/SP3* (rs4972593, *P*=0.020 for “CKD-DN”, originally for ESRD exclusively in women^11^) and *GLRA3* (rs1564939, *P*=0.016 for “CKD extremes”, originally for AER^10,28^), but these analyses also include individuals that overlap with the original studies. Apart from the *UMOD* locus, none of the 63 loci associated with eGFR in the general population^20^ were associated with DKD after correction for multiple testing (*P*<7.0×10^-4^, **Figure S7**).

### Gene and gene set analysis

We conducted gene-level analyses by employing two methods that aggregate SNP summary statistics over a gene region while accounting for linkage disequilibrium, MAGMA and PASCAL^29,30^. MAGMA identified five genes at a Bonferroni-corrected threshold (*P*<0.05/18,222 genes tested = 2.74×10^-6^): the collagen gene *COL20A1* associated with “CKD extreme” (full model *P*=5.77×10^-7^) and “ESRD vs. non-ESRD” (full model *P*=9.53×10^-7^), *SLC46A2* associated with “All vs. ctrl” (*P*=7.38×10^-7^), *SFXN4* associated with “Macro” (full model *P*=1.65×10^-7^), *GLT6D1* associated with “ESRD vs. macro” (*P*=1.49×10^-6^), and *SNX30* associated with “All vs ctrl” (*P*=2.49×10^-6^) (**Table S8**). Although PASCAL did not identify any significant gene level associations, the five MAGMA-identified genes had *P*<5.0×10^-4^ in PASCAL (**Table S9**). Both *SFXN4* and *CBX8* have been reported to be differentially methylated in patients with diabetes with and without nephropathy^31,32^.

Additionally, we used MAGMA, PASCAL, DEPICT, and MAGENTA to conduct gene-set analysis in our GWAS dataset. The four methods identified 12 significantly enriched gene sets (**Table S10**). One gene set, “negative regulators of RIG-I MDA5 signaling” was identified in two different pathway analyses (MAGMA and PASCAL) of our fully adjusted GWAS of ESRD *vs.* Macro. Several additional related and overlapping gene sets were identified, including “RIGI MDA5 mediated induction of IFN alpha beta pathways”, “TRAF3 dependent IRF activation pathway”, and “TRAF6 mediated IRF activation” (PASCAL) and “activated TLR4 signaling” (MAGENTA). RIG-I, MDA5 and the toll-like receptor TLR4 are members of the innate immune response system that respond to both cellular injury and infection^33,34^ and transduce highly intertwined signaling cascades. These include the signaling molecules TRAF3 and TRAF6, which induce expression of type I interferons and proinflammatory cytokines implicated in the progression of DKD^35,36^. Specifically, the TLR4 receptor and several of its ligands and downstream cytokines display differential levels of expression in DKD renal tubules *vs.* normal kidneys and *vs.* non-diabetic kidney disease controls^37^, and TLR4 knockout mice are protected from DKD and display marked reductions in interstitial collagen deposition in the kidney^38^. Other pathways of interest include “other lipid, fatty acid and steroid metabolism”, “nitric oxide signaling in the cardiovascular system”, and “Tumor necrosis factor (TNF) family member”, with both nitric oxide and TNF-α implicated in DKD^39,40,41^.

### Expression and epigenetic analyses

We interrogated gene expression datasets in relevant tissues to determine whether our top signals underlie expression quantitative trait loci (eQTL). We first analyzed genotype and RNAseq gene expression data from 96 whole human kidney cortical samples^42^ and microdissected human kidney samples (121 tubule and 119 glomerular samples^43^ from subjects of European descent without any evidence of renal disease (**Figure S8**). No findings in this data set achieved significance after correction for multiple testing. In the GTEx and eQTLgen datasets, *COL4A3* rs55703767 had a significant eQTL (*P*=5.63×10^-38^) with the *MFF* gene in blood, and rs118124843 near *DDR1* and *VARS2* had multiple significant eQTLs in blood besides *VARS2* (*P*=1.71×10^-5^; **Table S11**). Interestingly, rs142823282 near *TAMM41* was a *cis-* eQTL for *PPARG* (*P*=4.60×10^-7^), a transcription factor regulating adipocyte development, glucose and lipid metabolism; PPAR_Y_ agonists have been suggested to prevent DKD^44^.

To ascertain the potential functional role of our top non-coding signals, we mined ChIP-seq data derived from healthy adult human kidney samples^45^. SNP rs142823282 near *TAMM41* showed enrichment for the histone marks H3K27ac, H3K9ac, H3K4me1, and H3K4me3, suggesting that this is an active regulator of *TAMM41* or another nearby gene (**Figure S9**). Interestingly, in recent work we have shown that DNA methylation profiles in participants with T1D with/without kidney disease show the greatest differences in methylation sites near TAMM41^46^.

To establish whether the expression of our top genes shows enrichment in a specific kidney cell type, we queried an expression dataset of ~50,000 single cells obtained from mouse kidneys^47^. Expression was detected for six genes in the mouse kidney atlas: three *(COL4A3, SNCAIP*, and *BMP7)* were almost exclusively expressed in podocytes (Figure 5), supporting the significant role for podocytes in DKD.

**Figure 5.**
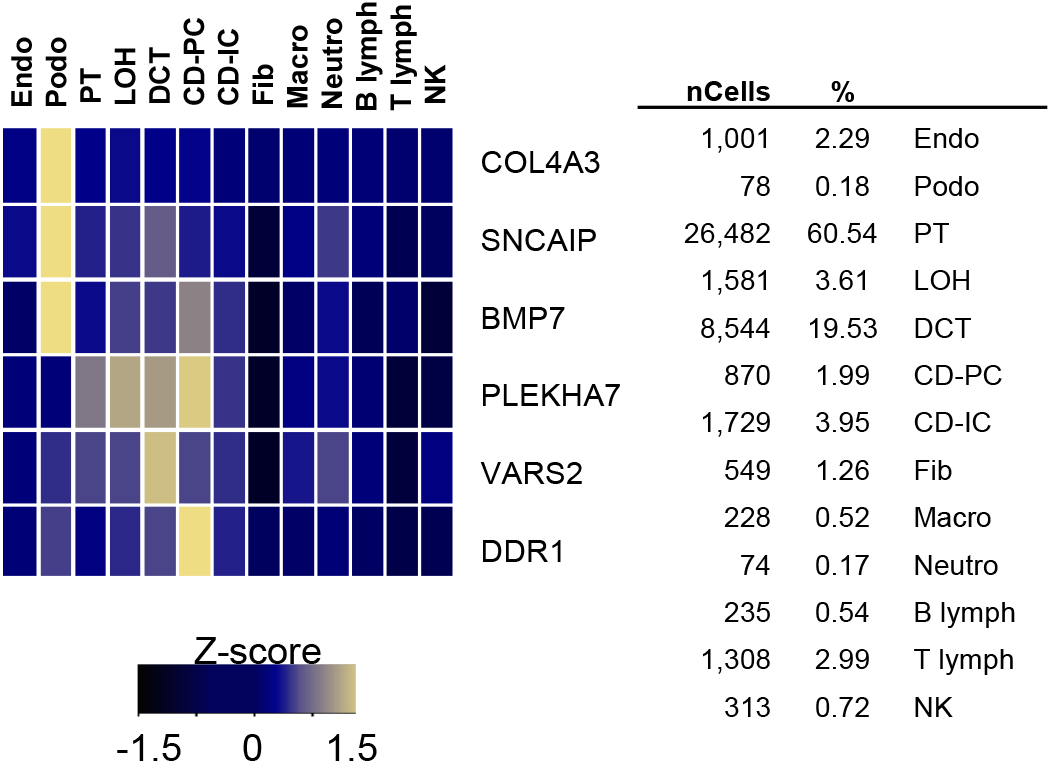
Single cell RNA-sequencing in mouse kidney shows *COL4A3, SNCAIP*, and *BMP7* are specifically expressed in podocytes. Mean expression values of the genes were calculated in each cluster. The color scheme is based on z-score distribution; the map shows genes with z-score>2. In the heatmap, each row represents one gene and each column is single cell type. Percentages of assigned cell types are summarized in the right panel. Endo, containing endothelial, vascular, and descending loop of Henle; Podo, podocyte; PT, proximal tubule; LOH, ascending loop of Henle; DCT, distal convoluted tubule; CD-PC, collecting duct principal cell; CD-IC, collecting duct intercalated cell; CD-Trans, collecting duct transitional cell; Fib, fibroblast; Macro, macrophage; Neutro, neutrophil; lymph, lymphocyte; NK, natural killer cell.

## DISCUSSION

Our genome-wide analysis of 19,406 participants with T1D identified 16 genome-wide significant loci associated with DKD, four of which remained significant after a conservative correction for multiple testing. Four of the 16 genome-wide significant signals are in or near genes with known or suggestive biology related to renal function/collagen *(COL4A3, BMP7, COLEC11*, and *DDR1)*, but this is the first time that naturally occurring variation (MAF > 1%) in these loci has been associated with DKD. Our most significant signal was a protective missense variant in *COL4A3*, rs55703767, reaching both genome-wide and study-wide significance with multiple definitions of DKD. Moreover, this variant demonstrated significant association with GBM width and with the degree of fibrosis in renal tubules and glomeruli, and its effect was dependent on glycemia.

COL4A3, with COL4A4 and COL4A5, make up the so-called “novel chains” of type IV collagen^48^, which together play both structural and signaling roles in the GBM. Specifically, COL4A3 is known to bind a number of molecules including integrins, heparin and heparin sulfate proteoglycans, and other components of the GBM such as laminin and nidogen. These interactions mediate the contact between cells and the underlying collagen IV basement membrane, and regulate various processes essential to embryonic development and normal physiology including cell adhesion, proliferation, survival and differentiation. Dysregulation of these interactions has been implicated in several pathological conditions including CKD^49^.

Mutations in *COL4A3* are responsible for the autosomal recessive form of Alport syndrome, a progressive inherited nephropathy, as well as benign familial hematuria, characterized by thin (or variable width) GBM, and thought to be a milder form of Alport syndrome^50^. The rs55703767 SNP is predicted to alter the third amino acid of the canonical triple-helical domain sequence of Glycine (G)-X-Y (where X and Y are often proline (P) and hydroxyproline (Y), respectively) from G-E-D (D=Aspartic) to G-E-Y^51^, potentially impacting the structure of the collagen complex. In addition, a recent study^52^ of candidate genes involved in renal structure reported rs34505188 in *COL4A3* (not in linkage disequilibrium with rs55703767, r^2^=0.0006) to be associated with ESRD in African Americans with T2D (MAF=2%, OR=1.55, *P*=5×10^-4^). Together with the trend towards association we have seen in SUMMIT and the glycemic interaction we have reported here, these findings suggest variation in *COL4A3* may be associated with DKD in T2D as well.

Given its association as a protective SNP, we can speculate that the rs55703767 variant may confer tensile strength or flexibility to the GBM, which may be of particular relevance in the glomerular hypertension associated with DKD. Alternatively, *COL4A3* may regulate the rates of production and/or turnover of other GBM components, affecting GBM width changes in diabetes. How these effects might confer protection in a manner dependent on ambient glucose concentrations is unknown. Future mechanistic studies will be required to determine the precise role of this variant in DKD; elucidation of its interaction with glycemia in providing protection might be relevant to other molecules implicated in diabetic complications.

In keeping with the collagen pathway, the synonymous exonic variant rs118124843, which reached genome-wide significance for the “Micro” phenotype, is located near *DDR1*, the gene encoding the discoidin domain-containing receptor 1. Based on chromatin conformation interaction data from Capture HiC Plotter (CHiCP),^53^ the rs118124843 containing fragment interacts with six gene promoter regions, including *DDR1*, suggesting that the variant regulates *DDR1* expression across multiple tissues (Table S11). DDR1 is a collagen receptor^54^ shown to bind type IV collagen^55^, and is highly expressed in kidneys, particularly upon renal injury^56^. Upon renal injury, *Ddr1-deficient* mice display lower levels of collagen^57^, decreased proteinuria, and an increased survival rate compared to wild-type controls^58^, with *Ddr1/Col4a3* double knockout mice displaying protection from progressive renal fibrosis and prolonged lifespan compared to *Col4a3* knockout mice alone^57^. Thus, through its role in collagen binding DDR1 has been suggested as a possible therapeutic target for kidney disease^57^.

The association of rs12615970, an intronic variant on chromosome 2 near the *COLEC11* gene, met genome-wide significance for the CKD phenotype, as well as nominal significance for multiple albuminuria-based traits. The rs12615970 containing fragment was found to interact with *COLEC11, ALLC*, and *ADI1* transcription start sites in chromatin conformation data on GM12878 cell line (Table S11)^53,59^. Collectin-11 is an innate immune factor synthesized by multiple cell types, including renal epithelial cells with a role in pattern recognition and host defense against invasive pathogens, through binding to fructose and mannose sugar moieties^60,61^. Mice with kidney-specific deficiency of *COLEC11* are protected against ischemia-induced tubule injury due to reduced complement deposition^62^, and mutations in *COLEC11* have been identified in families with 3MC syndrome, a series of rare autosomal recessive disorders resulting in birth defects and abnormal development, including kidney abnormalities^63^.

The intronic variant rs144434404, associated at study-wide significance with the microalbuminuria phenotype, resides within the bone morphogenetic protein 7 *(BMP7)* gene. *BMP7* encodes a secreted ligand of the transforming growth factor-beta superfamily of proteins. Developmental processes are regulated by the BMP family of glycosylated extracellular matrix molecules, via serine/threonine kinase receptors and canonical Smad pathway signaling. Coordinated regulation of both BMP and BMP-antagonist expression is essential for developing tissues, and changes in the levels of either BMP or BMP-antagonists can contribute to disease progression such as fibrosis and cancer^64^. BMP7 is required for renal morphogenesis, and *Bmp7* knockout mice die soon after birth due to reduced ureteric bud branching^65–67^. Maintenance of *Bmp7* expression in glomerular podocytes and proximal tubules of diabetic mice prevents podocyte loss and reduces overall diabetic renal injury^27^. More recently, we have identified a mechanism through which BMP7 orchestrates renal protection through Akt inhibition and highlights Akt inhibitors as potential anti-fibrotic therapeutics^68^. It is also noteworthy that the BMP7 antagonist grem-1 is implicated in DKD^69–71^ and gremlin has been implicated as a biomarker of kidney disease^72^.

Strengths of this analysis include the large sample size, triple that of the previous largest GWAS; the uniform genotyping and quality control procedures; standardized imputation for all studies (1,000 Genomes reference panel); the inclusion of exome array content; the exploration of multiple standardized phenotype definitions of DKD; and supportive data from various sources of human kidney samples. Several of the loci identified have known correlations with kidney biology, suggesting that these are likely true associations with DKD. However, we acknowledge a number of limitations. First, nine variants have low MAF and were driven by only two cohorts, indicating that further validation will be required to increase confidence in these associations. Second, seven variants were significantly associated with microalbuminuria only, a trait shown to be less heritable in previous studies. Even though the gene-level, gene set and pathway analyses had limited power, these analyses identified several additional potential DKD loci and pathways, some with relevance to kidney biology, that require further follow-up.

Diabetic complications are unquestionably driven by hyperglycemia and partially prevented by improved glycemic control in both T1D and T2D, but there has been doubt over what contribution, if any, inherited factors contribute to disease risk. In line with previous genetic studies, this study with a markedly expanded sample size identified several loci strongly associated with DKD risk. These findings suggest that larger studies, aided by novel analyses and including T2D, will continue to enhance our understanding of the complex pathogenesis of DKD, paving the way for molecularly targeted preventive or therapeutic interventions.

## AUTHOR CONTRIBUTIONS

Conceptualization, C.G., A.P.M., P-H.G., J.N.H, and J.C.F.; Formal Analysis, R.M.S., J.B.C., N.S., E.V., C-D.L., J-J.C., M.G.P., A.M.S., C.Q., V.N., J.P., A.L., R.M., H.K., R.K., B.E.K., A.J.C., A.D.P., W-M.C., N.R.v-Z., and S.O-G.; Investigation M.G.P., A.M.S., J.S. C-X.Q., V.N., J.K.H, A.J.M, J.K.S-B., D.M.M., A.P.M., A.D.P., C.F., E.A., T.A., M.L., C.M.B-K., H.M.K., S.M., D. T., B.G., S.B.B., C.N.A.P., L.S., B.P., R.W., V.R., V.P., E.P., L.R., R.V., N.M.P., L.C.G., M.I.M., H.F.G., A.M., H.F., K.B., P.R., T.C., G.Z., M.M., S.H., C.F., W-M.C., and S.O-G; Resources A.M.S., V.H., A.J.M, R.G.N., M.L.C., M.M, X.G., J.K.S-B., D.M.M., J.G., R.G.M., A.P.M., A.D.P., W-M.C., and S.O-G; Data Curation L.T.H., C-D. L., J-J.C., A.M.S., V.H., J.S., A.J.M., R.K., B.E.K., K.E.L., X.G., M.M., M.L.C., J.K.S-B., D.M.M., J.G., R.G.M., A.P.M., W-M.C., and S.O-G; Writing – Original Draft R.M.S., J.N.T., N.S., J.B.C., E.P.B., D.A., R.D., M.F.H., L.T.H., A.D.P., and C.G.; Writing – Review & Editing R.M.S., J.N.T., N.S., J.B.C., E. P.B., D.A., R.D., M.F.H., L.T.H., A.D.P., C.G., C.Q., J.P., J.S., M.M., M-L.C., I.H.dB., F.M., A.J.M., G.M., A.P.M., K.S., M.I.M., L.G., S.S.R., C.F., J.N.H., and J.C.F.; Visualization R.M.S, J.N.T., N.S., J.B.C., M.P.G., C.Q., C-D.L., J-J.C., J.P., J.S., R.K., B.E.K., K.E.L., A.J.M., G.M., A.P.M., and A.D.P.; Project Administration J.N.T.; Supervision A.D.P., A.K., P-H.G., C.G., A.P.M., H.M.C., S.S.R., M.K., K.S., J.N.H, and J.C.F.; Funding Acquisition A.D.P., A.K., J.C.F., and J.N.H.

## ACKNOWLEDGEM ENTS

This study was supported by a grant from the JDRF (17-2013-7) and National Institute of Diabetes and Digestive and Kidney Diseases (NIDDK) grants R01 DK081923 and R01 DK105154. R.M.S was supported in part by JDRF # 3-APF-2014-111-A-N, and National Heart, Lung and Blood Institute (NHLBI) R00 HL122515. J.N.T. was supported by NIDDK K12-DK094721. N.S. received funds from the European Foundation for the Study of Diabetes (EFSD) Young Investigator Research Award funds and Academy of Finland (299200). C.G., D. A., E.B., E.M., M.H. and R.D. are supported by SFI-HRB US Ireland Research Partnership SFI15/US/B3130. The FinnDiane study was funded by JDRF (17-2013-7), Folkhälsan Research Foundation, the Wilhelm and Else Stockmann Foundation, the Liv och Hälsa Foundation, Helsinki University Central Hospital Research Funds (EVO), the Novo Nordisk Foundation, and Academy of Finland (275614). The EDC study was supported by NIDDK grant DK34818 and by the Rossi Memorial Fund. The WESDR study was funded by National Eye Institute EY016379. The Sweden study was supported by Family Erling-Persson (K. B.) and Stig and Gunborg Westman foundations (H.F.G.). AusDiane acknowledges Dr. Bernhard Baumgartner from Department of Medicine, Diakonissen-Wehrle Hospital, Salzburg, Austria. LatDiane acknowledges the following doctors and researchers: A. Bogdanova, D. Grikmane, I. Care, A. Fjodorova, R. Graudiņa, A. Valtere, D. Teterovska, I. Dzīvīte-Krišāne, I. Kirilova, U. Lauga-Tūriņa, K. Geldnere, D. Seisuma, N. Fokina, S.Šteina, L. Jaunozola, I. Balcere, U. Gailiša, E. Menise, S. Broka, L. Akmene, N. Kapļa, J. Nagaiceva, A. Pētersons, A. Lejnieks, I. Konrāde, I. Salna, A. Dekante, A. Grāmatniece, A. Saliņa, A. Silda, V. Mešečko, V. Mihejeva, K. Kudrjavceva, I. Marksa, M. Cirse, D. Zeme, S. Kalva-Vaivode, J. Kloviņš, L. Nikitina-Zaķe, S. Skrebinska, Z. Dzērve, R. Mallons. LitDiane acknowledges Dr. Jurate Lasiene, Prof. Dzilda Velickiene, Dr. Vladimiras Petrenko and Nurses from the Department of Endocrinology, Hospital of Lithuanian University of Health Sciences. RomDiane acknowledges the following doctors and researchers: M. Anghel, DM Cheta, BA Cimpoca, D. Cimponeriu, CN Cozma, N. Dandu, D. Dobrin, I Duta, AM Frentescu, C. Ionescu-Tirgoviste, R. Ionica, R. Lichiardopol, AL Oprea, NM Panduru, A. Pop, G. Pop, R. Radescu, S. Radu, M. Robu, C. Serafinceanu, M. Stavarachi, O Tudose from the “N.C Paulescu” National Institute for Diabetes Nutrition and Metabolic Diseases from Bucharest, and M. Bacu, D. Clenciu, C. Graunteanu, M. Ioana, E. Mota and M. Mota from Craiova Emergency County Hospital.

SUMMIT Consortium acknowledgements: This work was supported by the following funders: IMI (SUMMIT 115006); Wellcome Trust 098381, 090532, 106310; NIH R01-MH101814; and JDRF 2-SRA-2014-276-Q-R. SDR: Thanks to Maria Sterner and Malin Neptin for GWAS genotyping in the Scannia Diabetes Registry. Grants: Swedish Research Council, ERC-Adv res grant 269045-GENE TARGET T2D, Academy of Finland grantas 263401 and 267882, Sigrid JUselius Foundation. FinnDiane: We thank M. Parkkonen, A. Sandelin, A-R. Salonen, T. Soppela and J. Tuomikangas for skillful laboratory assistance in the FinnDiane study. We also thank all the subjects who participated in the FinnDiane study and gratefully acknowledge all the physicians and nurses at each centre involved in the recruitment of participants (Table S12). The FinnDiane Study was supported by grants from the Folkhälsan Research Foundation, the Wilhelm and Else Stockmann Foundation, the Liv och Hälsa Foundation, Helsinki University Central Hospital Research Funds (EVO), the Novo Nordisk Foundation, European Foundation for the Study of Diabetes (EFSD) Young Investigator Research Award funds, and the Academy of Finland (275614 and 299200). GoDARTS: We are grateful to all the participants in this study, the general practitioners, the Scottish School of Primary Care for their help in recruiting the participants, and to the whole team, which includes interviewers, computer and laboratory technicians, clerical workers, research scientists, volunteers, managers, receptionists, and nurses. The study complies with the Declaration of Helsinki. We acknowledge the support of the Health Informatics Centre, University of Dundee for managing and supplying the anonymised data and NHS Tayside, the original data owner. The Wellcome Trust United Kingdom Type 2 Diabetes Case Control Collection (GoDARTS) was funded by The Wellcome Trust (072960/Z/03/Z, 084726/Z/08/Z, 084727/Z/08/Z, 085475/Z/08/Z, 085475/B/08/Z) and as part of the EU IMI-SUMMIT program.

We acknowledge the following conflicts of interest: P-H.G. has received investigator-initiated research grants from Eli Lilly and Roche, is an advisory board member for AbbVie, AstraZeneca, Boehringer Ingelheim, Cebix, Eli Lilly, Janssen, Medscape, Merck Sharp & Dohme, Novartis, Novo Nordisk and Sanofi; and has received lecture fees from AstraZeneca, Boehringer Ingelheim, Eli Lilly, Elo Water, Genzyme, Merck Sharp & Dohme, Medscape, Novo Nordisk and Sanofi. M.I.M. is a Wellcome Investigator and an NIHR Senior Investigator. The views expressed in this article are those of the author(s) and not necessarily those of the NHS, the NIHR, or the Department of Health. He serves on advisory panels for Pfizer, NovoNordisk, and Zoe Global; has received honoraria from Merck, Pfizer, NovoNordisk and Eli Lilly; has stock options in Zoe Global; and has received research funding from Abbvie, Astra Zeneca, Boehringer Ingelheim, Eli Lilly, Janssen, Merck, NovoNordisk, Pfizer, Roche, Sanofi Aventis, Servier and Takeda. P.R. has received consultancy and/or speaking fees (to his institution) from AbbVie, Astellas, AstraZeneca, Bayer, Boehringer Ingelheim, Bristol-Myers Squibb, Eli Lilly, MSD, Novo Nordisk and Sanofi Aventis and research grants from AstraZeneca and Novo Nordisk, and shares in Novo Nordisk.

## Online METHODS

### Cohorts in GWAS

The GWAS meta-analysis included up to 19,406 patients with type 1 diabetes and of European origin from 17 cohorts: The Austrian Diabetic Nephropathy Study (AusDiane); The Coronary Artery Calcification in Type 1 Diabetes (CACTI)^73^; the Diabetes Control and Complications Trial/Epidemiology of Diabetes Interventions and Complications (DCCT/EDIC)^23,24^; Pittsburgh Epidemiology of Diabetes Complications Study (EDC)^74^; The Finnish Diabetic Nephropathy (FinnDiane) Study^12,21^; French and Belgian subjects from the Genetics of Diabetic Nephropathy (GENEDIAB)^75^ and Genesis^76^ studies; Genetics of Kidneys in Diabetes US Study (GoKinD) from George Washington University (GWU-GoKinD)^77^; patients from the Joslin Kidney Study^77,78^; individuals with T1D from Italy^12^; The Latvian Diabetic Nephropathy Study (LatDiane)^79^; The Lithuanian Diabetic Nephropathy Study (LitDiane) [Reference pending, submitted]; The Romanian Diabetic Nephropathy Study (RomDiane)^80^; The Scottish Diabetes Research Network Type 1 Bioresource (SDRNT1BIO)^72,81^; individuals with T1D from Steno Diabetes Center^82^; individuals with T1D from Uppsala, Sweden^83,84^; UK GoKinD, Warren 3 and All Ireland (UK-ROI) study^40^; and The Wisconsin Epidemiologic Study of Diabetic Retinopathy (WESDR)^85^. All participants gave informed consent and all studies were approved by ethics committees from all participating institutions.

### RASS cohort

RASS was a double-blind placebo-controlled randomized trial of the angiotensin converting enzyme inhibitor (ACEi) enalapril and the angiotensin II receptor blocker (ARB) losartan on renal pathology among 285 normoalbuminuric, normotensive subjects with T1D and had normal or increased measured glomerular filtration rate (>90 ml/min/1.73m^2^)^17^. Beginning in 2005, participants were recruited from three centers: University of Minnesota (Minneapolis, Minnesota), McGill University (Montreal, Canada) and University of Toronto (Toronto, Canada) and included those with 2 to 20 years of diabetes and excluded those on any antihypertensive medications. Written informed consent was obtained from each participant and the study was approved by the relevant institutional review boards. RASS study participants were followed for 5 years with percutaneous kidney biopsy completed prior to randomization and at 5 years. Structural parameters measured by electron microscopy on biopsy included GBM width, measured by the electron microscopic orthogonal intercept method^17^.

### SUMMIT consortium

The SUMMIT consortium included up to 5193 subjects with type 2 diabetes, with and without kidney disease, of European ancestry. All studies were approved by ethics committees from relevant institutions and all participants gave informed consent^13^. Complete list of SUMMIT Consortium members provided in Table S13.

### Genotyping

Samples were genotyped on the HumanCore BeadChip (Illumina, San Diego, CA, USA), which contains 250,000 genome-wide tag SNPs (and other variants) and over 200,000 exome-focused variants. All samples were passed through a stringent quality control protocol. Following initial genotype calling with Illumina software, all samples were re-called with zCall, a calling algorithm specifically designed for rare SNPs from arrays. Once calling was completed for all cohorts, our pipeline updated variant orientation and position aligned to hg19 (Genome Reference Consortium Human Build 37, GRCh37). Variant names were updated using 1000 Genomes as a reference. The data were then filtered for low quality variants (e.g. call rates <95% or excessive deviation from Hardy-Weinberg equilibrium) or samples (e.g. call rates <98%, gender mismatch, extreme heterozygosity). Principal Component Analysis (PCA) was performed separately for each cohort in order to empirically detect and exclude outliers with evidence of non-European ancestry. Genotypes were expanded to a total of approximately 49 million by imputation, using 1,000 Genomes Project (phase 3 version 5) as a reference.

### Genotyping of RASS cohort

All RASS participants contributed DNA for genotyping on the Illumina HumanOmni1-Quad and HumanCoreExome beadchip arrays. Genotypes were called using BeadStudio/Genomestudio software (Illumina^®^). Quality control (QC) measures included removing duplicate samples, samples with evidence of contamination (heterozygosity range 0.25-0.32) and those with cryptic relatedness identity-by-state (IBS) (n=24). Principal component analyses were completed and 7 non-European outliers were removed. Of those genotyped, 1 participant was missing kidney biopsy data.

### Genotyping in SUMMIT consortium

Cohorts were genotyped on the Affymetrix SNP 6.0, the Illumina Omni express and the Illumina 610Quad arrays. QC measures included filtering out low frequency (<1% MAF) variants, filtering out low quality variants or samples, removal of duplicate samples, and removal of non-European samples based on principal component analysis.^13^

### Human kidney samples from University of Pennsylvania cohort for RNA-sequencing and cis-eQTL analysis

Human kidney tissue collection was approved by the University of Pennsylvania Institutional Review Board. Kidney samples were obtained from surgical nephrectomies. Nephrectomies were de-identified, and the corresponding clinical information was collected through an honest broker; therefore, no consent was obtained from the subjects. Tubular and glomerular eQTL data sets were generated by 121 samples of tubules and 119 samples of glomeruli, respectively. The cis window was defined as 1 megabase up- and downstream of the transcriptional start site (±1Mb). Whole kidney cis-eQTL (further just referred to as eQTL) data set was generated from 96 human samples were obtained from The Cancer Genome Atlas (TCGA) through the TCGA Data portal^42^.

### Mouse kidney single cell RNA-sequencing

Animal studies were approved by the Institutional Animal Care and Use Committee of the University of Pennsylvania. We mated Cdh16^Cre^ mice (Jackson Lab, 012237), Nphs2^Cre^ mice (Jackson Lab, 008205) and Scl^Cre^ mice (MGI number is 3579158) with Tomato-GFP (mT/mG) mice (Jackson Lab, 007576) to generate Cdh16^Cre^mT/mG, Scl^Cre^mT/mG and Nphs2^cre^mT/mG mice^51^.

### Genomic features of human kidney

Human kidney-specific chromatin immunoprecipitation followed by sequencing (ChIP-seq) data can be found at GEO: GSM621634, GSM670025, GSM621648, GSM772811, GSM621651, GSM1112806, GSM621638. Different histone markers were combined into chromatin states using ChromHMM^45^.

### Glomerular basement membrane measurement in RASS cohort

RASS study participants were followed for 5 years with percutaneous kidney biopsy completed prior to randomization and at 5 years. Structural parameters measured by electron microscopy on biopsy included GBM width, measured by the electron microscopic orthogonal intercept method^17^.

### RNA-sequencing of human kidney samples in the University of Pennsylvania cohort

Human kidney tissue was manually microdissected under a microscope in RNAlater for glomerular and tubular compartments. The local renal pathologist performed an unbiased review of the tissue section by scoring multiple parameters, and RNA were prepared using RNAeasy mini columns (Qiagen, Valencia, CA) according to manufacturer’s instructions. RNA quality was assessed with the Agilent Bioanalyzer 2100 and RNA integrity number scores above 7 were used for cDNA production. The library was prepared in the DNA Sequencing Core at University of Texas Southwestern Medical Center. One microgram total RNA was used to isolate poly(A) purified mRNA using the Illumina TruSeq RNA Preparation Kit. We sequenced samples for single-end 100bp, and the annotated RNA counts (fastq) were calculated by Illumina’s CASAVA 1.8.2. Illumina sequence quality was surveyed with FastQC. Adaptor and lower-quality bases were trimmed with Trim-galore. Trimmed reads were aligned to the Gencode human genome(GRCh37) with STAR-2.4.1d. The readcount of each sample was obtained using HTSeq-0.6.1 (htseq-count) and then normalized fragments per kilobase million values were used to perform association analysis with fibrosis and sclerosis using linear regression.

### Mouse kidney single cell RNA-sequencing

Kidneys were harvested from 4 to 8-week-old male mice with C57BL/6 background and dissociated into single cell suspension as described in our previous study^47^. The single cell sequencing libraries were sequenced on an Illumina HiSeq with 2×150 paired-end kit. The sequencing reads were demultiplexed, aligned to the mouse genome (mm10) and processed to generate gene-cell data matrix using Cell Ranger 1.3 (http://10xgenomics.com)^47^.

### RNAseq and microarray profiling of human kidney samples from the Pima cohort

Kidney biopsy samples from the Pima Indian cohort were manually micro-dissected into 119 glomerular and 100 tubule-interstitial tissues to generate gene expression profiles^86^. Expression profiling in the Pima Indian cohort kidney biopsies was carried out using Affymetrix GeneChip Human Genome U133 Array and U133Plus2 Array, as reported previously, and Affymetrix Human Gene ST Genechip 2.1^87,88^, and on RNA-seq (Illumina). The libraries were prepared using the ClonTech SMARTSeq v4 Ultra Low Input polyA selection kit. Samples were sequenced on a HiSeq 4000, single end, 75bp. Mapping to human reference genome GRCh38.7 was performed with STAR 2.5.2b (https://github.com/alexdobin/STAR). For annotation and quantification of mapping results we used cufflinks, cuffquant and cuffnorm in version 2.2.1 (https://cole-trapnell-lab.github.io/cufflinks/). After mapping and quantification, PCA and Hierarchical Clustering was used to identify outliers and reiterated until no more outliers could be identified.

## STATISTICAL ANALYSIS

### GWAS Analysis

Participant renal status was evaluated on the basis of both albuminuria and eGFR. We defined a total of 10 different case-control outcomes to cover the different aspects of renal complications (Figure 1). Five comparisons (“All vs. ctrl”, “Micro”, “DN”, “Macro”, and “ESRD vs. macro”) were based on albuminuria, measured by albumin excretion rate (AER) from overnight or 24-h urine collection, or by albumin creatinine ratio (ACR). Two out of three consecutive collections were required (when available) to classify the renal status of subjects as either normoalbuminuria, microalbuminuria, macroalbuminuria, or ESRD; for detailed thresholds, see **Table S9**. Controls with normal AER were required to have a minimum diabetes duration of 15 years; subjects with microalbuminuria/ macroalbuminuria/ ESRD were required to have minimum diabetes duration of 5/ 10/ 10 years, respectively, in order to exclude renal complications of non-diabetic origins. Two comparisons (“ESRD vs. ctrl” and “ESRD vs. non-ESRD”) were based on presence of end-stage renal disease as defined by eGFR< 15 mL/min or dialysis or renal transplant. Two phenotypes (“CKD” and “CKD extreme”) were defined based on estimated glomerular filtration rate (eGFR; evaluated with the CKD-EPI formula): Controls had eGFR ≥ 60ml/min/1.73m^2^ for both phenotypes, and minimum of 15 years of diabetes duration; cases had eGFR <60ml/min/1.73m^2^ for the “CKD” phenotype, and eGFR <15 ml/min/1.73m^2^ or dialysis or renal transplant for the “CKD extreme” phenotype, and minimum of 10 years of diabetes duration. For the “CKD-DN” phenotype that combined both albuminuria and eGFR data, controls were required to have both eGFR ≥60ml/min/1.73m^2^ and normoalbuminuria; cases had both eGFR <45ml/min/1.73m^2^ and micro-or macroalbuminuria, or ESRD.

A genome-wide association analysis of each of the case-control definitions was performed using logistic regression under an additive genetic model, adjusting for age, sex, diabetes duration, study site (where applicable) and principal components. As disease onset and progression is also closely related to BMI and HbA1c levels,^89^ we conducted a second set of analyses adjusting for BMI and HbA1c which we refer to as our fully adjusted covariate model. Allele dosages were used to account for imputation uncertainty. Inverse-variance fixed effects metaanalysis was performed using METAL and the following filters: INFO score >0.3, minor allele count >10, and presence of variant in at least two cohorts. The X chromosome was similarly analyzed for males and females both separately and in a combined analysis, with the exception of using hard call genotypes in place of allele dosages. The study-wide significance threshold (*P*<6.76×10^-9^) was calculated by applying a Bonferroni correction to the traditional GWAS threshold (*P*<5.00×10^-8^), based on the number of effectively independent tests, using methods previously described on the eigenvalues of the GWAS summary statistics correlation matrix^14^.

### SUMMIT GWAS analysis

Genome-wide association analyses were performed for DKD trait definitions harmonized with seven of our primary T1D analyses: “DN”, “Micro”, “Macro”, “ESRD”, “ESRD vs. non-ESRD”, “CKD”, and “CKD-DN” under an additive model, adjusting for age, gender and duration of diabetes.

### RASS GBM width analysis

We completed linear regression of the *COL4A3* variant (rs55703767) and within person mean GBM width (nm) from both baseline and 5 year measures, in additive and genotypic genetic models. Both univariate and multivariate analyses were run including sex, baseline age and diabetes duration, within person mean HbA1c over 5 years, indicators for treatment group assignment and treatment center. A two-sided significance threshold of alpha <0.05 was applied.

### Gene and gene set analysis

PASCAL gene and pathway scores were conducted on all 20 sets of GWAS summary statistics (10 outcomes and 2 covariate models). Gene scores were derived using the sum option, averaging association signal across each gene using the default 50kb window size. Pathway scores were then computed from pathway member gene scores where membership was defined using default pathway libraries from BioCarta, REACTOME, and KEGG. Using a similar approach, MAGMA (v1.06) gene and pathway scores were conducted on all GWAS summary statistics using both the default gene region defined by the transcription start and stop sites and a 5kb window definition. MAGMA pathway analysis included all 1077 of the PASCAL reported libraries plus an additional 252 pathways included in MSigDB canonical pathway set. MAGENTA (vs2, July 2011) pathway analysis included 4725 pathways with a minimum of five genes within the gene set. Gene sets were obtained with the MAGENTA distribution and included Gene ontology terms, PANTHER sets (biological processes, molecular functions, metabolic and signaling pathways), KEGG pathways, and Ingenuity pathways. DEPICT gene set enrichment uses a more comprehensive collection of gene sets that allows genes to have a continuous probability for gene set membership. We conducted DEPICT individually on all 20 sets of GWAS summary statistics with P< 1.0 × 10^-5^. We conducted two additional pooled analyses using genome-wide minimum P-values from: 1) All 20 analyses (10 phenotypes and 2 covariate models) and 2) Sixteen analyses of the 8 most related phenotypes (8 phenotypes and 2 covariate models) which excluded ESRD vs Macro and Micro.

### Human kidney cis-eQTL analysis (University of Pennsylvania data)

Nominal p-values were calculated for each SNP-gene pair with FastQTL using linear regression with an additive effects model, and adjusted by six genotype PCs.

### RNA-sequencing of human kidney samples (University of Pennsylvania data)

Normalized fragment per kilobase million values were used to perform association analysis with fibrosis and sclerosis using linear regression.

### eQTL analysis (Pima data)

Analysis was performed with Robust Multi-array Average quantile normalization^90^ after removing probes overlapping with variants identified by WGS. Batch effects between platforms were corrected using *ComBat^91^* and unknown batch effects were also adjusted using singular value decomposition with first four eigenvectors. eQTL mapping was performed using EPACTS (https://genome.sph.umich.edu/wiki/EPACTS) software tool using linear mixed model accounting for hidden familial relatedness, after inverse Gaussian transformation of expression levels, adjusting for age and sex.

### Mouse kidney single cell RNA-sequencing

To calculate the average expression level for each cluster, a z-score of normalized expression value was first obtained for every single cell. Then, we calculated the mean z-scores for individual cells in the same cluster, resulting in 16 values for each gene.

## DATA AND SOFTWARE AVAILABILITY

All cohorts can share genome-wide meta-analysis summary statistics. Individual level genotype data: due to restrictions set by the study consents and by EU and national regulations, individual genotype data cannot be shared for all cohorts.

## REFERENCES

1. Centers for Disease Control and Prevention. National Diabetes Statistics Report, 2017 Estimates of Diabetes and Its Burden in the United States Background. in Atlanta, GA: Centers for Disease Control and Prevention, US Department of Health and Human Services; 2017.

2. Tuttle, K.R. et al. Diabetic kidney disease: a report from an ADA Consensus Conference. Am J Kidney Dis 64, 510–33 (2014).

3. Krolewski, M., Eggers, P.W. & Warram, J.H. Magnitude of end-stage renal disease in IDDM: a 35 year follow-up study. Kidney Int 50, 2041–2046 (1996).

4. Harjutsalo, V., Katoh, S., Sarti, C., Tajima, N. & Tuomilehto, J. Population-based assessment of familial clustering of diabetic nephropathy in type 1 diabetes. Diabetes 53, 2449–54 (2004).

5. Quinn, M., Angelico, M.C., Warram, J.H. & Krolewski, A.S. Familial factors determine the development of diabetic nephropathy in patients with IDDM. Diabetologia 39, 940–5 (1996).

6. Seaquist, E., Goetz, F., Rich, S. & Barbosa, J. Familial clustering of diabetic kidney disease. Evidence for genetic susceptibility to diabetic nephropathy. N Engl J Med 320, 1161–1165 (1989).

7. Sandholm, N. et al. The Genetic Landscape of Renal Complications in Type 1 Diabetes. J Am Soc Nephrol 28, 557–574 (2017).

8. Iyengar, S.K. et al. Genome-Wide Association and Trans-ethnic Meta-Analysis for Advanced Diabetic Kidney Disease: Family Investigation of Nephropathy and Diabetes (FIND). PLoS Genet 11, e1005352 (2015).

9. Pattaro, C. et al. Genetic associations at 53 loci highlight cell types and biological pathways relevant for kidney function. Nat Commun 7, 10023 (2016).

10. Sandholm, N. et al. Genome-wide association study of urinary albumin excretion rate in patients with type 1 diabetes. Diabetologia 57, 1143–53 (2014).

11. Sandholm, N. et al. Chromosome 2q31.1 Associates with ESRD in Women with Type 1 Diabetes. Journal of the American Society of Nephrology 24, 1537–1543 (2013).

12. Sandholm, N. et al. New susceptibility loci associated with kidney disease in type 1 diabetes. PLoS Genet 8, e1002921 (2012).

13. van Zuydam, N.R. et al. A Genome-Wide Association Study of Diabetic Kidney Disease in Subjects With Type 2 Diabetes. Diabetes 67, 1414–1427 (2018).

14. Li, J. & Ji, L. Adjusting multiple testing in multilocus analyses using the eigenvalues of a correlation matrix. Heredity (Edinb) 95, 221–7 (2005).

15. Yagame, M. et al. Differential distribution of type IV collagen chains in patients with diabetic nephropathy in non-insulin-dependent diabetes mellitus. Nephron 70, 42–8 (1995).

16. Caramori, M.L., Parks, A. & Mauer, M. Renal lesions predict progression of diabetic nephropathy in type 1 diabetes. J Am Soc Nephrol 24, 1175–81 (2013).

17. Mauer, M. et al. Renal and retinal effects of enalapril and losartan in type 1 diabetes. N Engl J Med 361, 40–51 (2009).

18. Fufaa, G.D. et al. Structural Predictors of Loss of Renal Function in American Indians with Type 2 Diabetes. Clin J Am Soc Nephrol 11, 254–61 (2016).

19. Mauer, M., Caramori, M.L., Fioretto, P. & Najafian, B. Glomerular structural-functional relationship models of diabetic nephropathy are robust in type 1 diabetic patients. Nephrol Dial Transplant 30, 918–23 (2015).

20. Gorski, M. et al. 1000 Genomes-based meta-analysis identifies 10 novel loci for kidney function. Sci Rep 7, 45040 (2017).

21. Thorn, L.M. et al. Metabolic syndrome in type 1 diabetes: Association with diabetic nephropathy and glycemic control (the FinnDiane study). Diabetes Care 28, 2019–2024 (2005).

22. Morris, A.D. et al. The diabetes audit and research in Tayside Scotland (DARTS) study: electronic record linkage to create a diabetes register. DARTS/MEMO Collaboration. BMJ 315, 524–8 (1997).

23. Nathan, D.M. & Group, D.E.R. The diabetes control and complications trial/epidemiology of diabetes interventions and complications study at 30 years: overview. Diabetes Care 37, 9–16 (2014).

24. The Diabetes Control and Complications Trial Research Group. Implementation of treatment protocols in the Diabetes Control and Complications Trial. Diabetes Care 18, 361–76 (1995).

25. Blunsom, N.J., Gomez-Espinosa, E., Ashlin, T.G. & Cockcroft, S. Mitochondrial CDP-diacylglycerol synthase activity is due to the peripheral protein, TAMM41 and not due to the integral membrane protein, CDP-diacylglycerol synthase 1. Biochim Biophys Acta Mol Cell Biol Lipids 1863, 284–298 (2018).

26. Tamura, Y. et al. Tam41 is a CDP-diacylglycerol synthase required for cardiolipin biosynthesis in mitochondria. Cell Metab 17, 709–18 (2013).

27. Wang, S. et al. Renal bone morphogenetic protein-7 protects against diabetic nephropathy. J Am Soc Nephrol 17, 2504–12 (2006).

28. Sandholm, N. et al. Confirmation of GLRA3 as a susceptibility locus for albuminuria in Finnish patients with type 1 diabetes. Sci Rep 8, 12408 (2018).

29. Lamparter, D., Marbach, D., Rueedi, R., Kutalik, Z. & Bergmann, S. Fast and Rigorous Computation of Gene and Pathway Scores from SNP-Based Summary Statistics. PLoS Comput Biol 12, e1004714 (2016).

30. de Leeuw, C.A., Mooij, J.M., Heskes, T. & Posthuma, D. MAGMA: generalized gene-set analysis of GWAS data. PLoS Comput Biol 11, e1004219 (2015).

31. Bell, C.G. et al. Genome-wide DNA methylation analysis for diabetic nephropathy in type 1 diabetes mellitus. BMC Medical Genomics 3 (2010).

32. Sapienza, C. et al. DNA methylation profiling identifies epigenetic differences between diabetes patients with ESRD and diabetes patients without nephropathy. Epigenetics 6, 20–8 (2011).

33. Loo, Y.M. & Gale, M., Jr. Immune signaling by RIG-I-like receptors. Immunity 34, 680–92 (2011).

34. Mollen, K.P. et al. Emerging paradigm: toll-like receptor 4-sentinel for the detection of tissue damage. Shock 26, 430–7 (2006).

35. Mora, C. & Navarro, J.F. Inflammation and diabetic nephropathy. Curr Diab Rep 6, 463–8 (2006).

36. Wada, J. & Makino, H. Inflammation and the pathogenesis of diabetic nephropathy. Clin Sci (Lond) 124, 139–52 (2013).

37. Lin, M. et al. Toll-like receptor 4 promotes tubular inflammation in diabetic nephropathy. J Am Soc Nephrol 23, 86–102 (2012).

38. Ma, J. et al. TLR4 activation promotes podocyte injury and interstitial fibrosis in diabetic nephropathy. PLoS One 9, e97985 (2014).

39. Colhoun, H.M. & Marcovecchio, M.L. Biomarkers of diabetic kidney disease. Diabetologia 61, 996–1011 (2018).

40. McKnight, A.J. et al. Genetic polymorphisms in nitric oxide synthase 3 gene and implications for kidney disease: a meta-analysis. Am J Nephrol 32, 476–81 (2010).

41. Prabhakar, S.S. Role of nitric oxide in diabetic nephropathy. Semin Nephrol 24, 333–44 (2004).

42. Ko, Y.A. et al. Genetic-Variation-Driven Gene-Expression Changes Highlight Genes with Important Functions for Kidney Disease. Am J Hum Genet 100, 940–953 (2017).

43. Qiu, C. et al. Renal compartment-specific genetic variation analyses identify new pathways in chronic kidney disease. Nat Med 24, 1721–1731 (2018).

44. Chasman, D.I. et al. Integration of genome-wide association studies with biological knowledge identifies six novel genes related to kidney function. Hum Mol Genet 21, 5329–43 (2012).

45. Bernstein, B.E. et al. The NIH Roadmap Epigenomics Mapping Consortium. Nat Biotechnol 28, 1045–8 (2010).

46. Swan, E.J., Maxwell, A.P. & McKnight, A.J. Distinct methylation patterns in genes that affect mitochondrial function are associated with kidney disease in blood-derived DNA from individuals with Type 1 diabetes. Diabet Med 32, 1110–5 (2015).

47. Park, J. et al. Single-cell transcriptomics of the mouse kidney reveals potential cellular targets of kidney disease. Science 360, 758–763 (2018).

48. Kleppel, M.M., Fan, W., Cheong, H.I. & Michael, A.F. Evidence for separate networks of classical and novel basement membrane collagen. Characterization of alpha 3(IV)-alport antigen heterodimer. J Biol Chem 267, 4137–42 (1992).

49. Khoshnoodi, J., Pedchenko, V. & Hudson, B.G. Mammalian collagen IV. Microsc Res Tech 71, 357–70 (2008).

50. Kashtan, C.E. et al. Alport syndrome: a unified classification of genetic disorders of collagen IV alpha345: a position paper of the Alport Syndrome Classification Working Group. Kidney Int 93, 1045–1051 (2018).

51. Parkin, J.D. et al. Mapping structural landmarks, ligand binding sites, and missense mutations to the collagen IV heterotrimers predicts major functional domains, novel interactions, and variation in phenotypes in inherited diseases affecting basement membranes. Hum Mutat 32, 127–43 (2011).

52. Guan, M. et al. Association of kidney structure-related gene variants with type 2 diabetes-attributed end-stage kidney disease in African Americans. Hum Genet 135, 1251–1262 (2016).

53. Schofield, E.C. et al. CHiCP: a web-based tool for the integrative and interactive visualization of promoter capture Hi-C datasets. Bioinformatics 32, 2511–3 (2016).

54. Alves, F. et al. Distinct structural characteristics of discoidin I subfamily receptor tyrosine kinases and complementary expression in human cancer. Oncogene 10, 609–18 (1995).

55. Vogel, W., Gish, G.D., Alves, F. & Pawson, T. The discoidin domain receptor tyrosine kinases are activated by collagen. Mol Cell 1, 13–23 (1997).

56. Dorison, A. & Chantziantoniou, C. DDR1: A major player in renal diseases. Cell Adh Migr 12, 299–304 (2018).

57. Gross, O. et al. Loss of collagen-receptor DDR1 delays renal fibrosis in hereditary type IV collagen disease. Matrix Biol 29, 346–56 (2010).

58. Kerroch, M. et al. Genetic inhibition of discoidin domain receptor 1 protects mice against crescentic glomerulonephritis. FASEB J 26, 4079–91 (2012).

59. Mifsud, B. et al. Mapping long-range promoter contacts in human cells with high-resolution capture Hi-C. Nat Genet 47, 598–606 (2015).

60. Selman, L. & Hansen, S. Structure and function of collectin liver 1 (CL-L1) and collectin 11 (CL-11, CL-K1). Immunobiology 217, 851–63 (2012).

61. Hansen, S. et al. Collectin 11 (CL-11, CL-K1) is a MASP-1/3-associated plasma collectin with microbial-binding activity. J Immunol 185, 6096–104 (2010).

62. Farrar, C.A. et al. Collectin-11 detects stress-induced L-fucose pattern to trigger renal epithelial injury. J Clin Invest 126, 1911–25 (2016).

63. Rooryck, C. et al. Mutations in lectin complement pathway genes COLEC11 and MASP1 cause 3MC syndrome. Nat Genet 43, 197–203 (2011).

64. Walsh, D.W., Godson, C., Brazil, D.P. & Martin, F. Extracellular BMP-antagonist regulation in development and disease: tied up in knots. Trends Cell Biol 20, 244–56 (2010).

65. Zeisberg, M., Shah, A.A. & Kalluri, R. Bone morphogenic protein-7 induces mesenchymal to epithelial transition in adult renal fibroblasts and facilitates regeneration of injured kidney. J Biol Chem 280, 8094–100 (2005).

66. Vukicevic, S., Kopp, J.B., Luyten, F.P. & Sampath, T.K. Induction of nephrogenic mesenchyme by osteogenic protein 1 (bone morphogenetic protein 7). Proc Natl Acad Sci U S A 93, 9021–6 (1996).

67. Luo, G. et al. BMP-7 is an inducer of nephrogenesis, and is also required for eye development and skeletal patterning. Genes Dev 9, 2808–20 (1995).

68. Higgins, D.F. et al. BMP7-induced-Pten inhibits Akt and prevents renal fibrosis. Biochim Biophys Acta Mol Basis Dis 1863, 3095–3104 (2017).

69. Roxburgh, S.A. et al. Allelic depletion of grem1 attenuates diabetic kidney disease. Diabetes 58, 1641–50 (2009).

70. Dolan, V. et al. Expression of gremlin, a bone morphogenetic protein antagonist, in human diabetic nephropathy. Am J Kidney Dis 45, 1034–9 (2005).

71. McMahon, R. et al. IHG-2, a mesangial cell gene induced by high glucose, is human gremlin. Regulation by extracellular glucose concentration, cyclic mechanical strain, and transforming growth factor-beta1. J Biol Chem 275, 9901–4 (2000).

72. Afkarian, M. et al. Urinary excretion of RAS, BMP, and WNT pathway components in diabetic kidney disease. Physiol Rep 2, e12010 (2014).

73. Dabelea, D. et al. Effect of type 1 diabetes on the gender difference in coronary artery calcification: a role for insulin resistance? The Coronary Artery Calcification in Type 1 Diabetes (CACTI) Study. Diabetes 52, 2833–9 (2003).

74. Orchard, T.J. et al. Prevalence of complications in IDDM by sex and duration. Pittsburgh Epidemiology of Diabetes Complications Study II. Diabetes 39, 1116–24 (1990).

75. Marre, M. et al. Contribution of genetic polymorphism in the renin-angiotensin system to the development of renal complications in insulin-dependent diabetes: Genetique de la Nephropathie Diabetique (GENEDIAB) study group. J Clin Invest 99, 1585–95 (1997).

76. Hadjadj, S. et al. Different patterns of insulin resistance in relatives of type 1 diabetic patients with retinopathy or nephropathy: the Genesis France-Belgium Study. Diabetes Care 27, 2661–8 (2004).

77. Pezzolesi, M.G. et al. Genome-Wide Association Scan for Diabetic Nephropathy Susceptibility Genes in Type 1 Diabetes. Diabetes 58, 1403–1410 (2009).

78. Krolewski, A.S. Progressive renal decline: the new paradigm of diabetic nephropathy in type 1 diabetes. Diabetes Care 38, 954–62 (2015).

79. Sviklane, L. et al. Fatty liver index and hepatic steatosis index for prediction of non-alcoholic fatty liver disease in type 1 diabetes. J Gastroenterol Hepatol 33, 270–276 (2018).

80. Pop, A. et al. Insulin resistance is associated with all chronic complications in type 1 diabetes. J Diabetes 8, 220–8 (2016).

81. Akbar, T. et al. Cohort Profile: Scottish Diabetes Research Network Type 1 Bioresource Study (SDRNT1BIO). Int J Epidemiol 46, 796–796i (2017).

82. Rossing, P., Hougaard, P. & Parving, H.H. Risk factors for development of incipient and overt diabetic nephropathy in type 1 diabetic patients: a 10-year prospective observational study. Diabetes Care 25, 859–64 (2002).

83. Ma, J. et al. Genetic influences of the intercellular adhesion molecule 1 (ICAM-1) gene polymorphisms in development of Type 1 diabetes and diabetic nephropathy. Diabet Med 23, 1093–9 (2006).

84. Mollsten, A. et al. The effect of polymorphisms in the renin-angiotensin-aldosterone system on diabetic nephropathy risk. J Diabetes Complications 22, 377–83 (2008).

85. Klein, R., Klein, B.E., Moss, S.E. & Cruickshanks, K.J. The Wisconsin Epidemiologic Study of Diabetic Retinopathy: XVII. The 14-year incidence and progression of diabetic retinopathy and associated risk factors in type 1 diabetes. Ophthalmology 105, 1801–15 (1998).

86. Cohen, C.D., Frach, K., Schlondorff, D. & Kretzler, M. Quantitative gene expression analysis in renal biopsies: a novel protocol for a high-throughput multicenter application. Kidney Int 61, 133–40 (2002).

87. Berthier, C.C. et al. Enhanced expression of Janus kinase-signal transducer and activator of transcription pathway members in human diabetic nephropathy. Diabetes 58, 469–77 (2009).

88. Schmid, H. et al. Modular activation of nuclear factor-kappaB transcriptional programs in human diabetic nephropathy. Diabetes 55, 2993–3003 (2006).

89. Todd, J.N. et al. Genetic Evidence for a Causal Role of Obesity in Diabetic Kidney Disease. Diabetes 64, 4238–46 (2015).

90. Irizarry, R.A. et al. Exploration, normalization, and summaries of high density oligonucleotide array probe level data. Biostatistics 4, 249–64 (2003).

91. Johnson, W.E., Li, C. & Rabinovic, A. Adjusting batch effects in microarray expression data using empirical Bayes methods. Biostatistics 8, 118–27 (2007).

