## Supplement for "Genome-wide association study of diabetic kidney disease highlights biology involved in renal basement membrane collagen"

**SUPPLEMENTAL INFORMATION**

Figure S9. Functional annotation of *TAMM41*. 41
Supplemental Methods: Cohort descriptions 42
References 49

**Table S1. Cohorts contributing to analyses.**
The table can be found in a separate excel sheet, Supplemental_table_S1.xlsx

**Table S2. Characteristics of RASS participants.** Categorical variables display counts and percentage. Continuous values are mean ± standard deviation.

| **Variables (Total N = 253)** | **Freq(%)/Mean±SD** |
|---|---|
| Sex - Female | 134 (53%) |
| Age (years) | 30 ± 10 |
| T1D duration (years) | 11 ± 5 |
| Within-person mean HbA1c (%)<br>(mmol/mol) | 8.6 ± 1.4<br>70 ± 15 |
| Mean GBMW (nm) | 480 ± 88 |
| **rs55703767** – GG<br>GT<br>TT | 163 (64%)<br>80 (32%)<br>10 (4%) |

**Table S3. Multivariate analysis of association between rs55703767 and GBM width**

| Variables | | Adjusted model | | | Fully adjusted model* | |
|---|---|---|---|---|---|---|
| | | **Effect (SE)** | **P** | | **Effect (SE)** | **P** |
| rs55703767 (T allele)[¶] | | -22.8 (8.2) | 0.006 | | -19.7 (8.2)[¶] | 0.0172 |
| Females (vs males) | | -48.4 (9.3) | <.0001 | | -50.4 (9.3) | <.0001 |
| Age at baseline (yrs) | | -2.4 (0.5) | <.0001 | | -2.4 (0.5) | <.0001 |
| Diabetes duration (yrs) | | 3.8 (1.0) | 0.0002 | | 3.8 (1.0) | 0.0002 |
| Mean HbA1c (%) | | 27.2 (3.3) | <.0001 | | 27.4 (3.3) | <.0001 |
| Treatments | Placebo | - | - | | Reference | |
| | Enalapril | - | - | | -6.9 (11.2) | 0.538 |
| | Losartan | - | - | | 1.4 (10.9) | 0.896 |
| Centres | Montreal | - | - | | Reference | |
| | Toronto | - | - | | 0.8 (12.8) | 0.952 |
| | Minnesota | - | - | | 18.9 (13.7) | 0.169 |

* Fully adjusted model also included 3 principal components for population structure within Europeans.
[¶] SNP genotypes modelled as additive genetic effects.

**Table S4: Look-up of the lead loci in GWAS on eGFR in the general population** (Gorski et al., 2017)**.**

| | | Meta-analysis results | | | | | | | GWAS on eGFR (Gorski 2017) | | | | |
|---|---|---|---|---|---|---|---|---|---|---|---|---|---|
| Nearest Gene | SNP | EA | NEA | EAF | SE | OR | $P_{min}$ | $P_{Full}$ | EAF | β | SE | P | N |
| *COL4A3* | rs55703767 | T | G | 0.206 | 0.03 | 0.79 | **5.34E-12** | **8.19E-11** | 0.142 | 0.002 | 0.0013 | 0.132 | 110517 |
| *COL4A3* | rs55703767 | T | G | 0.209 | 0.04 | 0.79 | **9.28E-09** | **9.38E-09** | 0.142 | 0.002 | 0.0013 | 0.132 | 110517 |
| *COL4A3* | rs55703767 | T | G | 0.205 | 0.03 | 0.84 | **3.88E-10** | **9.68E-09** | 0.142 | 0.002 | 0.0013 | 0.132 | 110517 |
| *COL4A3* | rs55703767 | T | G | 0.208 | 0.04 | 0.77 | **5.30E-09** | **3.77E-08** | 0.142 | 0.002 | 0.0013 | 0.132 | 110517 |
| *PRNCR1* | rs551191707 | CA | C | 0.122 | 0.1 | 1.7 | **4.39E-08** | 3.15E-06 | | | | | |
| *STXBP6* | rs61983410 | T | C | 0.787 | 0.04 | 1.26 | 9.84E-08 | **3.06E-08** | 0.841 | -0.001 | 0.0012 | 0.336 | 110516 |
| *COLEC11* | rs12615970 | A | G | 0.867 | 0.05 | 1.31 | **9.43E-09** | 1.60E-07 | | | | | |
| *LINC01266* | rs115061173 | A | T | 0.014 | 0.41 | 9.39 | **4.07E-08** | 4.08E-05 | | | | | |
| *SNCAIP* | rs149641852 | T | G | 0.012 | 0.39 | 9.03 | **1.37E-08** | --- | 0.009 | 0.002 | 0.0042 | 0.643 | 109257 |
| *PAPLN* | rs113554206 | A | G | 0.012 | 0.3 | 4.62 | 5.39E-07 | **8.46E-09** | 0.007 | -0.005 | 0.0064 | 0.408 | 95870 |
| *STAC* | rs116216059 | A | C | 0.016 | 0.38 | 8.76 | **1.37E-08** | 1.41E-04 | 0.006 | 3.00E-04 | 0.0043 | 0.953 | 108165 |
| *HAND2-AS1* | rs145681168 | A | G | 0.986 | 0.33 | 0.18 | 2.06E-07 | **5.40E-09** | 0.993 | 0.003 | 0.0067 | 0.612 | 64752 |
| *TAMM41* | rs142823282 | A | G | 0.983 | 0.31 | 0.15 | **8.32E-10** | **1.13E-11** | | | | | |
| *VARS2* | rs118124843 | T | C | 0.011 | 0.24 | 3.78 | **4.42E-08** | **3.37E-08** | 0.031 | 0.011 | 0.0055 | *0.040* | 58794 |
| *MUC7* | rs191449639 | A | T | 0.005 | 0.61 | 32.46 | **1.32E-08** | **2.09E-08** | | | | | |
| *MBLAC1* | rs77273076 | T | C | 0.008 | 0.39 | 9.12 | **1.04E-08** | 2.28E-07 | 0.007 | 0.006 | 0.0051 | 0.236 | 108694 |
| *BMP7* | rs144434404 | T | C | 0.011 | 0.32 | 6.75 | **2.67E-09** | **4.65E-09** | 0.004 | 1.00E-04 | 0.0072 | 0.993 | 91428 |
| *PLEKHA7* | rs183937294 | T | G | 0.993 | 0.5 | 0.06 | **1.65E-08** | 2.10E-06 | | | | | |
| | rs185299109 | T | C | 0.007 | 0.53 | 20.7 | **1.28E-08** | 4.99E-07 | | | | | |

EA: Effect allele. Positive odds ratio indicates that EA is associated with higher risk; positive beta indicates that EA is associated with higher eGFR, i.e. lower renal risk.

### Table S5: Look-up of the lead loci in GWAS in the SUMMIT consortium.

| SNP | Chr:pos | EA | NEA | EAF | Notable gene(s) | Phenotype | N | OR | P-value |
|---|---|---|---|---|---|---|---|---|---|
| rs55703767 | 2:228121101 | T | G | 0.211 | *COL4A3* | **DN** | **5190** | **0.911** | **0.08** |
| | | | | 0.213 | | **CKD+DN** | **2243** | **0.867** | **0.09** |
| rs145681168 | 4:174500806 | G | A | 0.017 | *HAND2-AS1* | **Micro** | **3477** | **1.034** | **0.97** |
| rs149641852 | 5:121774582 | T | G | 0.018 | *SNCAIP* | **CKD** | **4676** | **1.032** | **0.30** |
| rs118124843 | 6:30887465 | T | C | 0.018 | *DDR1, VARS2* | **Micro** | **2439** | **1.137** | **0.63** |
| rs77273076 | 7:99728546 | T | C | 0.014 | *MBLAC1* | Micro | 3252 | 0.866 | 0.48 |
| rs61983410 | 14:26004712 | T | C | 0.184 | *STXBP6* | **Micro** | **3760** | **0.990** | **0.58** |
| rs144434404 | 20:55837263 | T | C | 0.011 | *BMP7* | **Micro** | **2439** | **1.100** | **0.78** |

Chr, chromosome; pos, position; EA: Effect allele; EAF, effect allele frequency; OR, odds ratio.

### Table S6: Association at lead loci stratified by HbA1c <7.5%.

| | | | | | ALL | | | | HbA1c < 7.5% | | | HbA1c >= 7.5% | | | |
|---|---|---|---|---|---|---|---|---|---|---|---|---|---|---|---|
| Locus | SNP | Pheno | EA | NEA | N | MAF | P | INFO | N (case/ctrl) | P | OR (95% CI) | N (case/ctrl) | P | OR (95% CI) | P_HET |
| *COL4A3* | rs55703767 | MACROESRD | G | T | 3611 | 0.19 | **2.16E-03** | 1.00 | 1165 (499/666) | 0.659 | 0.95 (0.76;1.19) | 2495 (884/1611) | **9.55E-04** | 0.77 (0.66;0.9) | 0.132 |
| *COL4A3* | rs55703767 | MACRO | G | T | 2803 | 0.19 | 0.06 | 1.00 | 837 (164/673) | 0.663 | 1.08 (0.78;1.49) | 2006 (373/1633) | **6.63E-03** | 0.75 (0.61;0.92) | 0.068 |
| *COL4A3* | rs55703767 | ALLvCTRL | G | T | 4271 | 0.19 | **7.04E-03** | 1.00 | 1344 (692/652) | 0.870 | 0.98 (0.8;1.2) | 2977 (1391/1586) | **1.76E-03** | 0.81 (0.71;0.92) | 0.114 |
| *COL4A3* | rs55703767 | CKDDN | G | T | 3059 | 0.19 | **1.17E-02** | 1.00 | 984 (379/605) | 0.973 | 1 (0.78;1.28) | 2102 (624/1478) | **7.90E-03** | 0.79 (0.67;0.94) | 0.136 |
| *PRNCR1* | rs551191707 | ESRDvMACRO | C | CA | 1371 | 0.14 | **2.50E-03** | 0.81 | 498 (340/158) | **1.92E-02** | 1.71 (1.09;2.67) | 885 (524/361) | **4.79E-02** | 1.38 (1;1.91) | 0.453 |
| *STXBP6* | rs61983410 | MICRO | T | C | 2976 | 0.23 | **3.75E-03** | 0.93 | 863 (195/668) | **1.34E-02** | 0.69 (0.52;0.93) | 2155 (526/1629) | 0.067 | 0.85 (0.71;1.01) | 0.248 |
| *COLEC11* | rs12615970 | CKD | A | G | 4264 | 0.14 | **3.13E-03** | 0.82 | 1432 (531/901) | 0.086 | 0.81 (0.63;1.03) | 3014 (833/2181) | **1.62E-02** | 0.8 (0.66;0.96) | 0.949 |
| *LINC01266* | rs115061173 | ESRD | T | A | 3119 | 0.00 | **1.89E-02** | 0.36 | 1012 (340/672) | 0.284 | 2.63 (0.45;15.36) | 2156 (524/1632) | 0.085 | 5.98 (0.78;45.9) | 0.550 |
| *SNCAIP* | rs149641852 | CKDEXTREMES | G | T | 3907 | 0.01 | **3.04E-03** | 0.33 | 1323 (415/908) | 0.559 | 1.53 (0.37;6.35) | 2765 (559/2206) | **6.27E-04** | 10.78 (2.76;42.09) | 0.052 |
| *PAPLN* | rs113554206 | MACRO | G | A | 2803 | 0.00 | 0.32 | 0.34 | 837 (164/673) | 0.793 | 0.39 (0;417.59) | 2006 (373/1633) | 0.114 | 21.87 (0.48;999.19) | 0.322 |
| *STAC* | rs116216059 | ESRDvALL | C | A | 4272 | 0.01 | 0.48 | 0.67 | 1340 (340/1000) | 0.867 | 0.88 (0.2;3.89) | 2984 (524/2460) | 0.453 | 0.67 (0.23;1.93) | 0.764 |
| *HAND2-AS1* | rs145681168 | MICRO | A | G | 2976 | 0.01 | 0.50 | 0.48 | 863 (195/668) | 0.509 | 3.48 (0.09;141.38) | 2155 (526/1629) | 0.395 | 0.61 (0.19;1.91) | 0.378 |
| *TAMM41* | rs142823282 | MICRO | A | G | 2976 | 0.00 | 0.93 | 0.15 | | | | 2155 (526/1629) | 0.886 | 1.19 (0.11;13.33) | NA |
| *VARS2* | rs118124843 | MICRO | C | T | 2976 | 0.01 | 0.93 | 1.00 | 863 (195/668) | 0.533 | 0.59 (0.11;3.16) | 2155 (526/1629) | 0.769 | 1.15 (0.46;2.85) | 0.492 |
| *MUC7* | rs191449639 | MACROESRD | T | A | 3611 | 0.00 | 0.09 | 0.28 | 1165 (499/666) | 0.487 | 2.17 (0.24;19.36) | 2495 (884/1611) | 0.246 | 3.58 (0.42;30.94) | 0.749 |
| *MBLAC1* | rs77273076 | MICRO | C | T | 2976 | 0.01 | **1.36E-04** | 0.37 | 863 (195/668) | **3.98E-03** | 168.26 (5.14;5507.78) | 2155 (526/1629) | **1.68E-03** | 11.25 (2.48;50.97) | 0.163 |
| *BMP7* | rs144434404 | MICRO | C | T | 2976 | 0.01 | 0.57 | 0.67 | 863 (195/668) | 0.407 | 0.49 (0.09;2.61) | 2155 (526/1629) | 0.911 | 0.94 (0.3;2.91) | 0.534 |
| *PLEKHA7* | rs183937294 | MICRO | T | G | 2976 | 0.00 | 0.22 | 0.26 | | | | 2155 (526/1629) | 0.288 | 3.79 (0.32;44.52) | NA |
| | rs185299109 | CKD | C | T | 4264 | 0.00 | 0.68 | 0.32 | 1432 (531/901) | 0.147 | 0.12 (0.01;2.09) | 3014 (833/2181) | 0.956 | 0.96 (0.21;4.45) | 0.212 |

Association stratified by HbA1c in the FinnDiane study. *P*-values <0.05 are given with scientific notation and bold. Lines with gray text had minor allele count (MAC)<10 in cases and/or controls and did not contribute to the meta-analysis.

### Table S7. Association of rs55703767 with DN in DCCT/EDIC subgroups.

| Cohort | Treatment Group | DN % | MAF | Last measure | | Time to Event | |
|---|---|---|---|---|---|---|---|
| | | | | OR (95%CI) | P value | HR (95%CI) | P value |
| Primary Prevention | Intensive | 3% | 0.22 | 2.86 (0.4-22) | 0.32 | 0.91 (0.2-4.0) | 0.90 |

| | | | | | | | |
|---|---|---|---|---|---|---|---|
| (diabetes dur 1-5 yrs) | Conventional | 10% | 0.21 | 0.67 (0.3-1.4) | 0.31 | 0.66 (0.32-1.33) | 0.24 |
| Secondary Intervention | Intensive | 5% | 0.20 | 0.86 (0.3-2.6) | 0.79 | 0.65 (0.22-1.9) | 0.43 |
| (diabetes dur 1-15 yrs) | Conventional | 13% | 0.22 | 0.18 (0.1-0.5) | 0.003 | 0.30 (0.13-0.68) | 0.004 |

OR=Odds Ratio for last measure, HR=Hazard Ratio for time to event phenotype

**Table S8. Significant (*P*<0.05/18,222 genes tested = $2.74 \times 10^{-6}$) gene level associations with diabetic kidney disease in MAGMA.**

| Gene | Phenotype | Model | Window | Number of SNPs | Total Sample Size | MAGMA P-value | PASCAL P-value |
|---|---|---|---|---|---|---|---|
| *SLC46A2* | All vs. ctrl | Min | nowindow | 66 | 17817 | $6.74 \times 10^{-7}$ | $1.57 \times 10^{-5}$ |
| | | | 5kbwindow | 93 | 17832 | $7.38 \times 10^{-7}$ | |
| | | Full | nowindow | 64 | 16821 | $8.13 \times 10^{-7}$ | $6.93 \times 10^{-5}$ |
| | | | 5kbwindow | 90 | 16855 | $1.03 \times 10^{-6}$ | |
| *SFXN4* | Macro | Full | nowindow | 69 | 11953 | $3.98 \times 10^{-7}$ | $1.45 \times 10^{-4}$ |
| | | | 5kbwindow | 86 | 11857 | $1.65 \times 10^{-7}$ | |
| *COL20A1* | Ckdextreme | Min | nowindow | 111 | 11165 | $2.47 \times 10^{-6}$ | $7.88 \times 10^{-5}$ |
| | | | 5kbwindow | 137 | 11603 | $2.01 \times 10^{-6}$ | |
| | | Full | nowindow | 110 | 8533 | $6.65 \times 10^{-7}$ | $4.47 \times 10^{-5}$ |
| | | | 5kbwindow | 136 | 9044 | $5.77 \times 10^{-7}$ | |
| | ESRD vs. All | Min | nowindow | 111 | 12063 | $1.34 \times 10^{-6}$ | $3.76 \times 10^{-5}$ |
| | | | 5kbwindow | 137 | 12362 | $1.04 \times 10^{-6}$ | |
| | | Full | nowindow | 110 | 8638 | $1.12 \times 10^{-6}$ | $5.81 \times 10^{-5}$ |
| | | | 5kbwindow | 136 | 9045 | $9.53 \times 10^{-7}$ | |
| *GLT6D1* | ESRD vs. Macro | min | 5kbwindow | 96 | 4248 | $1.49 \times 10^{-6}$ | $2.15 \times 10^{-5}$ |
| *SNX30* | All vs. ctrl | min | 5kbwindow | 434 | 18249 | $2.49 \times 10^{-6}$ | $1.05 \times 10^{-5}$ |

**Table S9. Top nominally significant gene level associations (P < $1.0 \times 10^{-5}$) with diabetic kidney disease in PASCAL.**

| Gene | Phenotype | Model | Number of SNPs | PASCAL P-value |
|---|---|---|---|---|
| *INIP* | All vs. ctrl | Min | 248 | $1.99 \times 10^{-6}$ |
| | | Full | 248 | $5.54 \times 10^{-6}$ |
| *LCN9* | ESRD vs. macro | Min | 301 | $5.25 \times 10^{-6}$ |
| *CBX8* | DN | Min | 119 | $8.47 \times 10^{-6}$ |

**Table S10: Significant gene set and pathway analysis results.** Significantly enriched gene sets identified from at least one of the following methods: MAGENTA (FDR < 0.05, MAGMA (P<0.05 empirical permutation multiple testing correction), PASCAL (P<0.05/1,078 gene sets tested = 4.64 × 10-5), and DEPICT (FDR < 0.01).

| Gene set | Gene set database | Phenotype | Model | Method |
|---|---|---|---|---|
| negative regulators of RIG I MDA5 signaling | REACTOME | ESRD vs. Macro | Full | MAGMA |
| Platelet aggregation plug formation | REACTOME | Micro | Min | MAGMA |
| negative regulators of RIG I MDA5 signaling | REACTOME | ESRD vs. Macro | Full | PASCAL |
| RIG I MDA5 mediated induction of IFN alpha beta pathways | REACTOME | ESRD vs. Macro | Full | PASCAL |
| TRAF3 dependent IRF activation pathway | REACTOME | ESRD vs. Macro | Full | PASCAL |
| TRAF6 mediated IRF activation | REACTOME | ESRD vs. Macro | Full | PASCAL |
| Nitric Oxide Signaling in the Cardiovascular System | Ingenuity | ESRD vs. ctrl | Min | MAGENTA |
| Nicotinic acetylcholine receptor signaling pathway | Panther | ESRD vs. non-ESRD | Min | MAGENTA |
| ACTIVATED TLR4 SIGNALLING | REACTOME | All vs. ctrl | Min | MAGENTA |
| Other lipid, fatty acid and steroid metabolism | PANTHER BIOLOGICAL PROCESS | CKD | Min | MAGENTA |
| DNA degradation | PANTHER BIOLOGICAL PROCESS | CKD | Min | MAGENTA |
| Tumor necrosis factor family member | PANTHER MOLECULAR FUNCTION | CKD-extreme | Min | MAGENTA |
| TUFM (Tu Translation Elongation Factor, Mitochondrial) PPI subnetwork | InWeb protein-protein interaction database | DN | Min | DEPICT |

### Table S11. eQTL associations and chromatin conformation interactions for the lead SNPs.

| SNP | Chr:pos | EA | NEA | EAF | Notable gene(s) | eQTL | | | | PC-HiC | |
|---|---|---|---|---|---|---|---|---|---|---|---|
| | | | | | | GENE | P | HIGH A | Tissue | *Gene* | *Score (Tissue)* |
| rs12615970 | 2:3745215 | G | A | 0.133 | *COLEC11* (B); | | | | | ***ALLC*** | 10.42 (GM12878); |
| | | | | | *ALLC (G)* | | | | | ***COLEC11,*** *AC010907.2* | 9.67 (GM12878); |
| | | | | | | | | | | ***ADI1,*** *AC142528.1* | 8.75 (GM12878); |
| | | | | | | | | | | *RP13-512J5.1* | 8.58 (GM12878); |
| | | | | | | | | | | *RPS7* | 8.13 (GM12878); |
| rs55703767 | 2:228121101 | T | G | 0.206 | *COL4A3* (M, B, N) | ***MFF*** | $5.63\times10^{-38}$ | T | blood | ***COL4A3, COL4A4*** | 8.89 (GM12878); |
| | | | | | | ***MFF*** | $9.0\times10^{-8}$ | T | Cells - Transformed fibroblasts | ***IRS****, RP11-395N3.2* | 9.36 (GM12878); |
| | | | | | | ***TM4SF20*** | $2.2\times10^{-7}$ | T | Cells - Transformed fibroblasts | | |
| rs115061173 | 3:926345 | A | T | 0.014 | *LINC01266* (N) | | | | | | |
| rs142823282 | 3:11910635 | G | A | 0.011 | *TAMM41* (N, B) | ***PPARG*** | $4.6\times10^{-7}$ | G | Colon - Sigmoid | ***TAMM41*** | 10.65 (GM12878); |
| rs116216059 | 3:36566312 | A | C | 0.016 | *STAC* (G) | | | | | ***DCLK3*** | 8.8 (GM12878); |
| | | | | | | | | | | ***STAC*** | 10.87 (GM12878); |
| rs191449639 | 4:71358776 | A | T | 0.005 | *MUC7* (N) | | | | | | |
| rs145681168 | 4:174500806 | G | A | 0.014 | *HAND2-AS1*(G, B) | | | | | ***HAND2****, HAND2-AS1* | 10.49 (GM12878); |
| rs149641852 | **5:121774582** | **T** | **G** | **0.012** | ***SNCAIP* (G)** | | | | | ***SNX24*** | 9.2 (GM12878); |
| | | | | | | | | | | *snoU13* | 8.93 (GM12878); |
| | | | | | | | | | | ***SNCAIP,*** *CTD-2544H17.2* | 9.69 (GM12878); |
| | | | | | | | | | | *CTD-2280E9.1* | 10.81 (GM12878); |
| rs118124843 | 6:30887465 | T | C | 0.011 | *DDR1* (B); | ***HLA-C*** | $1.00\times10^{-18}$ | C | eQTLgen blood | ***PSORS1C1*** | 12.3 (Endothelial Precursors); 12.3 (Endothelial Precursors); 7.84 (Megacaryocytes); 5.63 (Pancreatic islets); |
| | | | | | *VARS2* (G) | | | | | ***DPCR1, HCG21*** | 11.49 (GM12878); |
| | | | | | | ***HLA-U*** | $3.56\times10^{-10}$ | T | eQTLgen blood | *DDR1-AS1,* ***DDR1*** | 10.97 (GM12878); |
| | | | | | | ***PSORS1C3*** | $4.13\times10^{-9}$ | T | eQTLgen blood | *RNU6-1133P* | 7.27 (Macrophages M2); |
| | | | | | | ***NCR3*** | $9.35\times10^{-6}$ | C | eQTLgen blood | *RN7SL175P,* ***DDR1, GTF2H4, VARS2*** | 7.07 (Endothelial Precursors); 7.07 (Endothelial Precursors); 6.95 (Cardiomyocytes); 6.89 (Pancreatic |

| SNP | Chr:pos | EA | NEA | EAF | Notable gene(s) | eQTL | | | | PC-HiC | |
|---|---|---|---|---|---|---|---|---|---|---|---|
| | | | | | | GENE | P | HIGH A | Tissue | *Gene* | *Score (Tissue)* |
| | | | | | | | | | | | islets); 5.43 (Megacaryocytes); 5.09 (Macrophages M1); |
| | | | | | | ***HCG22*** | $1.5\times10^{-5}$ | T | eQTLgen blood | *C6orf15* | 6.95 (Macrophages M1); 6.95 (Macrophages M1); 6.18 (Macrophages M0); |
| | | | | | | ***VARS2*** | $1.71\times10^{-5}$ | C | eQTLgen blood | | |
| | | | | | | ***GTF2H4*** | $9.70\times10^{-7}$ | T | Esophagus - Gastroesophageal Junction | | |
| | | | | | | ***POU5F1*** | $3.3\times10^{-5}$ | T | Esophagus - Gastroesophageal Junction | | |
| | | | | | | ***PSORS1C3*** | $5.6\times10^{-5}$ | T | Esophagus - Gastroesophageal Junction | | |
| | | | | | | *C6orf48* | $1\times10^{-4}$ | C | Nerve - Tibial | | |
| rs77273076 | 7:99728546 | T | C | 0.008 | *MBLAC1* (N, B) | ***CNPY4*** | $1.17\times10^{-7}$ | C | eQTLgen blood | ***MBLAC1,*** *AC073842.19, RP11-506M12.1* | ***NA (withn the same fragment);*** |
| | | | | | | ***AP4M1*** | $1.04\times10^{-5}$ | C | eQTLgen blood | ***LAMTOR4, GAL3ST4, GPC2,*** *C7orf43, MIR4658* | 14.82 (CD34); 14.82 (CD34); 14.56 (GM12878); |
| | | | | | | ***ZSCAN21*** | $1.29\times10^{-5}$ | C | eQTLgen blood | ***LAMTOR4*** | 14.14 (CD34); 14.14 (CD34); 13.1 (GM12878); |
| | | | | | | | | | | ***GATS, STAG3, PVRIG,*** *AC005071.1* | 14.11 (CD34); 14.11 (CD34); 13.68 (GM12878); |
| | | | | | | | | | | ***MCM7, AP4M1*** | 14.11 (CD34); 14.11 (CD34); 13.68 (GM12878); |
| | | | | | | | | | | ***STAG3, GPC2*** | 13.75 (CD34); 13.75 (CD34); 13.36 (GM12878); |
| | | | | | | | | | | ***MCM7, COPS6,*** *MIR93, MIR106B, MIR25* | 13.69 (CD34); 13.69 (CD34); 13.67 (GM12878); |
| | | | | | | | | | | ***CNPY4, TAF6*** | 13.37 (GM12878); 13.37 (GM12878); 13.14 (CD34); |
| | | | | | | | | | | ***ZKSCAN1*** | 13.23 (GM12878); 13.23 (GM12878); 11.87 (CD34); |
| | | | | | | | | | | ***ZSCAN21*** | 13.14 (GM12878); |
| | | | | | | | | | | ***ZCWPW1, MEPCE*** | 12.72 (GM12878); 12.72 (GM12878); 11.48 (CD34); |
| | | | | | | | | | | ***PILRA*** | 12.65 (GM12878); |

| SNP | Chr:pos | EA | NEA | EAF | Notable gene(s) | eQTL | | | | PC-HiC | |
|---|---|---|---|---|---|---|---|---|---|---|---|
| | | | | | | GENE | P | HIGH A | Tissue | *Gene* | *Score (Tissue)* |
| | | | | | | | | | | ***TRIM4*** | 12.64 (GM12878); |
| | | | | | | | | | | ***ZCWPW1*** | 12.61 (GM12878); |
| | | | | | | | | | | ***SAP25, FBXO24, LRCH4,*** *RP11-44M6.3* | 12.59 (GM12878); |
| | | | | | | | | | | ***PILRB, PVRIG2P,*** *STAG3L5P-PVRIG2P-PILRB,* | 12.38 (GM12878); 12.38 (GM12878); 5.94 (Naive B); 5.02 (Total B); |
| | | | | | | | | | | ***TSC22D4, NYAP1****, AC092849.1, RN7SL161P, C7orf61* | 12.21 (GM12878); |
| | | | | | | | | | | *RP11-758P17.2, PPP1R35, RP11-758P17.3* | 12.02 (GM12878); 12.02 (GM12878); 11.55 (CD34); |
| | | | | | | | | | | ***AZGP1, AZGP1P1*** | 11.3 (Neutrophils); 11.3 (Neutrophils); 6.07 (Macrophages M2); 5.65 (Total CD4 MF); 5.65 (Total CD4 MF); 5.1 (Total CD4 Activated); |
| | | | | | | | | | | ***BUD31,*** *snoU13* | 11.03 (GM12878); |
| | | | | | | | | | | ***ZNF3*** | 10.83 (GM12878); |
| | | | | | | | | | | ***ZCWPW1*** | 8.39 (Naive B); 8.39 (Naive B); 5.52 (Total CD4 Activated); |
| | | | | | | | | | | ***PMS2P1*** | 5.16 (Foetal Thymus); |
| rs551191707 | 8:128100029 | CA | C | 0.122 | *PRNCR1* ( N) | *-NONE-* | | | | | |
| rs183937294 | 11:16937846 | G | T | 0.007 | *PLEKHA7* (G) | *-NONE-* | | | | *RNU6-585, PRP11-466H18.1* | 10.77 (cd34); 10.77 (cd34); 9.2 (GM12878); |
| | | | | | | | | | | *AC116533.1, SNORD14B, SNORD14A, rps13* | 10.19 (GM12878); |
| | | | | | | | | | | ***PLEKHA7, OR7E14P*** | 9.49 (GM12878); |
| | | | | | | | | | | ***SOX6,*** *C11orf58* | 9.15 (GM12878); |
| | | | | | | | | | | ***SERGEF,*** *RP1-59M18.2* | 7.56 (GM12878); |
| | | | | | | | | | | ***OTOG*** | 5.89 (Total CD8); |
| | | | | | | | | | | ***USH1C*** | 5.07 (Naive CD8); |

| SNP | Chr:pos | EA | NEA | EAF | Notable gene(s) | eQTL | | | | PC-HiC | |
|---|---|---|---|---|---|---|---|---|---|---|---|
| | | | | | | GENE | P | HIGH A | Tissue | *Gene* | *Score (Tissue)* |
| rs61983410 | 14:26004712 | T | C | 0.213 | *STXBP6* (N) | *-NONE-* | | | | *SNORD37* | 10.97 (GM12878); |
| rs113554206 | 14:73740250 | A | G | 0.012 | *PAPLN* (G) | *-NONE-* | | | | *RP4-647C14.3* | ***NA (within the same fragment);*** |
| | | | | | | | | | | ***NUMB*** | 21.42 (Endotheial precursors); 21.42 (Endotheial precursors); 17.38 (Pancreatic islets); 12.32 (Megacaryocytes); 11.56 (CD34); 9.65 (Neutrophils); 9.61 (Naive B); 9.06 (Total B); 7.55 (cardiomyocytes); 5.33 (Naive CD4); 5.11 (Naive CD8); |
| | | | | | | | | | | ***PAPLN,*** *RNU6-419P, RP4-647C14.2* | 13.67 (CD34); 13.67 (CD34); 13.24 (GM12878); |
| | | | | | | | | | | ***PAPLN*** | 13.24 (CD34); 13.24 (CD34); 12.47 (GM12878); |
| | | | | | | | | | | ***PSEN1*** | 12.08 (GM12878); 12.08 (GM12878); 5.4 (cardiomyocytes); |
| | | | | | | | | | | ***HEATR4*** | 12.05 (Endotheial precursors); 12.05 (Endotheial precursors); 10.98 (Megacaryocytes); 10.45 (GM12878); 10.37 (CD34); 9.92 (Neutrophils); 8.84 (Pancreatic islets); 7.98 (Monocytes); 7.3 (Total B); 7.02 (Naive B); 6.6 (cardiomyocytes); 5.81 (Erythroblasts); 5.53 (Total CD4 Activated); |
| | | | | | | | | | | *RP1-240K6.3* | 11.84 (GM12878); 11.84 (GM12878); 11.15 (CD34); 7.49 (Endotheial precursors); 5.62 (Pancreatic islets); 5.34 (Total B); 5.02 (Megacaryocytes); |
| | | | | | | | | | | ***DNAL1,*** *RNU6-240P* | 11.07 (GM12878); |
| | | | | | | | | | | ***PSEN1*** | 10.96 (Endotheial precursors); 10.96 (Endotheial precursors); 10.56 (CD34); |
| | | | | | | | | | | ***PNMA1*** | 10.9 (GM12878); |
| | | | | | | | | | | *RP3-414A15.2* | 10.64 (GM12878); 10.64 (GM12878); 7.24 (Monocytes); 5.99 (Neutrophils); |
| | | | | | | | | | | ***ZFYVE1*** | 10.53 (GM12878); |
| | | | | | | | | | | ***PTGR2,*** *Y_RNA, RP5-1021I20.4* | 10.42 (CD34); 10.42 (CD34); 10.16 (GM12878); |
| | | | | | | | | | | *RP4-693M11.3* | 10.33 (GM12878); |
| | | | | | | | | | | *RP4-687K1.2* | 9.96 (GM12878); |

| SNP | Chr:pos | EA | NE A | EAF | Notable gene(s) | eQTL | | | | PC-HiC | |
|---|---|---|---|---|---|---|---|---|---|---|---|
| | | | | | | GENE | P | HIGH A | Tissue | *Gene* | *Score (Tissue)* |
| | | | | | | | | | | ***HEATR4,*** *C14orf169, AC005280.1* | 9.52 (GM12878); 9.52 (GM12878); 5.35 (Pancreatic islets); |
| | | | | | | | | | | ***RBM25*** | 9.34 (GM12878); |
| | | | | | | | | | | *RP3-414A15.10* | 9.32 (GM12878); |
| | | | | | | | | | | ***ELMSAN1*** | 9.12 (GM12878); |
| | | | | | | | | | | ***CCDC176*** | 8.85 (GM12878); |
| | | | | | | | | | | ***RBM25,*** *RP11-109N23.5* | 8.74 (GM12878); |
| | | | | | | | | | | ***ACOT6*** | 8.74 (GM12878); |
| | | | | | | | | | | ***DNAL1*** | 8.7 (GM12878); |
| | | | | | | | | | | ***FAM161B****, RP5-1021I20.5* | 5.58 (Total CD8); |
| rs185299109 | 18:1811108 | T | C | 0.007 | | *-NONE-* | | | | *-NONE-* | |
| **rs144434404** | **20:55837263** | **T** | **C** | **0.011** | ***BMP7*** **(G, B)** | *-NONE-* | | | | *-NONE-* | |

Notable Genes: based on genetic findings (G), (B), (N), (M); eQTL associations were searched from GTEX and eQTLgen (cis-eQTL) data sets. HIGH A: allele associated with higher gene expression levels. Promoter Capture Hi-C (PCHi-C) data: searched from www.chicp.org (date accessed: 1.12.2018; Schofield EC, Carver T, Achuthan P, Freire-Pritchett P, Spivakov M, Todd JA, Burren OS. CHiCP: a web-based tool for the integrative and interactive visualization of promoter capture Hi-C datasets. Bioinformatics. (2016) 15:32(16):2511-3), including 16 primary blood cell types and foetal thymocytes (Javierre et al.), CD34 and GM12878 cell line (Mifsud et al.), pancreatic isles (Miguel-Escalada et al,), and hESC derived cardiomyocytes (Choy et al.). Score: CHiCAGO score, values >5 were considered significant and listed. Protein coding genes are highleghted with bold typing.

**Table S12. Physicians and nurses at health care centers participating in the collection of FinnDiane patients.**

| FinnDiane Study Centers | Physicians and nurses |
|---|---|
| Anjalankoski Health Centre | S. Koivula, T. Uggeldahl |
| Central Finland Central Hospital, Jyväskylä | T. Forslund, A. Halonen, A. Koistinen, P. Koskiaho, M. Laukkanen, J. Saltevo, M. Tiihonen |
| Central Hospital of Åland Islands, Mariehamn | M. Forsen, H. Granlund, A-C. Jonsson, B. Nyroos |
| Central Hospital of Kanta-Häme, Hämeenlinna | P. Kinnunen, A. Orvola, T. Salonen, A. Vähänen |
| Central Hospital of Länsi-Pohja, Kemi | H. Laukkanen, P. Nyländen, A. Sademies |
| Central Ostrabothnian Hospital District, Kokkola | S. Anderson, B. Asplund, U. Byskata, P. Liedes, M. Kuusela, T. Virkkala |
| City of Espoo Health Centre | |
| Espoonlahti | A. Nikkola, E. Ritola |
| Tapiola | M. Niska, H. Saarinen |
| Samaria | E. Oukko-Ruponen, T. Virtanen |
| Viherlaakso | A. Lyytinen |
| City of Helsinki Health Centre | |
| Puistola | H. Kari, T. Simonen |
| Suutarila | A. Kaprio, J. Kärkkäinen, B. Rantaeskola |
| Töölö | P. Kääriäinen, J. Haaga, A-L. Pietiläinen |
| City of Hyvinkää Health Centre | S. Klemetti, T. Nyandoto, E. Rontu, S. Satuli-Autere |
| City of Vantaa Health Centre | |
| Korso | R. Toivonen, H. Virtanen |
| Länsimäki | R. Ahonen, M. Ivaska-Suomela, A. Jauhiainen |
| Martinlaakso | M. Laine, T. Pellonpää, R. Puranen |
| Myyrmäki | A. Airas, J. Laakso, K. Rautavaara |
| Rekola | M. Erola, E. Jatkola |
| Tikkurila | R. Lönnblad, A. Malm, J. Mäkelä, E. Rautamo |
| Heinola Health Centre | P. Hentunen, J. Lagerstam |
| Helsinki University Central Hospital, Department of Medicine, Division of Nephrology | A. Ahola, J. Fagerudd, M. Feodoroff, D. Gordin, O. Heikkilä, K Hietala, L. Kyllönen, J. Kytö, S. Lindh, K. Pettersson-Fernholm, M. Rosengård-Bärlund, M. Rönnback, A. Sandelin, A-R Salonen, L. Salovaara, L. Thorn, J. Tuomikangas, T. Vesisenaho, J. Wadén |
| Herttoniemi Hospital, Helsinki | V. Sipilä |
| Hospital of Lounais-Häme, Forssa | T. Kalliomäki, J. Koskelainen, R. Nikkanen, N. Savolainen, H. Sulonen, E. Valtonen |

| FinnDiane Study Centers | Physicians and nurses |
|---|---|
| Iisalmi Hospital | E. Toivanen |
| Jokilaakso Hospital, Jämsä | A. Parta, I. Pirttiniemi |
| Jorvi Hospital, Helsinki University Central Hospital | S. Aranko, S. Ervasti, R. Kauppinen-Mäkelin, A. Kuusisto, T. Leppälä, K. Nikkilä, L. Pekkonen |
| Jyväskylä Health Centre, Kyllö | K. Nuorva, M. Tiihonen |
| Kainuu Central Hospital, Kajaani | S. Jokelainen, P. Kemppainen, A-M. Mankinen, M. Sankari |
| Kerava Health Centre | H. Stuckey, P. Suominen |
| Kirkkonummi Health Centre | A. Lappalainen, M. Liimatainen, J. Santaholma |
| Kivelä Hospital, Helsinki | A. Aimolahti, E. Huovinen |
| Koskela Hospital, Helsinki | V. Ilkka, M. Lehtimäki |
| Kotka Heath Centre | E. Pälikkö-Kontinen, A. Vanhanen |
| Kouvola Health Centre | E. Koskinen, T. Siitonen |
| Kuopio University Hospital | E. Huttunen, R. Ikäheimo, P. Karhapää, P. Kekäläinen, M. Laakso, T. Lakka, E. Lampainen, L. Moilanen, L. Niskanen, U. Tuovinen, I. Vauhkonen, E. Voutilainen |
| Kuusamo Health Centre | T. Kääriäinen, E. Isopoussu |
| Kuusankoski Hospital | E. Kilkki, I. Koskinen, L. Riihelä |
| Laakso Hospital, Helsinki | T. Meriläinen, P. Poukka, R. Savolainen, N. Uhlenius |
| Lahti City Hospital | A. Mäkelä, M. Tanner |
| Lapland Central Hospital, Rovaniemi | L. Hyvärinen, S. Severinkangas, T. Tulokas |
| Lappeenranta Health Centre | P. Linkola, I. Pulli |
| Lohja Hospital | T. Granlund, M. Saari, T. Salonen |
| Loimaa Health Centre | A. Mäkelä, P. Eloranta |
| Länsi-Uusimaa Hospital, Tammisaari | I-M. Jousmaa, J. Rinne |
| Malmi Hospital, Helsinki | H. Lanki, S. Moilanen, M. Tilly-Kiesi |
| Mikkeli Central Hospital | A. Gynther, R. Manninen, P. Nironen, M. Salminen, T. Vänttinen |
| Mänttä Regional Hospital | I. Pirttiniemi, A-M. Hänninen |
| North Karelian Hospital, Joensuu | U-M. Henttula, P. Kekäläinen, M. Pietarinen, A. Rissanen, M. Voutilainen |
| Nurmijärvi Health Centre | A. Burgos, K. Urtamo |
| Oulankangas Hospital, Oulainen | E. Jokelainen, P-L. Jylkkä, E. Kaarlela, J. Vuolaspuro |
| Oulu Health Centre | L. Hiltunen, R. Häkkinen, S. Keinänen-Kiukaanniemi |
| Oulu University Hospital | R. Ikäheimo |
| Päijät-Häme Central Hospital | H. Haapamäki, A. Helanterä, S. Hämäläinen, V. Ilvesmäki, H. Miettinen |
| Palokka Health Centre | P. Sopanen, L. Welling |
| Pieksämäki Hospital | V. Javtsenko, M. Tamminen |

| FinnDiane Study Centers | Physicians and nurses |
|---|---|
| Pietarsaari Hospital | M-L. Holmbäck, B. Isomaa, L. Sarelin |
| Pori City Hospital | P. Ahonen, P. Merensalo, K. Sävelä |
| Porvoo Hospital | M. Kallio, B. Rask, S. Rämö |
| Raahe Hospital | A. Holma, M. Honkala, A. Tuomivaara, R. Vainionpää |
| Rauma Hospital | K. Laine, K. Saarinen, T. Salminen |
| Riihimäki Hospital | P. Aalto, E. Immonen, L. Juurinen |
| Salo Hospital | A. Alanko, J. Lapinleimu, P. Rautio, M. Virtanen |
| Satakunta Central Hospital, Pori | M. Asola, M. Juhola, P. Kunelius, M-L. Lahdenmäki, P. Pääkkönen, M. Rautavirta |
| Savonlinna Central Hospital | E. Korpi-Hyövälti, T. Latvala, E. Leijala |
| South Karelia Central Hospital, Lappeenranta | T. Ensala, E. Hussi, R. Härkönen, U. Nyholm, J. Toivanen |
| Tampere Health Centre | A. Vaden, P. Alarotu, E. Kujansuu, H. Kirkkopelto-Jokinen, M. Helin, S. Gummerus, L. Calonius, T. Niskanen, T. Kaitala, T. Vatanen |
| Tampere University Hospital | I. Ala-Houhala, T. Kuningas, P. Lampinen, M. Määttä, H. Oksala, T. Oksanen, K. Salonen, H. Tauriainen, S. Tulokas |
| Tiirismaa Health Centre, Hollola | T. Kivelä, L, Petlin, L. Savolainen |
| Turku Health Centre | I. Hämäläinen, H. Virtamo, M. Vähätalo |
| Turku University Central Hospital | K. Breitholz, R. Eskola, K. Metsärinne, U. Pietilä, P. Saarinen, R. Tuominen, S. Äyräpää |
| Vaajakoski Health Centre | K. Mäkinen, P. Sopanen |
| Valkeakoski Regional Hospital | S. Ojanen, E. Valtonen, H. Ylönen, M. Rautiainen, T. Immonen |
| Vammala Regional Hospital | I. Isomäki, R. Kroneld, M. Tapiolinna-Mäkelä |
| Vaasa Central Hospital | S. Bergkulla, U. Hautamäki, V-A. Myllyniemi, I. Rusk |

**Table S13: Members of the SUMMIT consortium.**

| Partner | Name | Position |
|---|---|---|
| 1 | **Michael Mark** | **Coordinator, WP6 leader** |
| Boehringer-Ingelheim | Markus Albertini | Project manager |
| Ingelheim, Germany | Carine Boustany | Chronic Kidney Disease, Head of Lab |
| | Alexander Ehlgen | Transmed |
| | Martin Gerl | Biomarker & Bioanlysis, Groupleader |
| | Jochen Huber | In vivo Scientist CMDR, Head of Lab |
| | Corinna Schölch | Biomarker & Bioanlysis, Head of Lab |
| | Heike Zimdahl-Gelling | Pharmacogenomics, Head of Lab |
| 2 | **Leif Groop** | Prof. Endocrinology; Coordinator Managing entity IMI-JU; PI; **WP1 and WP6 leader** |
| Lund University | Elisabet Agardh | Prof. Ophthalmology |
| Clinical Research Centre | Emma Ahlqvist | Postdoc |
| Malmö, Sweden | Tord Ajanki | Communication strategist |
| | Nibal Al Maghrabi | Research nurse |
| | Peter Almgren | Biostatistician |
| | Jan Apelqvist | Diabetologist |
| | Eva Bengtsson | Assis. Prof. Cardiovascular research |
| | Lisa Berglund | Postdoc |
| | Harry Björckbacka | Assis. Prof. Cardiovascular research |
| | Ulrika Blom-Nilsson | LUDC administrator |
| | Mattias Borell | Website, server management |
| | Agneta Burström | Research nurse |
| | Corrado Cilio | Assoc. Prof. Cellular autoimmunity |
| | Magnus Cinthio | Assist. Prof. Electrical Measurements, Lund Technical University |
| | Karl Dreja | Nephrologist |
| | Pontus Dunér | Postdoc Exp. Cardiovasc. Research |
| | Daniel Engelbertsen | PhD student Exp. Cardiovasc. Research |
| | Joao Fadista | Postdoc |
| | Maria Gomez | Assoc. Prof. Cardiovascular disease, **WP4 co-leader** |
| | Isabel Goncalves | Assis. Prof. Cardiovascular research |

| | | |
|---|---|---|
| | Bo Hedblad | Prof. Cardiovascular epidemiology |
| | Anna Hultgårdh | Prof. Vessel Wall Biology |
| | Martin E. Johansson | Pathologist |
| | Cecilia Kennbäck | Laboratory Engineer |
| | Jasmina Kravic | Database manager |
| | Claes Ladenvall | Genetic statistician |
| | Åke Lernmark | Prof. Type 1 diabetes and celiac disease |
| | Eero Lindholm | Physician, Researcher Diabetic Complications |
| | Charlotte Ling | Assist. Prof. Epigenetics |
| | Holger Luthman | Prof. Medical genetics |
| | Olle Melander | Assoc. Prof. Hypertension and cardiovascular disease |
| | Malin Neptin | Biomedical analyst |
| | Jan Nilsson | Prof. Experimental Cardiovascular research, **WP3 leader** |
| | Peter Nilsson | Prof. Internal medicine |
| | Tobias Nilsson | PhD student Electrical Measurements, Lund Technical University |
| | Gunilla Nordin Fredriksson | Prof. Cardiovascular research |
| | Marju Orho-Melander | Prof. Genetic epidemiology |
| | Emilia Ottoson-Laakso | PhD student |
| | Annie Persson | Research nurse |
| | Margaretha Persson | Laboratory Engineer |
| | Mats-Åke Persson | Database manager |
| | Jacqueline Postma | Project manager |
| | Elisabeth Pranter | Research nurse |
| | Sara Rattik | PhD student Exp. Cardiovasc. Research |
| | Gunnar Sterner | Chief physician Internal Medicine Research Unit |
| | Lilian Tindberg | Research nurse |
| | Maria Wigren | Postdoc Exp. Cardiovasc. Research |
| | Anna Zetterqvist | PhD student |
| | Mikael Åkerlund | Postdoc |
| | Gerd Östling | Laboratory Engineer |
| 3 | **Timo Kanninen** | Technical director; PI |

| | | |
|---|---|---|
| Biocomputing Platforms | Anni Ahonen-Bishopp | Software development manager |
| (BC Platforms) | Anita Eliasson | Financial and administrative director |
| Espoo, Finland | Timo Herrala | System (server) specialist |
| | Päivi Tikka-Kleemola | Service manager |
| 4 | **Anders Hamsten** | Prof. Cardiovascular disease; Atherosclerosis Research Unit; PI |
| Karolinska Institute | Christer Betsholtz | Prof. Vascular biology |
| Stockholm, Sweden | Ami Björkholm | Administrator |
| | Ulf de Faire | Professor emeritus Cardiovascular epidemiology |
| | Fariba Foroogh | Research engineer |
| | Guillem Genové | Scientist |
| | Karl Gertow | Research Assist. Prof. Cardiovascular genetics |
| | Bruna Gigante | Assoc. Professor Cardiovascular epidemiology |
| | Bing He | Postdoc |
| | Karin Leander | Assoc. Professor Cardiovascular epidemiology |
| | Olga McLeod | Postdoc |
| | Maria Nastase-Mannila | Postdoc |
| | Jaako Patrakka | Postdoc |
| | Angela Silveira | Assoc. Prof. Cardiovascular genetics |
| | Rona Strawbridge | Postdoc |
| | Karl Tryggvason | Prof. Medical Chemistry |
| | Max Vikström | Statistician |
| | John Öhrvik | Professor |
| | Anne-May Österholm | Postdoc |
| 5 | **Barbara Thorand** | Nutritional scientist, epidemiologist |
| Helmholtz Centre | Christian Gieger | Statistician |
| Munich, Germany | Harald Grallert | Biologist |
| | Tonia Ludwig | Statistician |
| | Barbara Nitz | Scientist |
| | Andrea Schneider | Data manager |
| | Rui Wang-Sattler | Scientist |

| | | |
|---|---|---|
| | Astrid Zierer | Statistician |
| 6 | **Giuseppe Remuzzi** | Institute director; PI |
| Mario Negri Institute for | Ariela Benigni | Head of department Molecular Medicine |
| Pharmacological Research | Roberta Donadelli | Scientist |
| | Maria Domenica Lesti | Researcher |
| Bergamo, Italy | Marina Noris | Head Laboratory Immunology and genetics of transplantation and rare diseases |
| | Norberto Perico | Senior scientist |
| | Annalisa Perna | Biostatistician |
| | Rossella Piras | Postdoc |
| | Piero Ruggenenti | Head of department Renal medicine, Assist. Prof. Nephrology and dialysis |
| | Erica Rurali | Postdoc |
| 7 | **David Dunger (att: Jane Horsford)** | Prof. Paediatrics; PI |
| University of Cambridge | Ludo Chassin | Senior Data Manager |
| UK | Neil Dalton, London | Clinical biochemistry |
| | John Deanfield, London | Paediatric cardiology |
| | Jane Horsford | PA to Prof. Dunger |
| | Clare Rice | Operations manager/financial contact |
| | James Rudd | Cardiovascular imaging |
| | Neil Walker | Head Data services |
| | Karen Whitehead | Technician |
| | Max Wong | Postdoc |
| 8 | **Helen Colhoun** | Prof. Public health and epidemiology; PI; Vice coordinator Managing entity; **WP2 leader** |
| | Fiona Adams | |
| University of Dundee | Tahira Akbar | PA to Helen Colhoun |
| Scotland | Jill Belch | Prof. Vasucular disease |
| | Harshal Deshmukh | PhD student |
| | Fiona Dove | |
| | Angela Ellingford | NHS Tayside Diabetic Retinopathy Screening Programme manager |
| | Bassam Farran | Statistician |
| | Mike Ferguson | Dean of research Biological chemistry and drug discovery |

| | | |
|---|---|---|
| | Gary Henderson | |
| | Graeme Houston | Consultant radiologist/senior lecturer |
| | Faisel Khan | Reader, Vascular & Inflammatory Diseases Research Unit |
| | Graham Leese | Consultant diabetologist/reader |
| | Yiyuan Liu | PhD student |
| | Shona Livingstone | Senior statistician |
| | Helen Looker | Epidemiologist |
| | Margaret McCann | Project assistant |
| | Stuart McGurnaghan | Lead data programmer |
| | Andrew Morris | Prof. Diabetic medicine |
| | David Newton | |
| | Colin Palmer | Prof. Pharmacogenomics |
| | Ewan Pearson | Consultant diabetologist/senior lecturer |
| | Gillian Reekie | Research Nurse |
| | Natalie Smith | Research Nurse |
| 9 | **Angela Shore** | Prof. Cardiovascular Science, PI |
| Peninsula Medical School | Kuni Aizawa | Postdoc |
| Exeter, UK | Claire Ball | Research nurse |
| | Nick Bellenger | Cardiologist |
| | Francesco Casanova | Associate Research Fellow Vascular medicine |
| | Tim Frayling | Prof. Genetics |
| | Phil Gates | Senior lecturer Cardiovascular science |
| | Kim Gooding | Postdoc Vascular medicine |
| | Andrew Hatttersley | Prof. Molecular medicine |
| | Roland Ling | Consultant opthalmologist |
| | David Mawson | Research technician |
| | Robin Shandas | Prof. Bioengineering (Colorado) |
| | David Strain | Stroke physician, clinical lecturer |
| | Clare Thorn | Postdoc Vascular medicine |
| 10 | **Ulf Smith** | Prof. ; PI |

| | | |
|---|---|---|
| University of Gothenburg | Ann Hammarstedt | Researcher Molecular and clinical medicine |
| Sweden | Hans Häring | Prof. University of Tübingen |
| | Oluf Pedersen | Prof. Steno Centre, Copenhagen |
| | Georgio Sesti | Prof. Universtiy of Catanzaro |
| 11 | **Per-Henrik Groop** | Prof. Diabetes genetics; PI |
| | Emma Fagerholm | MSc; PhD student, genetics |
| Folkhälsan | Carol Forsblom | Clinical coordinator |
| Helsinki, Finland | Valma Harjutsalo | PhD; FinnDiane Co-PI |
| | Maikki Parkkonen | Laboratory manager |
| | Niina Sandholm | DSc(PhD); GWAS and bioinformatics, FinnDiane Co-PI |
| | Nina Tolonen | MD PhD |
| | Iiro Toppila | BSc, MSc; bioinformatician |
| | Erkka Valo | MSc; PhD student, bioinformatician |
| 12 | **Veikko Salomaa** | Prof. Epidemiology; PI; **deputy leader WP2** |
| The National Institute for Health and Welfare | Aki Havulinna | DSc. (tech), statistician |
| Helsinki, Finland | Kati Kristiansson | PhD |
| | Pia Okamo | THL press officer |
| | Tomi Peltola | PhD |
| | Markus Perola | Professor |
| | Arto Pietilä | Statistician |
| | Samuli Ripatti | Professor, Statistics |
| | Marketta Taimi | Research assistant |
| 13 | **Seppo Ylä-Herttuala** | Prof.; PI; **WP4 leader** |
| University of Eastern Finland | Mohan Babu | PhD student |
| Kuopio, Finland | Marike Dijkstra | PhD student |
| | Erika Gurzeler | PhD student |
| | Jenni Huusko | PhD student |
| | Ivana Kholová | Postdoc |
| | Markku Laakso | Prof. |

| | | |
|---|---|---|
| | Mari Merentie | PhD student |
| | Marja Poikolainen | PA Prof Ylä-Herttuala |
| 14 | **Mark McCarthy** | Prof. Human type 2 diabetes; Oxford Centre for Diabetes, Endocrinology and Metabolism; Wellcome Trust Centre for Human Genetics; PI; **deputy leader WP1** |
| University of Oxford | Will Rayner | Database manager |
| UK | Neil Robertson | Informatics |
| | Natalie van Zuydam | Postdoc |
| 15 | **Claudio Cobelli** | Prof. ; PI; **WP5 leader** |
| University of Padova | Barbara Di Camillo | Assist. Prof. |
| Italy | Francesca Finotello | PhD student |
| | Francesco Sambo | Postdoctoral fellow |
| | Gianna Toffolo | Prof. |
| | Emanuele Trifoglio | PhD student |
| 16 | **Riccardo Bellazzi** | Prof. Bioengineering; PI; **deputy leader WP5** |
| | Nicola Barbarini | Postdoctoral fellow |
| University of Pavia | Mauro Bucalo | Software engineer |
| Italy | Christiana Larizza | Assist. Prof. |
| | Paolo Magni | Assoc. Prof. |
| | Alberto Malovini | Postdoctoral fellow |
| | Simone Marini | Postdoctoral fellow |
| | Francesca Mulas | Postdoctoral fellow |
| | Silvana Quaglini | Prof. |
| | Lucia Sacchi | Assist. Prof. |
| | Francesca Vitali | |
| 17 | **Ele Ferrannini** | Prof. Medicine; PI |
| | Beatrice Boldrini | Postdoctoral fellow |
| University of Pisa | Michaela Kozakova | Senior investigator Medical Pathophysiology |
| Italy | Andrea Mari | Senior researcher Biomedical engineering (ISIB-CNR, Padova) |
| | Carmela Morizzo | Biologist, Sonographer Cardiovascular ultrasound |

| | | |
|---|---|---|
| | Lucrecia Mota | EGIR administrative office |
| | Andrea Natali | Assoc. Prof. Medicine |
| | Carlo Palombo | Assoc. Prof. Medicine; **deputy leader WP3** |
| | Elena Venturi | Researcher |
| | Mark Walker | Prof. Molecular diabetic medicine (Univ Newcastle-upon-Tyne ) |
| 18 | **Carlo Patrono** | Prof.Pharmacology; PI |
| Catholic University of Rome | Francesca Pagliaccia | PhD student |
| Italy | Bianca Rocca | Assist. Prof. Pharmacology |
| 19 | **Pirjo Nuutila** | Prof. ; PI |
| University of Turku | Johanna Haukkala | PhD student |
| Finland | Juhani Knuuti | Prof. ; Director Turku PET Centre |
| | Anne Roivainen | Prof. |
| | Antti Saraste | Adj. Prof. |
| 20 | **Paul McKeague** | Prof. Genetic Epidemiology; PI |
| University of Edinburgh | Norma Brown | Research administrator, Public Health Services |
| Scotland | Marco Colombo | Bioinformaticist |
| 21 | **Birgit Steckel-Hamann** | Deputy coordinator; PI, Manager IMI, LRL |
| Eli Lilly | Krister Bokvist | Biostatistician |
| | Sudha Shankar | Diabetologist |
| | Melissa Thomas | Translational Science |
| 22 | **Li-ming Gan** | Prof.; Translational Science Director Cardiovascular Disease; PI, **WP3 leader** |
| AstraZeneca | Suvi Heinonen | PhD, Internal AZ postdoc, Bioscience |
| | Ann-Cathrine Jönsson-Rylander | PhD, Assoc. Prof., Team Leader Bioscience, **WP4 leader** |
| | Remi Momo | Postdoctoral fellow |
| | Volker Schnecke | Informatician Translational Science, **WP5 leader** |
| | Robert Unwin | Translational Science Director Diabetic Nephropathy |
| | Anna Walentinsson | Geneticist Translational Science |

| | | |
|---|---|---|
| | Carl Whatling | Bioscientist |
| 23 | **Everson Nogoceke** | Pre-clinical and clinical aspects of metabolic and vascular disease; PI; **WP2 leader** |
| Roche | Gonzalo Durán Pacheco | Senior Research Statistician |
| | Ivan Formentini | Biomarker & Experimental Medicine Leader |
| | Thomas Schindler | Pre-clinical and clinical and clinical biomarkers |
| 24 | **Piero Tortoli** | Professor of Electronics |
| University of Florence | Luca Bassi | Postdoctoral fellow |
| | Enrico Boni | Postdoctoral fellow |
| | Alessandro Dallai | Postdoctoral fellow |
| | Francesco Guidi | Technician |
| | Matteo Lenge | PhD student |
| | Riccardo Matera | PhD student |
| | Alessandro Ramalli | PhD student |
| | Stefano Ricci | Assist. Prof. |
| | Jacopo Viti | PhD student |
| 25 | **Bernd Jablonka** | SAD internal IMI coordinator |
| Sanofi-aventis | Dan Crowther | Biomarker researcher |
| | Johan Gassenhuber | Biostatistician |
| | Sibylle Hess | Biomarker researcher |
| | Thomas Hübschle | Pharmacologist Diabetes |
| | Hans-Paul Juretschke | Imaging |
| | Hartmut Rütten | Head Translational Medicine |
| | Thorsten Sadowski | Pharmacologist Diabetes |
| | Paulus Wohlfart | Pharmacologist Diabetes |
| 26 | **Julia Brosnan** | Biochemist, (pre)clinical research CVD, Pfizer US; **WP2 leader** |
| Pfizer | Valerie Clerin | Cardio-renal biologist, WP2 |
| | Eric Fauman | Computational biologist |
| | Craig Hyde | Statistician |
| | Anders Malarstig | Human genetics, Pfizer Europé; **WP1 leader** |

| | | |
|---|---|---|
| | Nick Pullen | Renal Disease Research Director |
| | Mera Tilley | |
| | Theresa Tuthill | Imaging specialist |
| | Ciara Vangjeli | Cardiovascular genetic epidemiologist, Pfizer Europe |
| | Daniel Ziemek | Computational biologist |

**Figure S1a-p. Regional plots of newly discovered DKD associations**

a **chr2:3745215** – rs12615970 – COLEC11 – CKD min

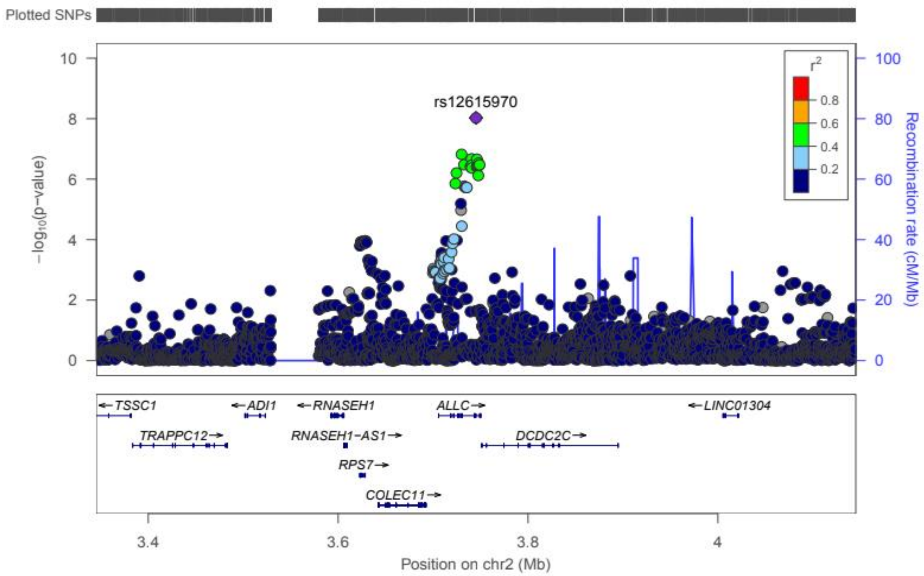

b **chr2:228121101** – rs55703767 – *COL4A3* – DN min

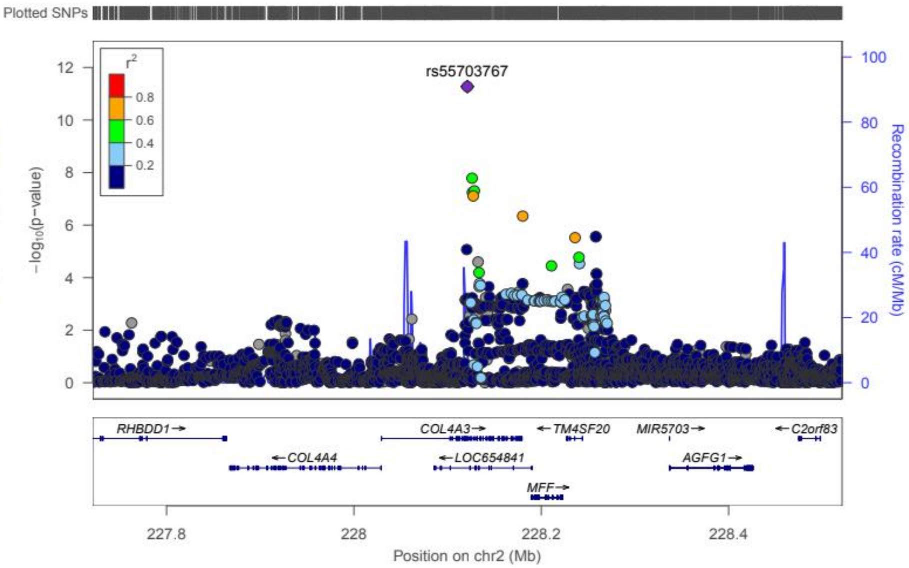

c **chr3:926345** – rs115061173 – *LINC01266* – ESRD vs ctrl min

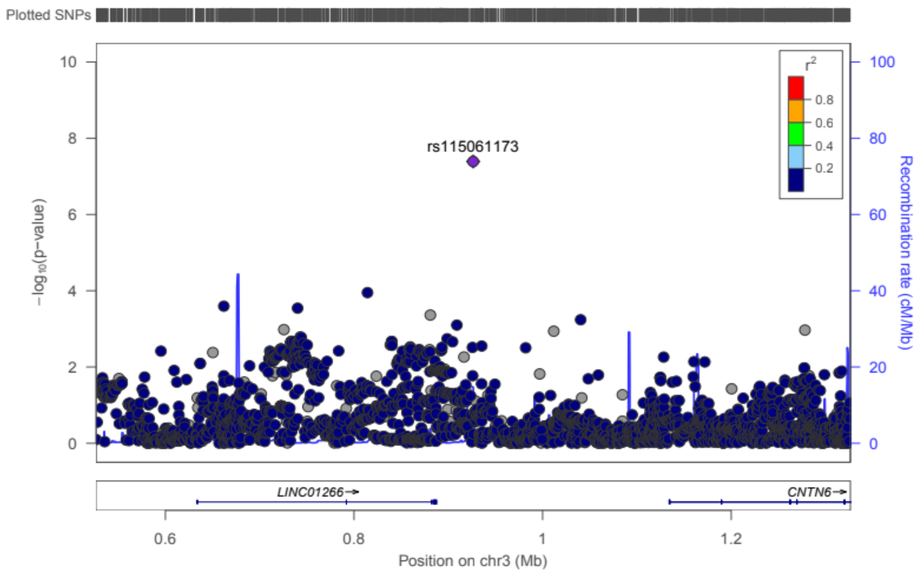

d **chr3:11910635** – rs142823282 – *TAMM41* – Micro full

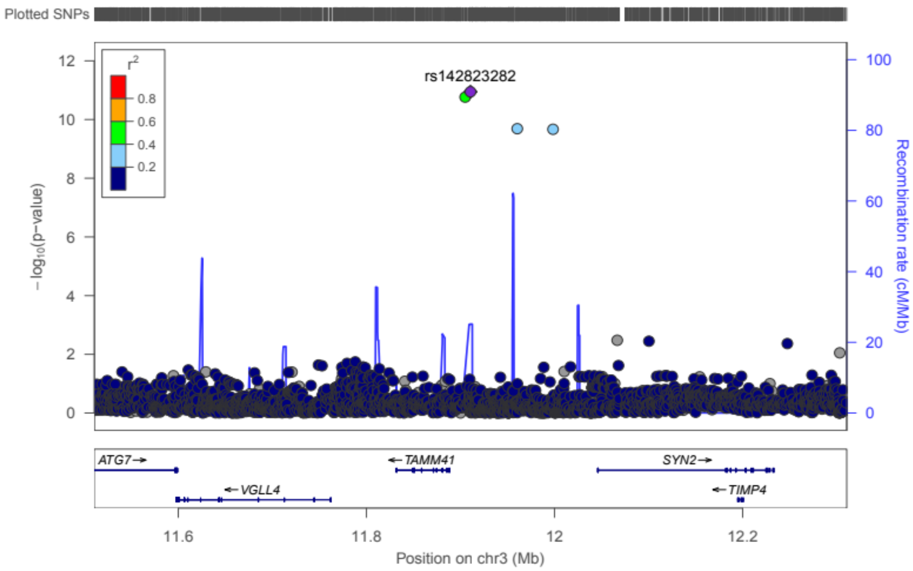

e **chr3:36566312** – rs116216059 – *STAC* – ESRD vs. non-ESRD min

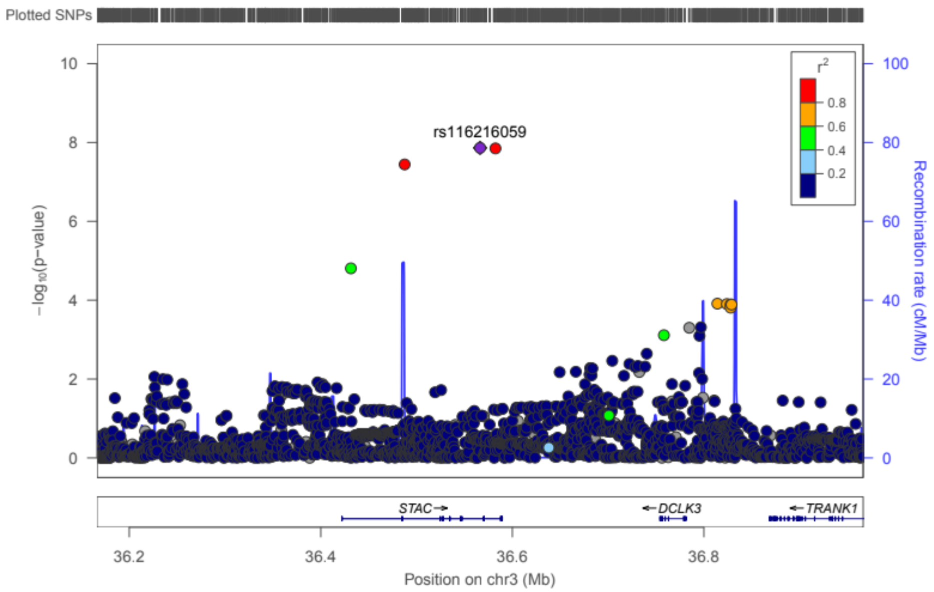

f **chr4:71358776** – rs191449639 – *MUC7* – DN min

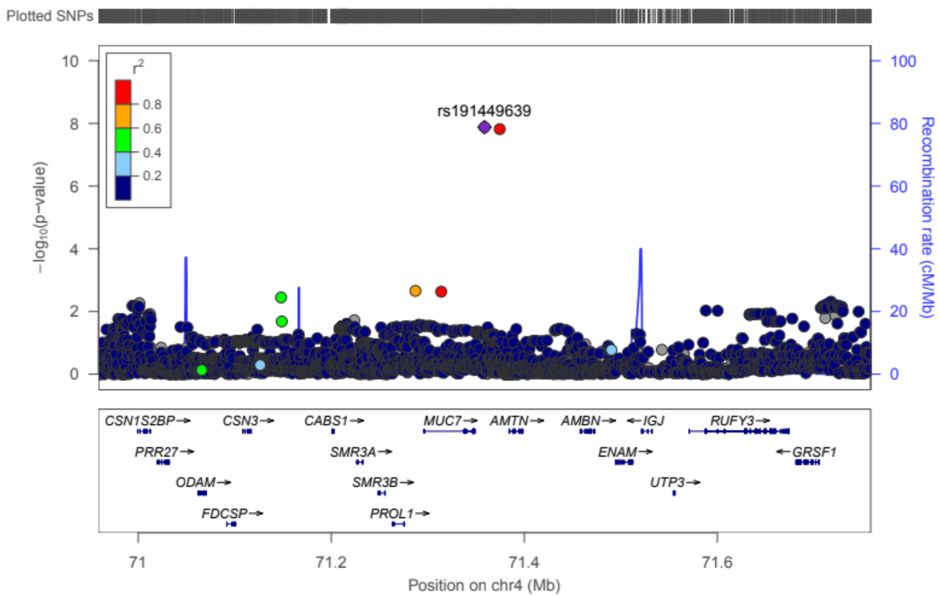

g **chr4:174500806** – rs145681168 – *HAND2-AS1* – Micro full

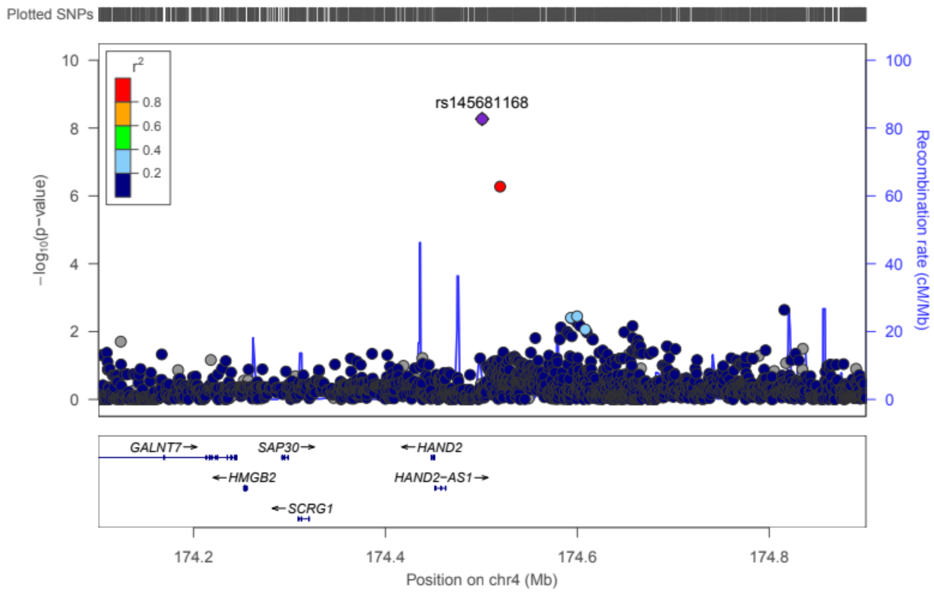

h **chr5:121774582** – rs149641852 – *SNCAIP* – CKD extreme min

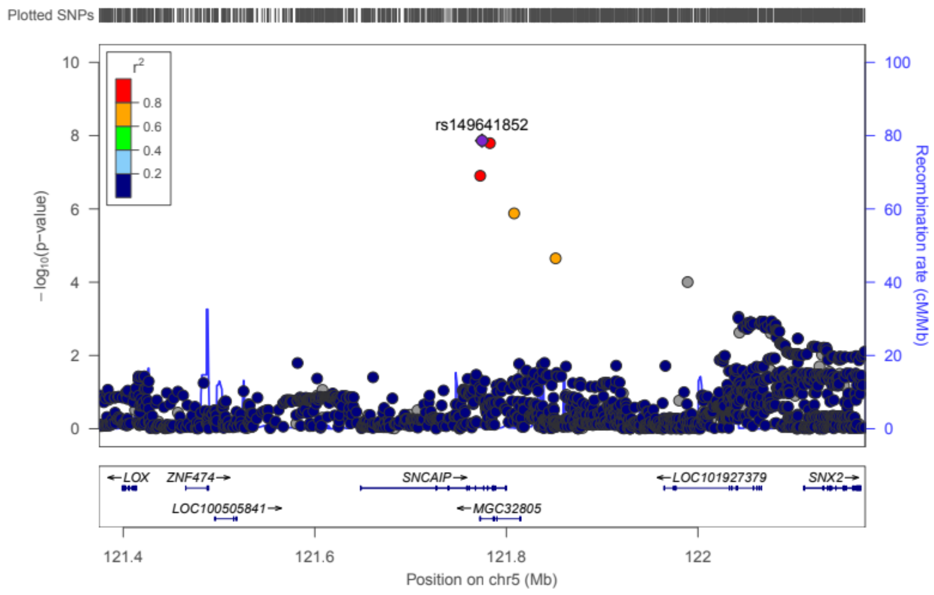

i **chr6:30865279** – rs118124843 – *DDR1* – Micro full

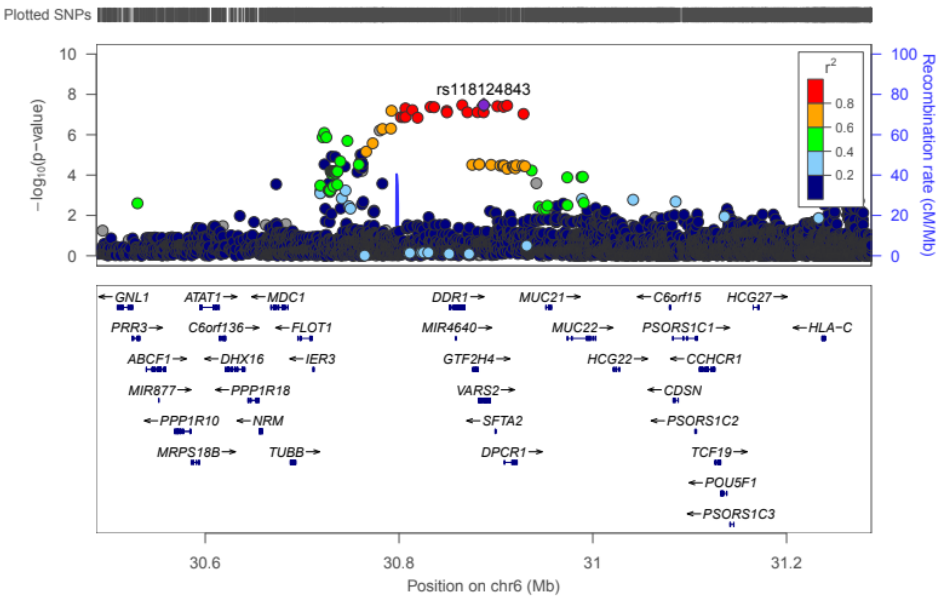

j **chr7:99728546** – rs77273076 – *MBLAC1* – Micro min

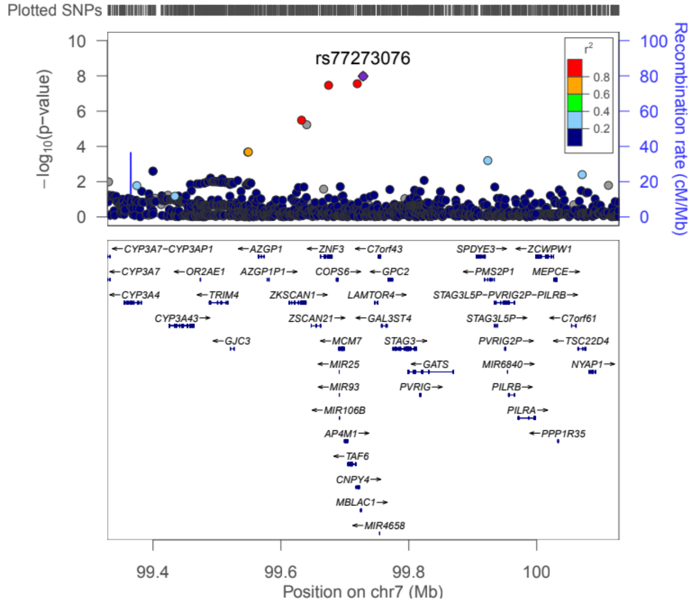

k **chr8:128100029** – rs551191707 – *PRNCR1* – ESRD vs macro min

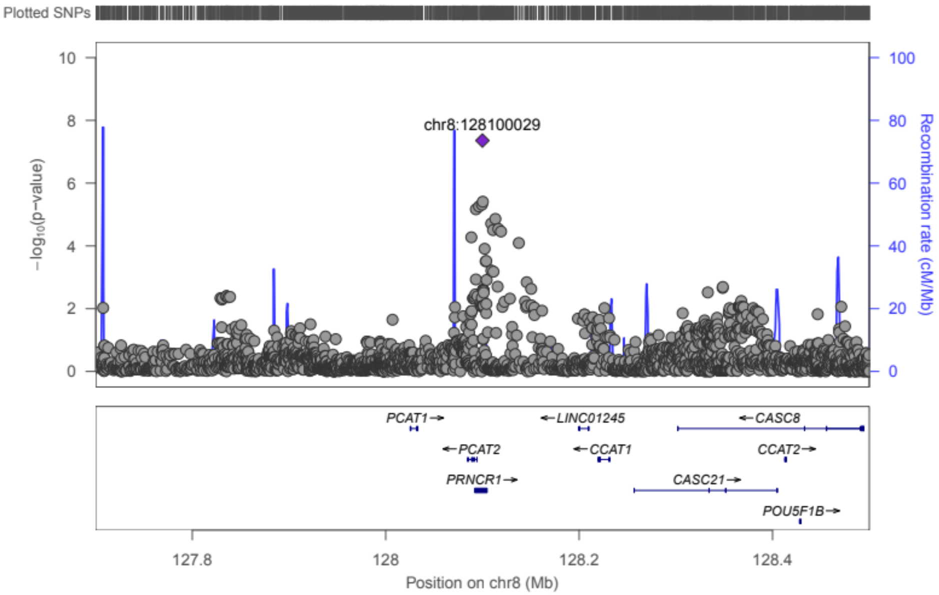

l **chr11:16937846** – rs183937294 – *PLEKHA7* – Micro min

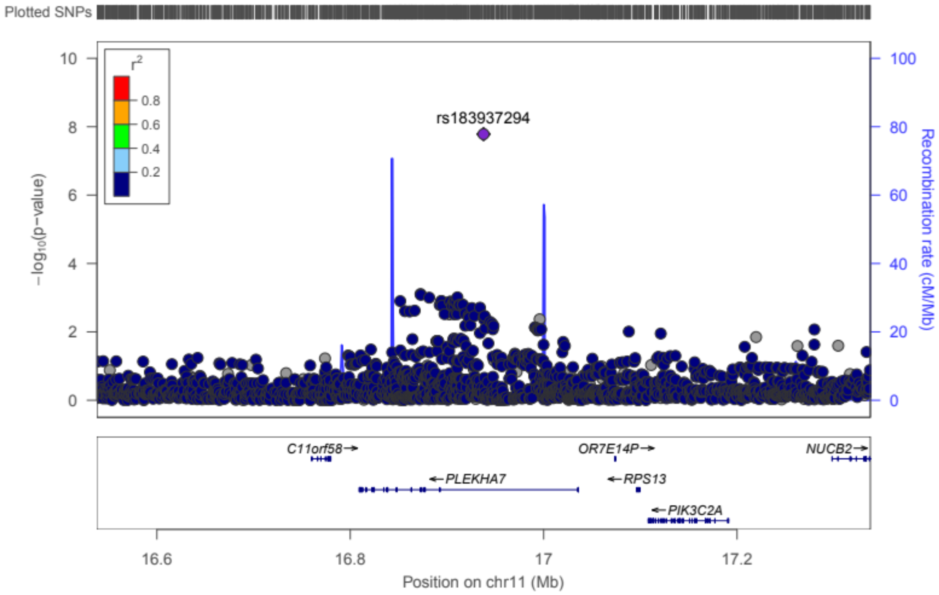

m **chr14:26004712** – rs61983410 – *STXBP6* – Micro full

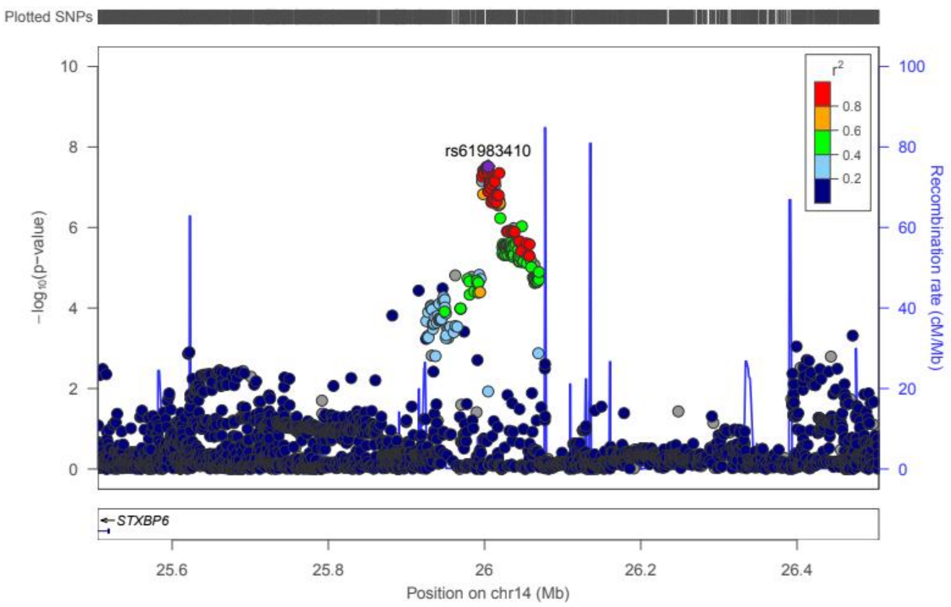

n **chr14:73740250** – rs113554206 – *PAPLN* – Macro full

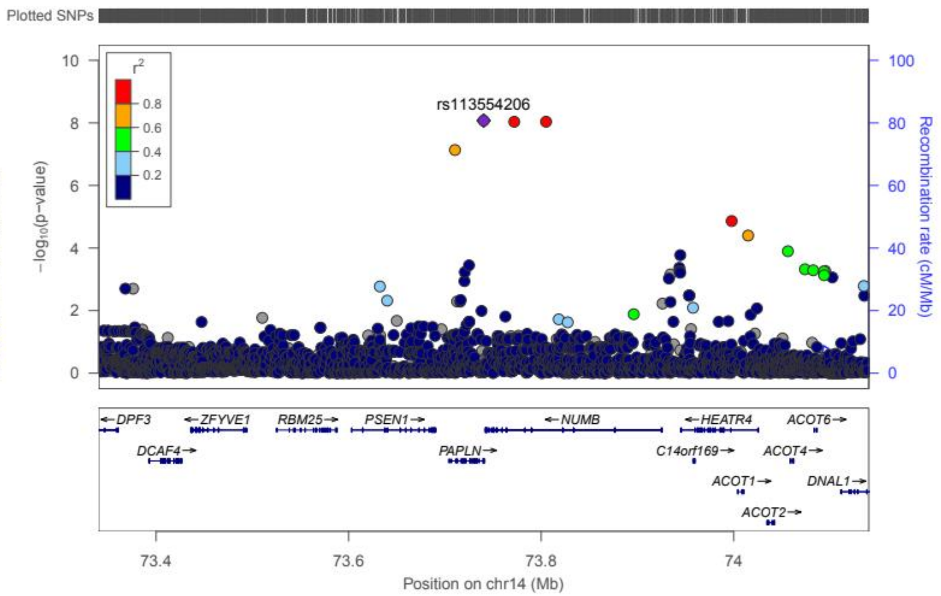

o **chr18:1811108** – rs185299109 – 18p11 – CKD min

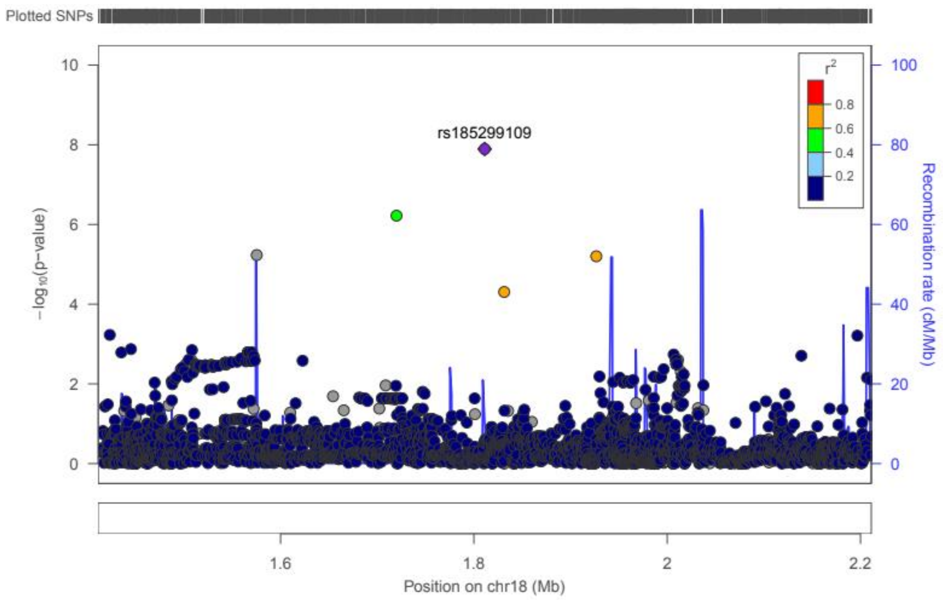

p **chr20:55837263** – rs144434404 – *BMP7* – Micro min

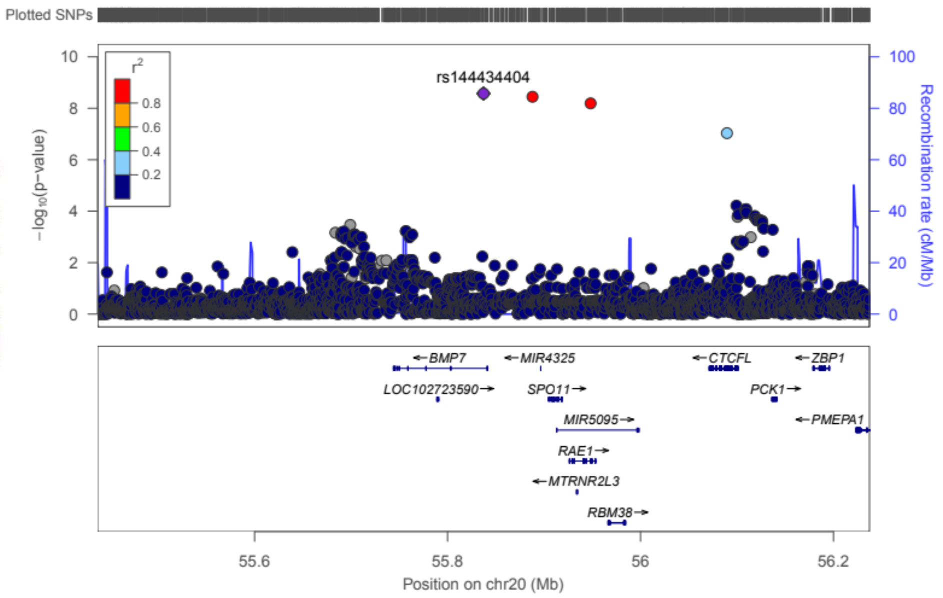

**Figure S2. Correlation of expression of *COL4A3* with degree of fibrosis and eGFR in microdissected kidney samples.**

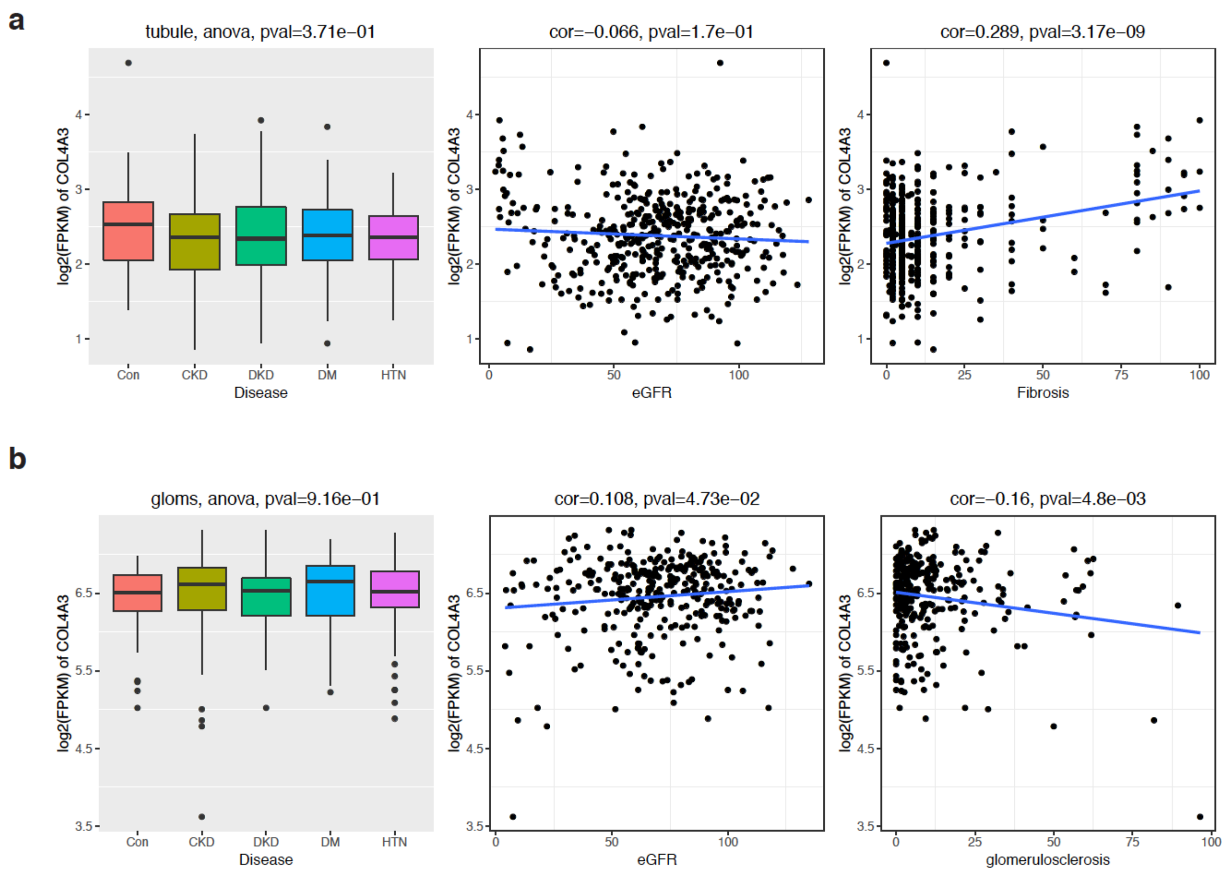

**Figure S3. Genotype – phenotype associations at the lead loci when stratified by mean $HbA_{1c}$ <7.5% in the FinnDiane study.** Only loci with a minor allele count ≥10 in each stratum are shown.

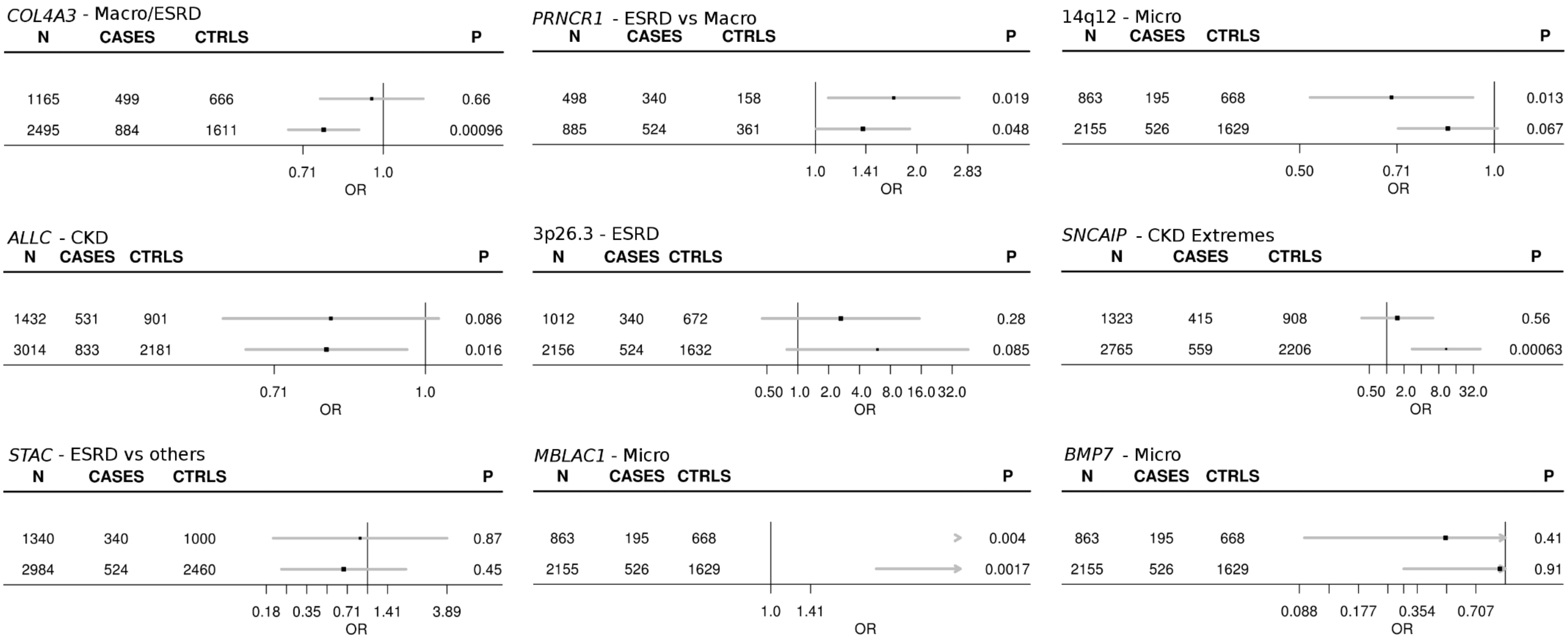

**Figure S4: Genotype – phenotype associations at the lead rs55703767 (COL4A3) locus when stratified by mean HbA1c <7.5% in up to 3226 individuals with type 2 diabetes (T2D) from the GoDARTS.**

For All vs. ctrl phenotype, 1632 individuals (848 cases, 784 controls) had HbA1c<7.5%, and 1572 individuals (874 cases, 698 controls) had HbA1c>=7.5%.

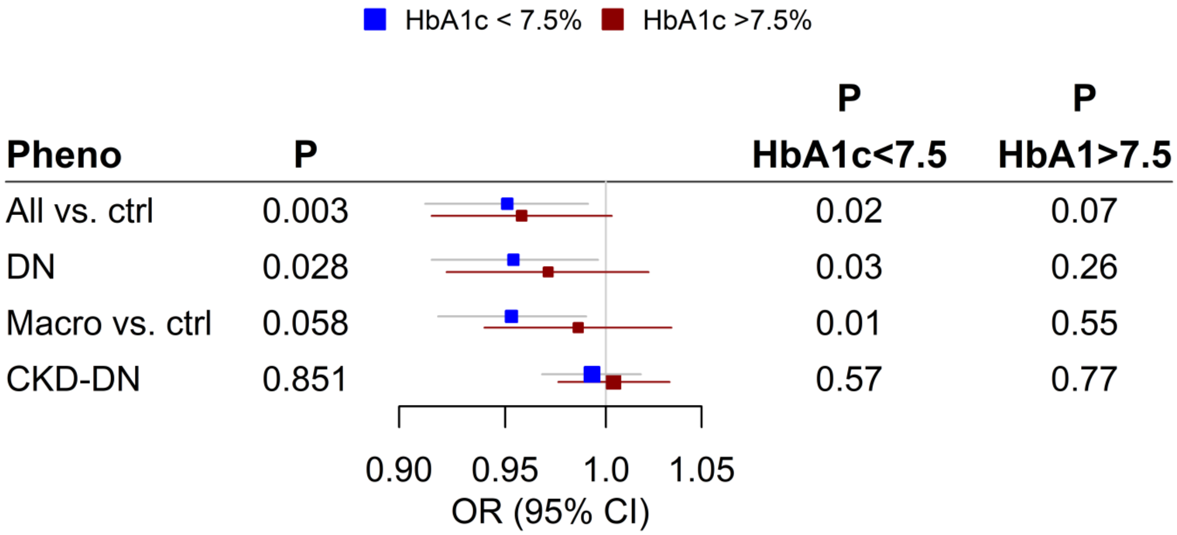

**Figure S5: Association at previously reported loci (p<5×10-8) for renal complications in individuals with diabetes.** *AFF3 and RGMA-MCTP2* were originally reported for ESRD (T1D) (Sandholm et al., 2012); *CDCA7/SP3* for ESRD in women (T1D) (Sandholm et al., 2013); *ERBB4* for DN (T1D) (Sandholm et al., 2012); *GABRR1* for microalbuminuria (T2D)(Van Zuydam et al., 2018); *GLRA3* for albuminuria (T1D) (Sandholm et al., 2014); *PRKAG2* and *UMOD* for eGFR (Pattaro et al., 2016; Van Zuydam et al., 2018); and *SCAF8/CNKSR3* for DN (T2D) (Iyengar et al., 2015).

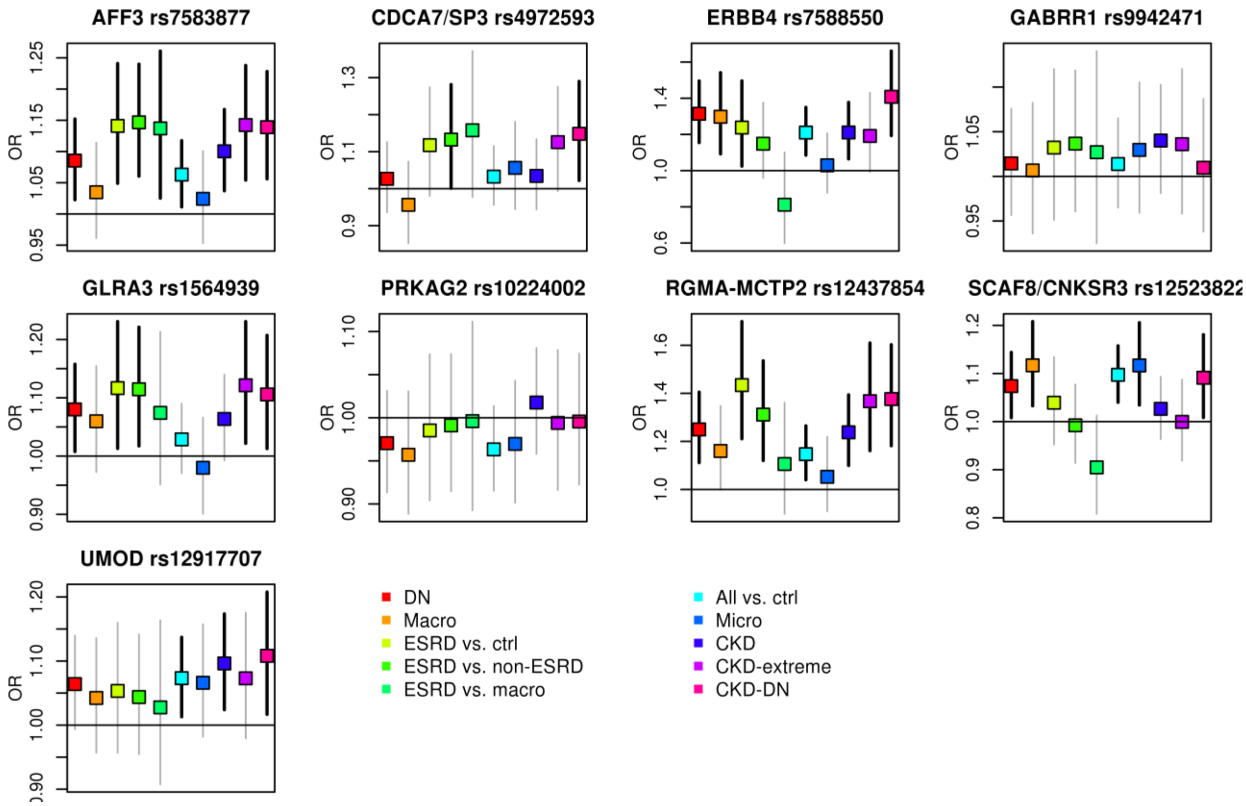

**Figure S6: Forest plots of the associations at the previously reported lead loci from the GENIE consortium with largely overlapping studies. A:** *RGMA-MCTP2* rs12437854. **B:** *AFF3* rs7583877. **C:** *ERBB4* rs7588550. Meta-analysis results for *RGMA-MCTP2*: Previous $P = 2.0\times10^{-9}$, OR = 1.80 (95% confidence interval 1.48, 2.17), Current $P = 7.4\times10^{-4}$, OR = 1.31 (1.12, 1.54); Meta-analysis results for *AFF3*: Previous $p=1.20\times10^{-8}$, OR = 1.29 (1.18, 1.40), Current p=5.97x10-4, OR=1.15 (1.06, 1.24). Meta-analysis results for *ERBB4*: Previous $P = 2.1\times10^{-7}$, OR = 0.66 (0.56, 0.77), Current $P = 3.5\times10^{-5}$, OR = 0.76 (0.67, 0.87).

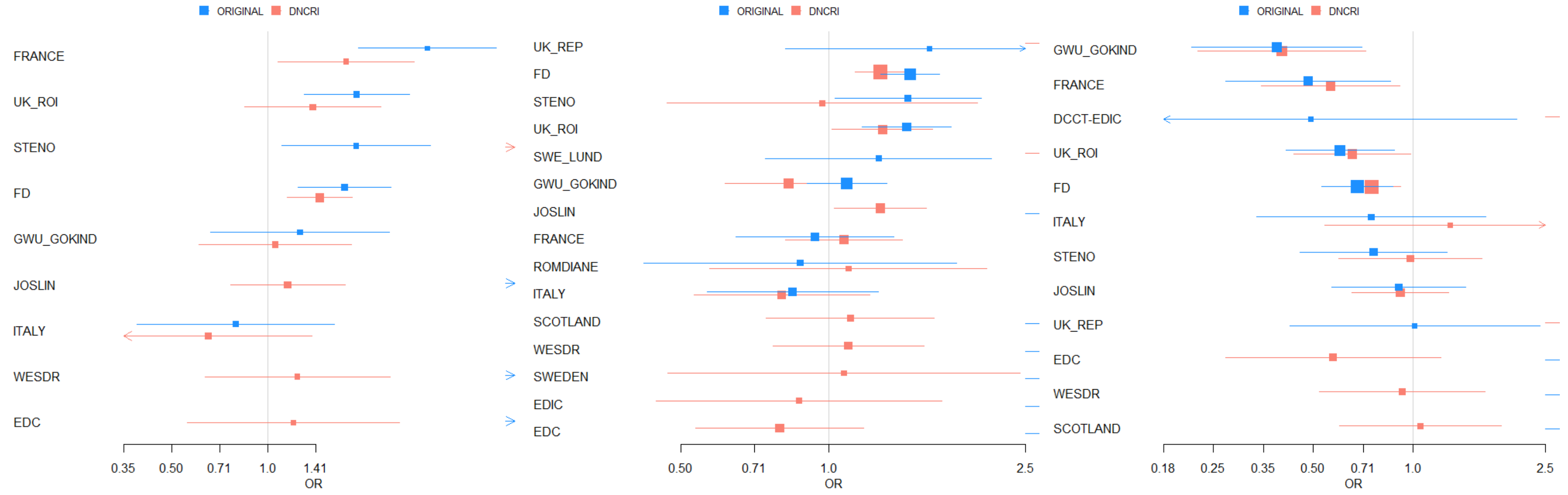

**Figure S7: Meta-analysis results for the loci that have previously been associated with DKD, or with eGFR or AER in the general population.** Figure shows OR [95% CI] for the 25 loci with $p<0.05$ for at least one sub-phenotype; associations with $p<0.05$ are indicated with black confidence intervals. Results are plotted so that odds ratio (OR)>1 indicates association in the same direction with the original report (for eGFR, this means that the allele associated with higher risk of DN is associated with lower eGFR). A total of 69 loci were evaluated, including loci for DKD (5 loci: *AFF3*, *RGMA-MCTP2*, *ERBB4* (Sandholm 2012), *CDCA7/SP3* (Sandholm2014), *SCAF8/CNKSR3* (Iyengar 2015)), for albuminuria in individuals with diabetes (*GLRA3* (Sandholm 2013), 3 suggestive loci *CUBN, HST6ST1* and *RAB38* (Teumer 2016)), for eGFR in individuals with diabetes (*UMOD*, Pattaro et al. 2016 and Van Zuydam et al. 2018, *PRKAG2* Van Zuydam et al. 2018) or without diabetes (61 loci, Gorski 2017). Associations at *AFF3, RGMA-MCTP2, ERBB4, SCAF8/CNKSR3*, and *UMOD* remained significant after correction for 69 tested loci ($p<7\times10^{-4}$).

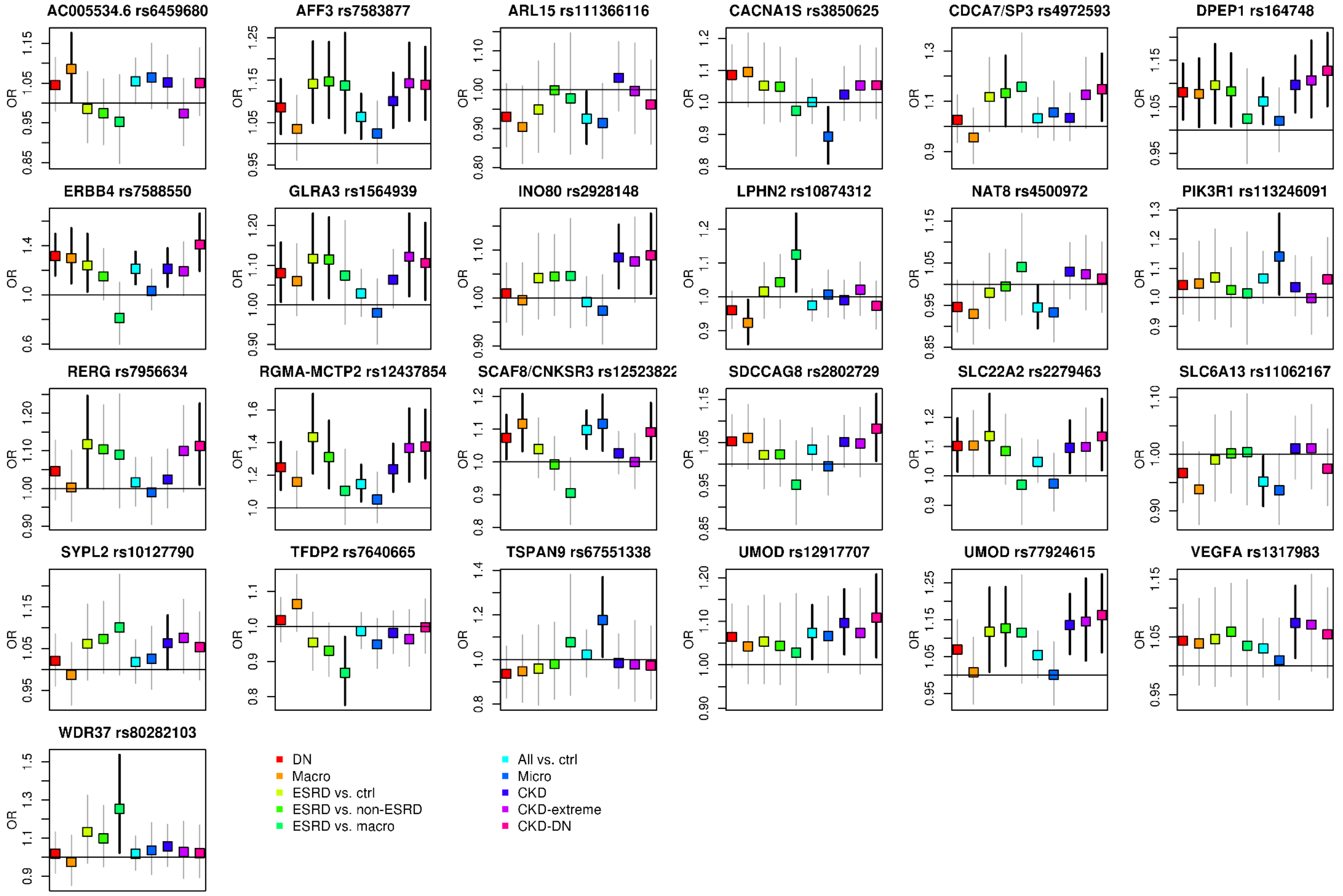
AC005534.6 rs6459680
AFF3 rs7583877
ARL15 rs111366116
CACNA1S rs3850625
CDCA7/SP3 rs4972593
DPEP1 rs164748
ERBB4 rs7588550
GLRA3 rs1564939
INO80 rs2928148
LPHN2 rs10874312
NAT8 rs4500972
PIK3R1 rs113246091
RERG rs7956634
RGMA-MCTP2 rs12437854
SCAF8/CNKSR3 rs1252382
SDCCAG8 rs2802729
SLC22A2 rs2279463
SLC6A13 rs11062167
SYPL2 rs10127790
TFDP2 rs7640665
TSPAN9 rs67551338
UMOD rs12917707
UMOD rs77924615
VEGFA rs1317983
WDR37 rs80282103
OR
DN
Macro
ESRD vs. ctrl
ESRD vs. non-ESRD
ESRD vs. macro
All vs. ctrl
Micro
CKD
CKD-extreme
CKD-DN

**Figure S8. Expression of quantitative trait loci (eQTL) analysis in microdissected tubule samples.** Boxplots showing normalized gene expression by stratified by homozygous common (red), heterozygous (green), and homozygous rare (blue) genotype. We identified nominal associations for rs55703767 in tubule samples with *IRS1* (a) and in glomerular samples with *RP11-395N3.2* and *AGFG1* (b). We also found nominal associations of rs61983410 with the gene encoding Cathepsin G, *CTSG*, in both eQTL analysis of whole kidney samples (c) and microdissected tubule samples (d).

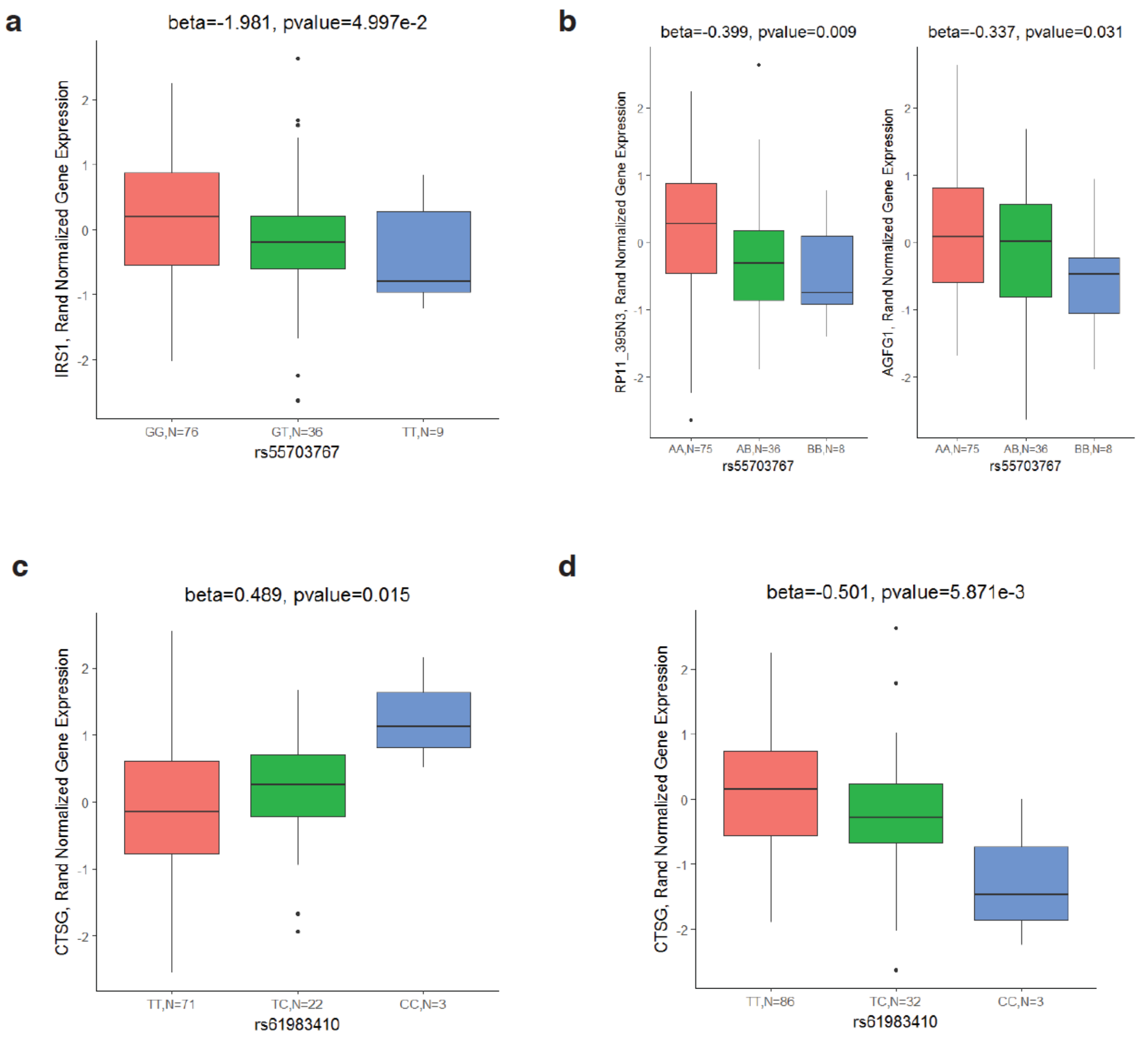

**Figure S9. Functional annotation of *TAMM41*.** ChIP-seq data derived from healthy adult human kidney samples (Bernstein et al., 2010) shows enrichment for H3K27ac, H3K9ac, H3K4me1, and H3K4me3, suggesting that this locus an active regulator of *TAMM41*.

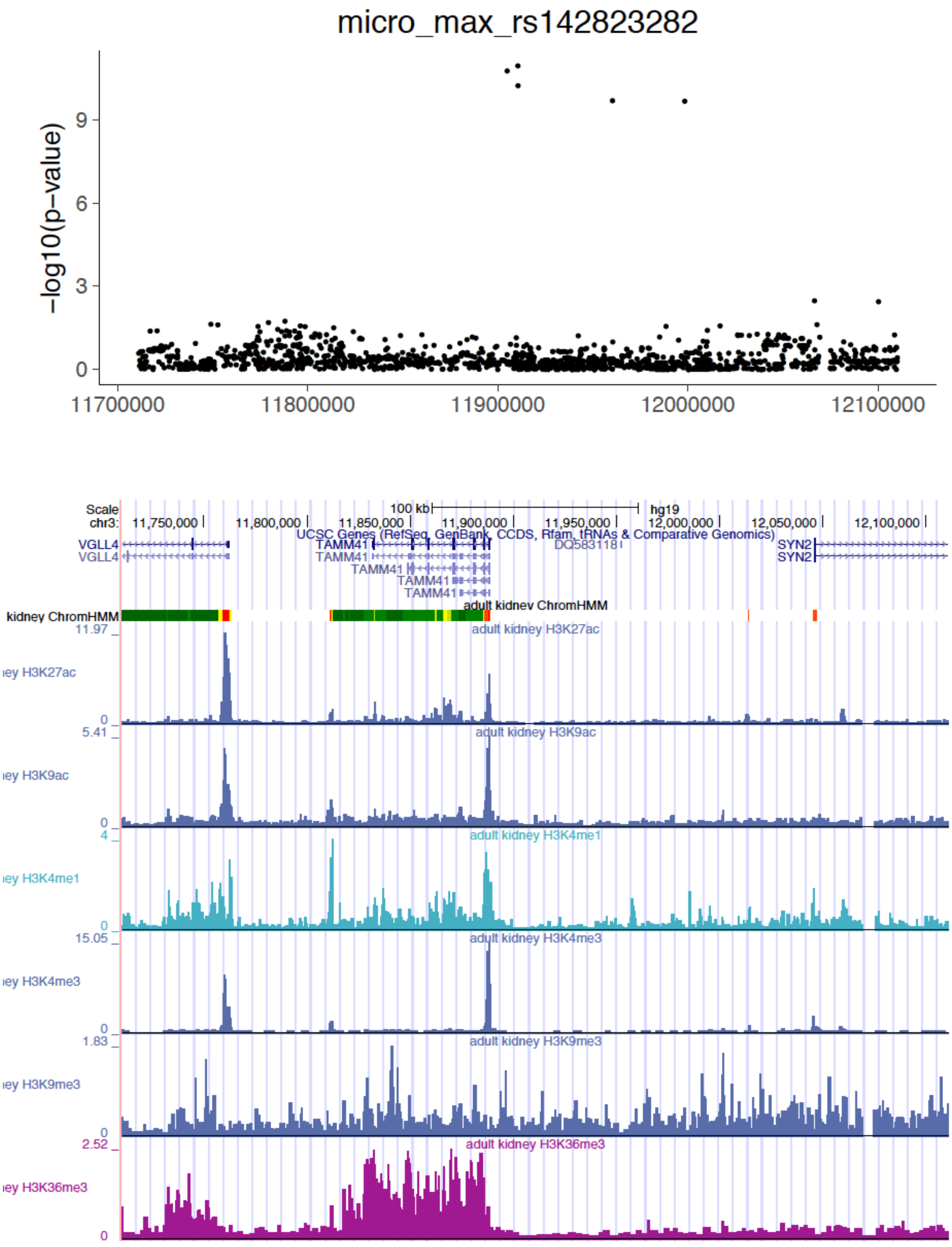

**Supplemental Methods: Cohort descriptions**

**CACTI:** The Coronary Artery Calcification in Type 1 Diabetes (CACTI) study enrolled 656 subjects with diabetes diagnosed before age 30 years, treated with insulin within 1 year of diagnosis, and diabetes duration of at least 10 years on enrollment (Dabelea et al., 2003).

**DCCT/EDIC:** The Diabetes Control and Complications Trial (DCCT) was a multi-center randomized clinical trial to compare intensive and conventional insulin therapy on the development and progression of early vascular and neurological complications of type 1 diabetes (T1D). Renal outcomes were defined as time in years from DCCT baseline until the event. AERs were measured annually in DCCT and every other year in the post-study Epidemiology of Diabetes Interventions and Complications (EDIC) cohort. Persistent microalbuminuria was defined as the time to two consecutive AER >30 mg/24 hours (>20.8 µg/min); severe nephropathy was the time to AER >300 mg/24 hours (>208 µg/min) with prior persistent microalbuminuria, or ESRD. 22% developed persistent microalbuminuria during follow-up (268 events, 976 censored), while 10% developed severe nephropathy (132 events, 1,172 censored) (Nathan, 2014; The Diabetes Control and Complications Trial Research Group;, 1995).

**EDC:** The Pittsburgh Epidemiology of Diabetes Complications (EDC) is a historical cohort study based on incident cases of childhood onset (prior to age 17 years) T1D, diagnosed or seen within one year of diagnosis (1950-80) at Children's Hospital of Pittsburgh (Orchard et al., 1990). The cohort, which has been shown to be epidemiologically representative of the Allegheny County, Pennsylvania, T1D population (Wagener et al., 1982), was first assessed for the EDC study between 1986 and 1988 (mean participant age and diabetes duration were 28 and 19 years, respectively). Subsequently, biennial examinations were conducted for 10 years, with a further detailed examination at 18 and 25 years from enrollment. All EDC study participants provided informed consent, and all study procedures were approved by the University of Pittsburgh Institutional Review Board (IRB). Microalbuminuria was defined as albumin excretion rate (AER) 20-200 µg/min (30-300 mg/24 hours), overt nephropathy as AER >200 µg/min (>300 mg/24 hours) and albuminuria as >20 µg/min (>30 mg/24 hours) in at least two of three validated timed urine collections. End-stage renal disease was defined as receiving dialysis or renal transplantation.

**FinnDiane: Finnish Diabetic Nephropathy Study (FinnDiane)** is an ongoing nationwide Finnish multicenter study of adult participants with T1D described previously [Thorn et al. 2005, Sandholm et al. 2012]. The participants were invited to the study by their attending physician who filled a questionnaire on the medical status of the patient and performed a clinical examination. A subset of the patients participated at one or more follow-up visits with a similar setting. Additional health related information was obtained from Finnish hospital discharge registry and from the patients' medical records. Further patients were included to the FinnDiane study through collaboration with the Finnish National Institute for Health and Welfare; for these participants, health related data was obtained from the hospital discharge registry and from the medical records. For this study, participants were limited to those with T1D diagnosed prior to age 40 years and with insulin treatment begun within 2 calendar years from diabetes onset. Disease status was defined by urine albumin excretion rate (AER) or urine albumin to creatinine ratio (ACR) in at least two out of three consecutive urine collections at local centers: microalbuminuria was defined as AER 20-200 µg/min or 30-300 mg/24h or an ACR of 2.5-25 mg/mmol for men and 3.5-35 mg/mmol for women in overnight, 24-hour or spot urine collections, respectively. Similarly, the limit for macroalbuminuria was AER >200 µg/min or >300 mg/24h or ACR > 25 mg/mmol for men and >35 mg/mmol for women. ESRD was defined as ongoing dialysis treatment or receipt of transplanted kidney. Control patients with normal AER were required to have T1D duration of at least 15 years (Sandholm et al., 2012; Thorn et al., 2005).

**France-Belgium:** The GENEDIAB ('Génétique de la Néphropathie Diabétique, Genetics of Diabetic Nephropathy) and Genesis subjects were recruited in France, and in France-Belgium, respectively. Patients with T1D were selected on the following criteria: 1) age at diabetes onset before age 35 years, and 2) definitive insulin use within one year after diagnosis. Diabetic nephropathy was classified according to the highest three AER measurements within the last 5 years. Categories included: 1) controls (normoalbuminuria), 2) incipient nephropathy (microalbuminuria), 3) established nephropathy (proteinuria), and 4) advanced nephropathy (serum creatinine >150 mol/L and/or renal replacement therapy) (Hadjadj et al., 2004; Marre et al., 1997).

**GoKinD US: Genetics of Kidneys in Diabetes US Study (GoKinD):** The GoKinD study consists of a DKD case-control cohort of individuals diagnosed with T1D prior to 31 years of age who began insulin treatment within 1 year of T1D diagnosis. Controls were 18-59 years of age, with T1D for at least 15 years but without DKD. DKD definition includes individuals with end-state renal disease (ESRD), dialysis or kidney transplant and persistent macroalbuminuria (at least 2 out of 3 tests positive for albuminuria by dipstick ≥1+, or ACR >300 µg albumin/mg of urine creatinine). Cases were defined as people 18-54 years of age, with T1D for at least 10 years and DKD. Individuals were recruited at two study centers, George Washington University and the Joslin Diabetes Center using differing methods (Pezzolesi et al., 2009). The Joslin GoKinD subjects were analyzed jointly with subjects from the Joslin Microalbuminuria and 50-years medalists (see below).

**The InterDiane Consortium**: The International Diabetic Nephropathy Consortium (InterDiane) was initiated in 2010 based on the protocol of the FinnDiane Study. The aim of the study is to identify risk factors for diabetic nephropathy and other chronic complications in patients with T1D. The participating studies follow the main protocol of the FinnDiane Study and use the same standardized questionnaires for data acquisition. T1D was defined as diabetes onset <40 years with insulin treatment initiated within one year of diagnosis. The main renal phenotype information has been collected at a baseline visit but in some countries prospective patient visits have been performed and additional phenotype information has been gathered. The last available phenotype information has been used in the analyses. Patients included fulfil the harmonized case and control criteria of the present study. InterDiane centers included in this study come from Romania, Austria, Latvia and Lithuania.

- **AusDiane: The Austrian Diabetic Nephropathy Study (AusDiane)** was initiated in 2012 in the state of Salzburg in Austria, and is part of the InterDiane Consortium (please see also the InterDiane cohort description). The patients have been studied during a regular visit at two hospitals (Department of Internal Medicine 1,

Paracelsus Medical University Hospital Salzburg and Diakonissen-Wehrle Hospital Salzburg). Recruitment was done consecutively in the outpatient departments of these two hospitals. Clinical data were collected mainly as part of the Type 1 diabetes Registry of the state of Salzburg. Patients have been studied repeatedly every 1 to 1.5 years to improve the phenotype. The last available phenotype is used for the analysis. This study comprises 71 patients with normal AER and diabetes duration ≥15 years, 13 with microalbuminuria, 4 with macroalbuminuria and 2 with ESRD and with GWAS data available and passing the inclusion criteria. Renal status was assessed by morning urine samples at least once every year. The study received ethical approval from the local ethics committee (Ethikkommission Salzburg). Written consent was obtained prior to participation in the study.

- **The Latvian Diabetic Nephropathy Study (LatDiane)** was initiated in 2012 and is part of the InterDiane Consortium (please see also the InterDiane cohort description). Recruitment of patients took place in Pauls Stradins University Hospital (Riga). The patients were recruited from the Endocrinology department of Pauls Stradins University Hospital and from out-patient clinics of Riga and Riga district (cities Jelgava, Jurmala, Ogre, Salaspils ect). The study comprises 80 patients with normal AER and diabetes duration ≥15 years, 33 with microalbuminuria, 18 with macroalbuminuria and 7 with ESRD and with GWAS data available and passing the inclusion criteria. Patients from out-patients clinics of Riga and Riga district were invited for a separate recruitment visit following the invitation of their endocrinologist. Patients undergoing treatment or correction of therapy in Endocrinology department of Pauls Stradins University Hospital were recruited in the department. Renal status was assessed based on available data of albuminuria (albumin content in 24-hour urine or albumin/creatinine in morning spot urine). In addition, during the recruitment visit, morning spot urine was collected from all patients, and sent for albumin/creatinine measurement. For patients without available data on measurements of albuminuria before recruitment to the LatDiane Study, albumin/creatinine determination in morning spot urine was repeated also several weeks after recruitment. Follow-up visits are planned for 2018. The study received ethical approval from the Latvian Central Ethics Committee. Written consent was obtained prior to participation in the study (Sviklāne et al., 2018).

- **The Lithuanian Diabetic Nephropathy Study (LitDiane)** was initiated in 2013 and is part of the InterDiane Consortium (please see also the InterDiane cohort description). Patients with T1D have been collected in a single center at the Hospital of Lithuanian University of Health Sciences (HLUHS) in Kaunas. Patients were included in the study from out-patient and inpatient departments of Endocrinology clinic of HLUHS during separate study visit. Medical records were reviewed for each patient and prospective visits are performed once a year. Renal status was classified based on the urinary albumin excretion rate (AER) in at least two out of three consecutive urine collections as: normal AER (<30mg/24h in a 24-hour urine collection), incipient diabetic nephropathy (microalbuminuria; AER ≥30 and <300mg/24h) or overt diabetic nephropathy (macroalbuminuria; AER ≥300mg/24h). Patients on dialysis or with a kidney transplant were considered to have end-stage renal disease (ESRD). As a measure of renal function estimated GFR (eGFR) was calculated with the Chronic Kidney Disease Epidemiology Collaboration (CKD-EPI) formula. At the time of analysis, the study comprised 39 patients with normal AER, 32 with microalbuminuria, 9 with macroalbuminuria and 10 with ESRD and with GWAS data available and passing the inclusion criteria. The study received ethical approval from the Kaunas Regional Biomedical Research Ethics Committee (No. BE-2-16, 13-March-2013). Written consent was obtained prior to participation in the study. [references pending; submitted]
- **The Romanian Diabetic Nephropathy Study (RomDiane)** was initiated in 2010 in Romania as the pilot study of the InterDiane Consortium. Patients have been studied in a cross-sectional manner in two centers in Bucharest and one in Craiova between 2010 and 2012. Renal status was assessed based on the AER or ACR in two out of three consecutive urine collections at local centers. This study comprises 89 patients with normal AER and diabetes duration ≥15 years, 48 with microalbuminuria, 70 with macroalbuminuria and 28 with ESRD, and with GWAS data available and passing the inclusion criteria. The study received ethical

approval from the local ethics committee. Written consent was obtained prior to participation in the study (Pop et al., 2016).

**Italy:** Subjects with T1D were recruited at the Complications of Diabetes Unit of the San Raffaele Scientific Institute, Milan, Italy. Diabetic nephropathy was defined as a median AER >200 µg min-1 in three overnight collections of sterile urine in patients with T1D for at least 10 years, concomitant diabetic retinopathy and absence of clinical or laboratory evidence of cardiac failure or other renal or urinary tract disease. Patients without nephropathy had a median AER <20 µg/min (Sandholm et al., 2012).

**Joslin Cohort:** There were 2,271 Joslin patients with T1D included in this study. These patients were derived from three cohorts included in the ongoing Joslin Kidney Study (Krolewski, 2015). Recruitment of 1,600 patients into the 1$^{st}$ Joslin Kidney Study on Natural History of Microalbuminuria in T1D took place between 1991 and 1993, and the cohort was followed through 2004. Recruitment of 1,108 patients into the 2$^{nd}$ Joslin Kidney Study on Natural History of Early Renal Decline in T1D took place between 2003 and 2012 and the follow-up of this cohort is still ongoing. The Joslin Proteinuria Cohort that included 630 patients was assembled from among those who developed proteinuria while attending the Joslin Clinic between 1991 and 2004. The follow-up of this cohort is still ongoing. In the analysis of data for this study, the kidney phenotypes of patients at the enrollment into the Joslin Kidney Study were considered. Genotyping data were available for 244 patients with ESRD, 475 patients with proteinuria, 470 patients with microalbuminuria and 1,189 patients with normoalbuminuria.

**SDRNT1BIO:** The Scottish Diabetes Research Network Type 1 Bioresource is a prospective cohort study of 6,127 individuals from across Scotland. Participants aged 16 years and over with a clinical diagnosis of T1D and insulin use within a year of onset were recruited from primary and secondary care across Scotland between 2010 and 2013. Serum, plasma, whole blood and urine samples were collected at study day allowing eGFR and albuminuria status to be obtained. Further retrospective and prospective biochemistry, co-morbidity and lifestyle data were linked from routine electronic health care records, providing serial estimates of renal status (Akbar et al., 2016).

**Steno:** Patients with T1D attending the outpatient clinic at Steno Diabetes Center were invited to participate in a study of genetic risk factors for diabetes complications. T1D was considered present if the age at onset of diabetes was ≤35 years and time to definite insulin therapy ≤1 year. DKD was defined by persistent albuminuria (>300 mg/24 h) in two out of three consecutive measurements, presence of retinopathy, and absence of other kidney or urinary tract disease. Absence of DKD (controls) was defined as persistent normoalbuminuria (<30 mg/24 h) after more than 15 years of T1D in patients not treated with ACE inhibitors or angiotensin-II receptor blockers. ESRD was defined as chronic dialysis or kidney transplantation (Rossing et al., 2002).

**Sweden:** All patients with T1D were Swedish and diagnosed before 30 years of age. The patients with macroalbuminuria (urinary AER ≥ 200 µg min-1 in at least two consecutive overnight samples) were defined as case. The patients with AER <20 µg min-1 were considered as control (Ma et al., 2006).

**UK-ROI:** In the United Kingdom (UK) GoKinD, Warren 3 and All Ireland (UK-ROI) study, data were collected under a parallel protocol to that of the GoKinD study in the United States (see above). Briefly, all individuals are white with parents and grandparents born in the UK or Ireland and who had T1D diagnosed before 31 years of age. Cases have DKD diagnosed by the onset of proteinuria (>0.5 g/24 hr) >10 years since diagnosis of diabetes; controls are diabetic individuals without evidence of proteinuria (or microalbuminuria) >15 years after onset of diabetes (McKnight et al., 2010).

**WESDR:** The Wisconsin Epidemiologic Study of Diabetic Retinopathy was an epidemiologic study of subjects with diabetes diagnosed before 30 years of age and taking insulin. Outcomes collected included proteinuria on a urine dipstick test (Klein et al., 1998).

### References

Akbar, T., McGurnaghan, S., Palmer, C.N.A., Livingstone, S.J., Petrie, J., Chalmers, J., Lindsay, R.S., McKnight, J.A., Pearson, D.W.M., Patrick, A.W., et al. (2016). Cohort Profile: Scottish Diabetes Research Network Type 1 Bioresource Study (SDRNT1BIO). Int. J. Epidemiol. *46*, dyw152.

Bernstein, B.E., Stamatoyannopoulos, J.A., Costello, J.F., Ren, B., Milosavljevic, A., Meissner, A., Kellis, M., Marra, M.A., Beaudet, A.L., Ecker, J.R., et al. (2010). The NIH Roadmap Epigenomics Mapping Consortium. Nat. Biotechnol. *28*, 1045–1048.

Dabelea, D., Kinney, G., JK, S.-B., JE, H., RH, E., Ehrlich, J., Garg, S., RF, H., and Rewers, M. (2003). Effect of type 1 diabetes on the gender difference in coronary artery calcification: a role for insulin resistance? The Coronary Artery Calcification in Type 1 Diabetes (CACTI) Study. Diabetes *52*, 2833–2839 7p.

Gorski, M., van der Most, P.J., Teumer, A., Chu, A.Y., Li, M., Mijatovic, V., Nolte, I.M., Cocca, M., Taliun, D., Gomez, F., et al. (2017). 1000 Genomes-based meta-analysis identifies 10 novel loci for kidney function. Sci. Rep. *7*, 45040.

Hadjadj, S., Péan, F., Gallois, Y., Passa, P., Aubert, R., Weekers, L., Rigalleau, V., Bauduceau, B., Bekherraz, A., Roussel, R., et al. (2004). Different Patterns of Insulin Resistance in Relatives of Type 1 Diabetic Patients With Retinopathy or Nephropathy. Diabetes Care *27*, 2661–2668.

Iyengar, S.K., Sedor, J.R., Freedman, B.I., Kao, W.H.L., Kretzler, M., Keller, B.J., Abboud, H.E., Adler, S.G., Best, L.G., Bowden, D.W., et al. (2015). Genome-Wide Association and Trans-ethnic Meta-Analysis for Advanced Diabetic Kidney Disease: Family Investigation of Nephropathy and Diabetes (FIND). PLoS Genet. *11*, 1–19.

Klein, R., Klein, B.E., Moss, S.E., and Cruickshanks, K.J. (1998). The Wisconsin Epidemiologic Study of Diabetic Retinopathy: XVII. The 14-year incidence and progression of diabetic retinopathy and associated risk factors in type 1 diabetes. Ophtha *105*, 1801–1815.

Krolewski, A.S. (2015). Progressive renal decline: The new paradigm of diabetic nephropathy in type 1 diabetes. Diabetes Care *38*, 954–962.

Ma, J., Möllsten, A., Prázny, M., Falhammar, H., Brismar, K., Dahlquist, G., Efendic, S., and Gu, H.F. (2006). Genetic influences of the intercellular adhesion molecule 1 (ICAM-1) gene polymorphisms in development of Type 1 diabetes and diabetic nephropathy. Diabet. Med. *23*, 1093–1099.

Marre, M., Jeunemaitre, X., Gallois, Y., Rodier, M., Chatellier, G., Sert, C., Dusselier, L., Kahal, Z., Chaillous, L., Halimi, S., et al. (1997). Contribution of genetic polymorphism in the renin-angiotensin system to the development of renal complications in insulin-dependent diabetes: Genetique de la Nephropathie Diabetique (GENEDIAB) study group. J. Clin. Invest. *99*, 1585–1595.

McKnight, A.J., Patterson, C.C., Sandholm, N., Kilner, J., Buckham, T.A., Parkkonen, M., Forsblom, C., Sadlier, D.M., Groop, P.H., Maxwell, A.P., et al. (2010). Genetic polymorphisms in nitric oxide synthase 3 gene and implications for kidney disease: a meta-analysis. Am. J. Nephrol. *32*, 476–481.

Nathan, D.M. (2014). The diabetes control and complications trial/epidemiology of diabetes interventions and complications study at 30 years: Overview. Diabetes Care *37*, 9–16.

Orchard, T.J., Dorman, J.S., Maser, R.E., Becker, D.J., Drash, A.L., Ellis, D., LaPorte, R.E., and Kuller, L.H. (1990). Prevalence of complications in IDDM by sex and duration. Pittsburgh Epidemiology of Diabetes Complications Study II. Diabetes *39*, 1116–1124.

Pattaro, C., Teumer, A., Gorski, M., Chu, A.Y., Li, M., Mijatovic, V., Garnaas, M., Tin, A., Sorice, R., Li, Y., et al. (2016). Genetic associations at 53 loci highlight cell types and biological pathways relevant for kidney function. Nat. Commun. *7*, 1–19.

Pezzolesi, M.G., Poznik, G.D., Mychaleckyj, J.C., Paterson, A.D., Barati, M.T., Klein, J.B., Ng, D.P.K., Placha, G., Canani, L.H., Bochenski, J., et al. (2009). Genome-Wide Association Scan for Diabetic Nephropathy Susceptibility Genes in Type 1 Diabetes. *58*,

1403–1410.

Pop, A., Clenciu, D., Anghel, M., Radu, S., Socea, B., Mota, E., Mota, M., Panduru, N.M., and RomDiane Study Group (2016). Insulin resistance is associated with all chronic complications in type 1 diabetes. J. Diabetes *8*, 220–228.

Rossing, P., Hougaard, P., and Parving, H.-H. (2002). Risk Factors for Development of Incipient and Overt Diabetic Nephropathy in Type 1 Diabetic Patients. Diabetes Care *25*, 859–864.

Sandholm, N., Salem, R.M., McKnight, A.J., Brennan, E.P., Forsblom, C., Isakova, T., McKay, G.J., Williams, W.W., Sadlier, D.M., Mäkinen, V.P., et al. (2012). New Susceptibility Loci Associated with Kidney Disease in Type 1 Diabetes. PLoS Genet. *8*, 1–13.

Sandholm, N., McKnight, A.J., Salem, R.M., Brennan, E.P., Forsblom, C., Harjutsalo, V., Makinen, V.-P., McKay, G.J., Sadlier, D.M., Williams, W.W., et al. (2013). Chromosome 2q31.1 Associates with ESRD in Women with Type 1 Diabetes. J. Am. Soc. Nephrol. *24*, 1537–1543.

Sandholm, N., Forsblom, C., Mäkinen, V.-P., McKnight, A.J., Österholm, A.-M., He, B., Harjutsalo, V., Lithovius, R., Gordin, D., Parkkonen, M., et al. (2014). Genome-wide association study of urinary albumin excretion rate in patients with type 1 diabetes. Diabetologia *57*, 1143–1153.

Sviklāne, L., Olmane, E., Dzērve, Z., Kupčs, K., Pīrāgs, V., and Sokolovska, J. (2018). Fatty liver index and hepatic steatosis index for prediction of non-alcoholic fatty liver disease in type 1 diabetes. J. Gastroenterol. Hepatol. *33*, 270–276.

The Diabetes Control and Complications Trial Research Group; (1995). Implementation of treatment protocols in the diabetes control and complications trial. Diabetes Care *18*, 361–376.

Thorn, L.M., Forsblom, C., Fagerudd, J., Thomas, M.C., Pettersson-Fernholm, K., Saraheimo, M., Wadén, J., Rönnback, M., Rosengård-Bärlund, M., Björkesten, C.-G.A., et al. (2005). Metabolic syndrome in type 1 diabetes: association with diabetic nephropathy and glycemic control (the FinnDiane study). Diabetes Care *28*, 2019–2024.

Wagener, D.K., Sacks, J.M., LaPorte, R.E., and MacGregor, J.M. (1982). The Pittsburgh study of insulin-dependent diabetes mellitus. Risk for diabetes among relatives of IDDM. Diabetes *31*, 136–144.

Van Zuydam, N.R., Ahlqvist, E., Sandholm, N., Deshmukh, H., Rayner, N.W., Abdalla, M., Ladenvall, C., Ziemek, D., Fauman, E., Robertson, N.R., et al. (2018). A genome-wide association study of diabetic kidney disease in subjects with type 2 diabetes. Diabetes *67*, 1–84.
